# Molecular basis for two stereoselective Diels-Alderases that produce decalin skeletons

**DOI:** 10.1101/2021.02.01.429105

**Authors:** Keisuke Fujiyama, Naoki Kato, Suyong Re, Kiyomi Kinugasa, Kohei Watanabe, Ryo Takita, Toshihiko Nogawa, Tomoya Hino, Hiroyuki Osada, Yuji Sugita, Shunji Takahashi, Shingo Nagano

**Affiliations:** Department of Chemistry and Biotechnology, Graduate School of Engineering, Tottori University, Tottori, Japan; Natural Product Biosynthesis Research Unit, RIKEN Center for Sustainable Research Science, Wako, Japan; Faculty of Agriculture, Setsunan University, Hirakata, Japan; Laboratory for Biomolecular Function Simulation, RIKEN Center for Biosystems Dynamics Research, Kobe, Japan; Center for Drug Design Research, National Institutes of Biomedical Innovation, Health, and Nutrition, Ibaraki, Japan; Graduate School of Pharmaceutical Sciences, The University of Tokyo, Tokyo, Japan; Chemical Biology Research Group, RIKEN Center for Sustainable Research Science, Wako, Japan; Theoretical Molecular Science Laboratory, RIKEN Cluster for Pioneering Research, Wako, Japan; Computational Biophysics Research Team, RIKEN Center for Computational Science, Kobe, Japan; Centre for Research on Green Sustainable Chemistry, Tottori University, Tottori, Japan

## Abstract

Molecular chirality, discovered by Louis Pasteur in the middle of the 19th century^1^, is found in most primary and secondary metabolites. Particularly, the so-called natural products are rich in chiral centres^2^. The stereochemistry of natural products is strictly recognized in living organisms, and is thus closely related to their biological functions. The Diels–Alder (DA) reaction, which forms a six-membered ring with up to four chiral centres, is a fundamental practical reaction for C–C bond formation in synthetic chemistry^3^. Nature has also adopted this reaction to elaborate the complex structures of natural products using enzymes derived from various progenitor proteins^4-7^. Although enzymes catalysing the DA reaction, Diels–Alderases (DAases), have attracted increasing attention, little is known about the molecular mechanism by which they control the stereochemistry and perform catalysis. Here, we solved the X-ray crystal structures of a pair of decalin synthases, Fsa2 and Phm7, that catalyse intramolecular DA reactions to form enantiomeric decalin scaffolds during biosynthesis of the HIV-1 integrase inhibitor equisetin and its stereochemical opposite, phomasetin^8,9^. Based on the crystal structures, docking simulations followed by all-atom molecular dynamics simulations provided dynamic binding models demonstrating the folding of linear polyenoyl tetramic acid substrates in the binding pocket of these enzymes, explaining the stereoselectivity in the construction of decalin scaffolds. Site-directed mutagenesis studies verified the binding models and, in combination with density functional theory calculations, clarified how hydrophilic amino acid residues in the Phm7 pocket regulate and catalyse the stereoselective DA reaction. This study highlights the distinct molecular mechanisms of the enzymatic DA reaction and its stereoselectivity *experimentally* and *computationally*. We anticipate that clarified molecular mechanism herein provides not only the basic understanding how these important enzymes work but also the guiding principle to create artificial enzymes that produce designer bioactive molecules.

## Introduction

Enzymes that create complex carbon frameworks with multiple chiral centres, such as polyketide synthase (PKS) and terpene cyclase, are gaining increasing attention not only in natural product chemistry but also in the chemical industry^10-13^. Enzymes catalysing the Diels–Alder (DA) reactions ([4+2] cycloadditions), the so-called Diels–Alderases (DAases), form two C–C bonds and up to four chiral centres to generate a cyclohexene from a conjugated diene and substituted alkene; these enzymes play key roles in controlling stereochemistry during the formation of polycyclic structures (Supplementary Fig. 1)^4-7^. Since the discovery of SpnF, the first monofunctional enzyme reported to catalyse a DA reaction^14^, many DAases have been identified in the biosynthetic pathways of bacterial, fungal, and plant origins. Unlike PKSs, nonribosomal peptide synthetases, and terpene cyclases, which share active site–containing domain structures involved in their specific functions^13,15^, DAases have no common structural features and are derived from distinct progenitor enzymes or proteins. SpnF, which catalyses DA reaction in the spinosyn A biosynthesis^14^, contains *S*-adenosylmethionine (SAM), and its overall structure belongs to the SAM-dependent methyltransferase family^16^, and the overall structures of PyrI4 and PyrE3, which are involved in the pyrroindomycin biosynthesis^17^, are very similar to those of lipocalin family proteins and FAD-dependent monooxygenases, respectively^18,19^. Although such DAases evolved independently from their own progenitors, they all exhibit high stereoselectivity and catalytic efficiency. The molecular basis of this emerging group of enzymes is gradually being revealed by genetic, biochemical and structural analyses, in combination with computational investigations^20-22^. However, the mechanisms underlying the remarkable features of naturally occurring DAases, such as the origin of stereoselectivity and their performanceas catalysts, have remained elusive.

Fsa2-family decalin synthases (DSs) found in filamentous fungi^8^ catalyse stereoselective DA reaction during the biosynthesis of decalin-containing pyrrolidin-2-ones (DPs), which exhibit various biological activities^23,24^; this class of molecules includes the HIV-1 integrase inhibitors equisetin (**1**) and phomasetin (**2**)^25,26^ and the telomerase inhibitor UCS1025A^27,28^ (Fig. 1a). Via intramolecular DA reaction, Fsa2 and its homolog Phm7 create a decalin scaffold with enantiomeric configurations from similar linear polyenoyl tetramic acids (e.g., **3** and **4**) (Fig. 1b)^8,9,29^. We have shown that replacement of *phm7* in a **2**-producing fungus with *fsa2* resulted in the production of a **1**-type decalin scaffold (2*S*,3*R*,8*S*,11*R*)^9^, indicating that these enzymes determine the stereochemistry of the decalin scaffold during DP biosynthesis. Another homologous enzyme, MycB^30^, also produces a **1**-type decalin scaffold, and the **2**-type decalin scaffold is produced by CghA^31^ and UcsH^32^, which are involved in the biosynthesis of Sch 210972 and UCS1025A, respectively. Interestingly, but not surprisingly, another decalin scaffold (2*R*,3*S*,8*S*,11*R*) is produced by PvhB, which is involved in the varicidin A biosynthesis^33^ (Fig. 1a). Thus, structural comparisons of the DSs that differ in function, i.e., stereochemical output, provide insight into the mechanisms of the stereoselective DA reactions.

**Fig. 1:**
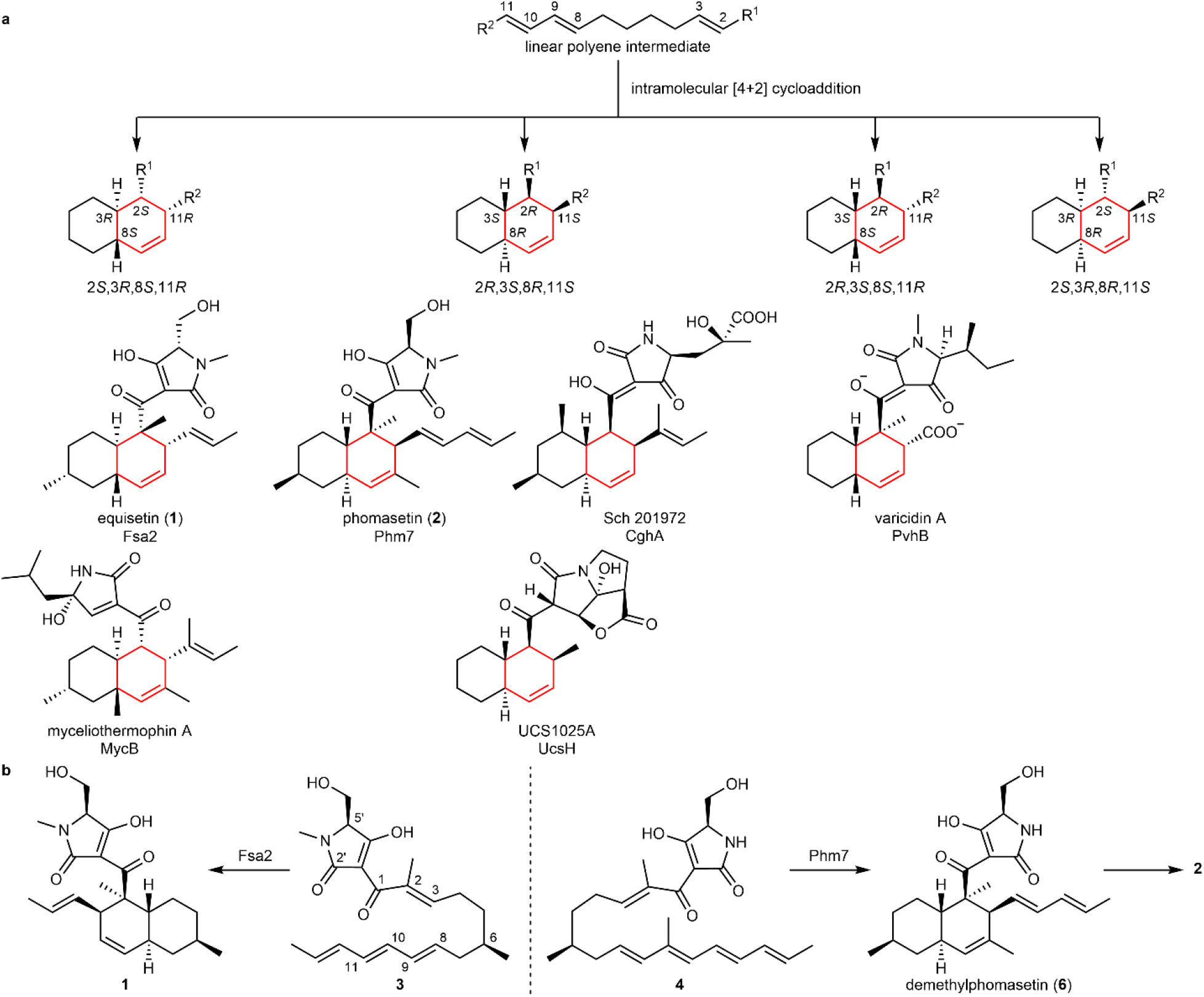
The Fsa2-family decalin synthases (DSs) catalysing stereoselective Diels-Alder (DA) reaction. **a**, Four possible configurations of decalin formed via intramolecular DA reaction, and corresponding Fsa2-family DSs. **b**, The reactions catalysed by Fsa2 and Phm7 to form enantiomeric decalin scaffolds. Linear tetraenoyl tetramic acid **3** was chemically synthesized and confirmed to be a Fsa2 substrate to form **1**^29^, whereas pentaenoyl tetramic acid **4** is a likely Phm7 substrate to yield *N*-demethylphomasetin (**6**), which is further converted to **2**.

In this study, we revealed the molecular basis of two stereoselective enzymes, Fsa2 and Phm7, that catalyse DA reaction to form enantiomeric decalin scaffolds. X-ray crystal structures of substrate-free Fsa2 and Phm7, and Phm7 bound to an inhibitor (hereafter referred to as inhibitor-bound Phm7), were determined at 2.17, 1.62, and 1.61-Å resolution, respectively. The substrate-bound poses were modelled using docking simulations followed by all-atom molecular dynamics (MD) simulations. We employed the generalized replica-exchange with solute tempering (gREST) method^34^, which allows extensive sampling of possible binding poses of the substrates^35,36^. Site-directed mutagenesis studies were performed to verify the binding models for stereoselective synthesis and examine the amino acid residues involved in substrate interactions. Density functional theory (DFT) calculations were performed to reveal the reaction mechanism in detail, particularly the stereoselectivity and rate acceleration by Phm7. This powerful combination of experimental methods and calculations, which we used to investigate two enzymes that produce enantiomeric decalin scaffolds, provides insight into the molecular mechanism of enzyme-mediated DA reaction and its stereoselectivity.

## Results

### Structure analyses of Fsa2, Phm7, and inhibitor-bound Phm7

The crystal structure of Fsa2, which produces the 2*S*,3*R*,8*S*,11*R* decalin scaffold found in equisetin (**1**, Fig. 1a, b, see Supplementary Fig. 2 for compound structures used in this study), has a β-sandwich and a β-barrel domain at the N- and C-termini, respectively (Fig. 2a). The structures of both domains exhibit structural similarity with those of lipocalin family proteins that bind heme, steroids, and other hydrophobic ligands in the pocket located in their β structures^37^. Interestingly, however, Fsa2 does not have a pocket in either the N- or C-domains (Supplementary Fig. 3). Instead, a large pocket is present between the two domains. Considering the likely substrate structure and volume, we speculate that the large cavity created by the two domains is an active- and substrate-binding site. Phm7 catalyses intramolecular DA reaction to produce the enantiomeric decalin scaffold (Fig. 1b). Despite their distinct stereoselectivity and low sequence similarity (36 % sequence identity), the crystal structure of Phm7 is similar to that of Fsa2 (RMSD = 0.849 Å), and the shape and volume of their large pockets are also similar (Fig. 2b).

**Fig. 2:**
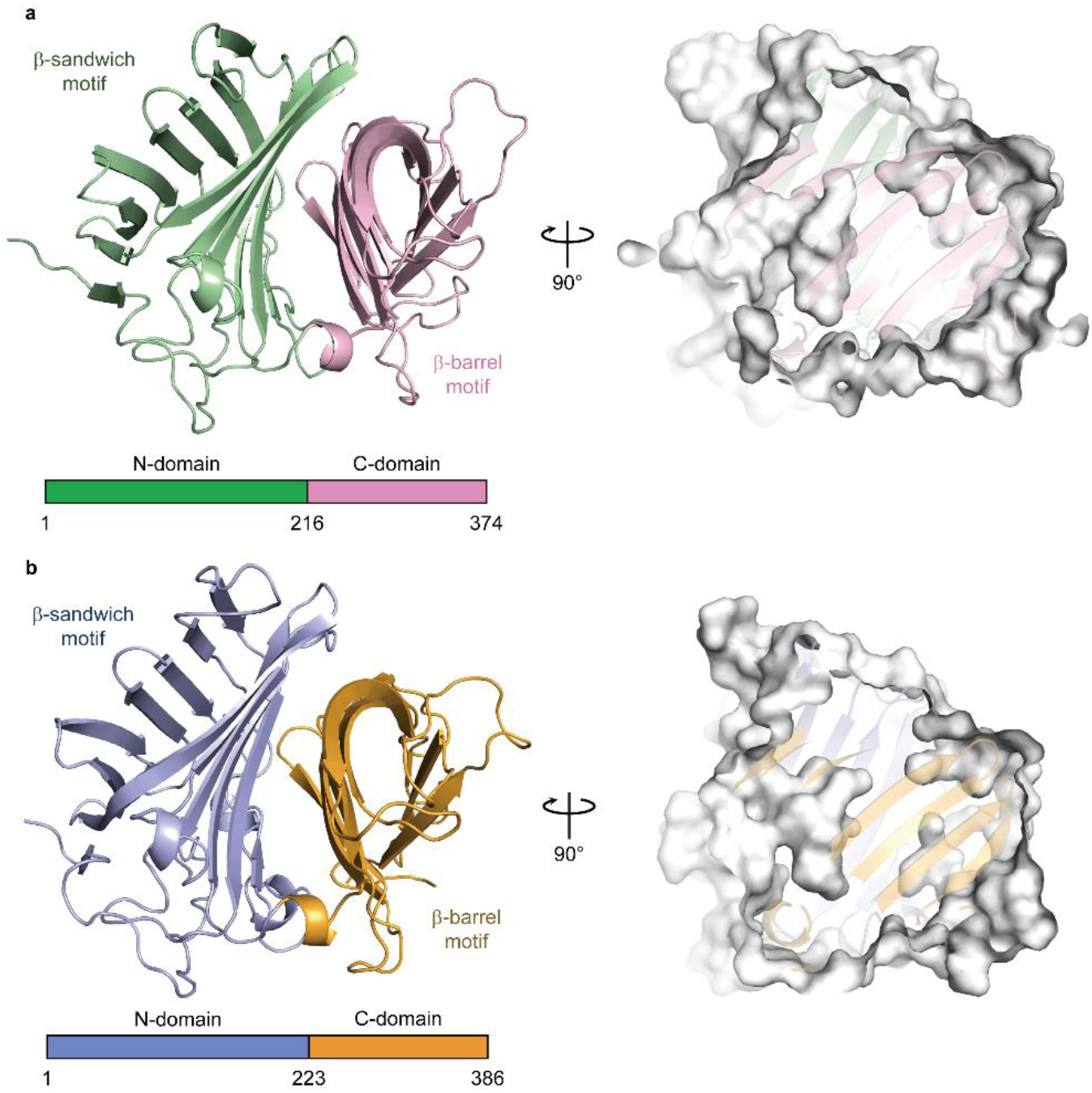
Crystal structures of Fsa2 and Phm7. **a**,**b**, Overall structures of Fsa2 (**a**) and Phm7 (**b**). Both DSs consist of two domains: N-domains for residues 1–215 of Fsa2 and 1–222 of Phm7, and C-domains for residues 216–374 and 223–386. Surface models (right panels) show shapes of the pocket between the two domains of both enzymes.

To probe the active site of the DSs, we performed Phm7 ligand screening using microscale thermophoresis (MST)^38^ (Supplementary Fig. 4). We found that 3-aminomethyl-*p*-menthane (**5**), which was similar to the phomasetin (**2**) substructure, dose-dependently inhibited Phm7 activity *in vitro* and production of **2** in the fungus *Pyrenochaetopsis* sp. RK10-F058 (Supplementary Fig. 5). Compound **5** also inhibited equisetin (**1**) production in the fungus *Fusarium* sp. FN080326 (Supplementary Fig. 6). We next performed co-crystallization of Phm7 and **5** and obtained the crystal structure of the inhibitor-bound Phm7 at 1.61-Å resolution (Supplementary Fig. 7). The inhibitor was located on the lower side of the pocket and formed hydrophobic interactions with Y178, W223, F226, L245, and L381. The amino group of **5** was surrounded by Y68, E51, and Y178. Because **5** can bind to both Phm7 and Fsa2 to inhibit their functions (Supplementary Figs. 4–6), the lower side of the pocket between the two domains would form the active site of these enzymes. Indeed, CghA, exhibiting the same stereoselectivity as Phm7 (Fig.1a)^31^, was recently reported to have two β domains, as in Phm7 and Fsa2^39^. The product binding site is almost the same as the **5**-binding site of Phm7.

### Docking and MD simulations for binding pose determination

To determine whether the pocket would be large enough to accommodate the substrates, substrate **4** was docked into the crystal structure of the inhibitor-bound form of Phm7 using AutoDock Vina. Substrate **4** fits within the pocket in various binding poses, including a folded form in which a U-shaped folded alkyl chain including asymmetric C6 of **4** was located at the lower side of the pocket where **5** can bind, and the tetramic acid and polyene moieties were extended into the inner upper part of the pocket (Supplementary Fig. 8a). Likewise, folded substrate **3** fits within the pocket of Fsa2 (Supplementary Fig. 8b). These docking simulations using the crystal structures indicated that the pocket between the N- and C-domains of Phm7 and Fsa2 has enough room for binding of the folded substrates.

To explore the binding poses of the substrates and their dynamics in the pockets, we carried out all-atom MD simulations using the gREST method^34^, which extensively samples possible binding poses otherwise elusive in conventional simulations. A variety of poses were obtained (see Supplementary Movies S1 and S2 for collected binding poses of Phm7 and Fsa2, respectively), and clustering analysis of the simulation trajectories resulted in four major bound poses, including “folded” and “extended” conformations, for both **4** and **3** (Supplementary Fig. 9). In Fig. 3a, the major cluster of Phm7 (63% of Phm7, pA in Supplementary Fig. 9a) shows a well-defined bound pose for **4**. In this cluster, the tetramic acid moiety and polyene tail of the folded poses were located at the upper front and back of the pocket, respectively, whereas the U-shaped part was found at the lower side of the pocket. Similar tetramic acid–front and polyene–back poses were also found in the folded conformations in Fsa2 (33% of Fsa2, fA in Supplementary Fig. 9b), although bound **3** fluctuated in the pocket to a greater extent than in Phm7 (Fig. 3b). In both Phm7 and Fsa2, the electrostatic potential inside the pocket had a large negative value as it moved deeper into the pocket (Supplementary Fig. 10). The bottom surface of the pocket was significantly hydrophobic, whereas the hydrophilic surface was found in the upper wall of the pocket (Supplementary Fig. 11). These inhomogeneous electrostatic, hydrophobic, and hydrophilic environments of the pocket coincide with the common tetramic acid–front and polyene–back orientation.

**Fig. 3:**
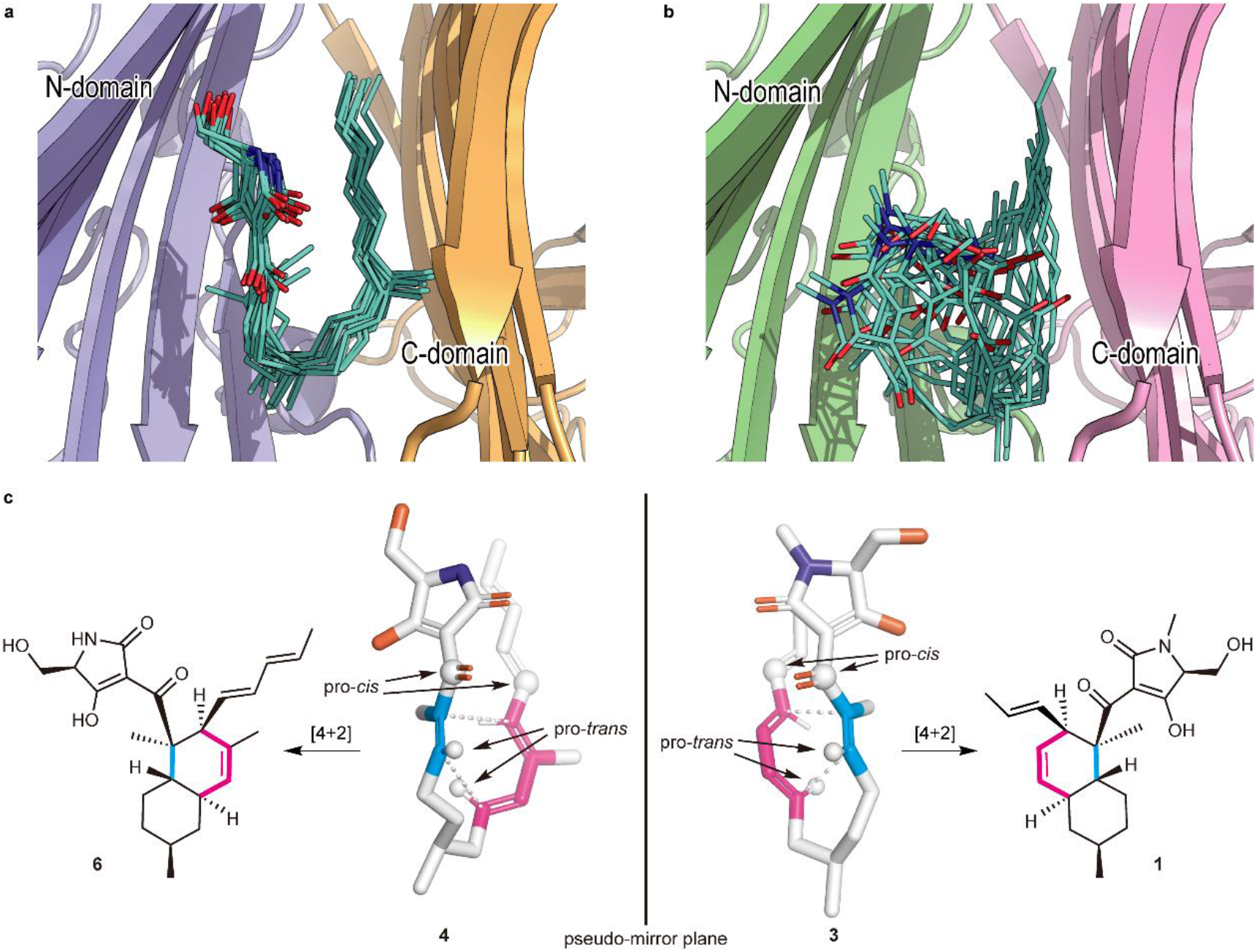
Predicted binding model. **a, b**, A collective view of 12 representative snapshots taken from the top 70% of the main folded clusters (chosen based on the RMSD values from the cluster centre in ascending order) for Phm7 (**a**) and Fsa2 (**b**), respectively. **c**, Conformations of the representative poses of **4** (left) and **3** (right) in the pockets. The diene and dienophile moieties of the substrate are indicated in pink and cyan, respectively. The corresponding cycloadducts are also shown.

The representative folded conformation of the major cluster of Phm7, which dominates the cluster (Supplementary Fig. 12a), explains the configuration of the stereoselective DA reaction product with the 2*R*,3*S*,8*R*,11*S* decalin scaffold (Fig. 3c). In contrast to Phm7, the pocket of Fsa2 allows various “folded” conformations of **3** (Supplementary Fig. 12b). Nevertheless, a certain number of poses are consistent with the configuration of the **1**-type (2*S*,3*R*,8*S*,11*R*) decalin scaffold (Fig. 3c). Comparison of the carbon chain conformation (C– C–C–C dihedral angles) between substrates **3** and **4** revealed that the diene and dienophile moieties exhibit pseudo-enantiomeric conformations, consistent with the stereochemical relationship of the transition state structures corresponding to decalin scaffolds of **1** and **2** (Supplementary Fig. 13). The LigPlot analysis of the representative pose of **4** shows that the tetramic acid moiety forms hydrogen bonds with the side chains of E51, N84, and K356, and with the main chain carbonyl of G64. The U-shaped part and polyene tail of the substrate fit with the hydrophobic region of the pocket lined by L49, W223, Y232, L245, F341, W342, L381, and I383 (Supplementary Fig. 14a). Similarly, the hydrophobic U-shaped part and polyene tail of **3** are surrounded by many hydrophobic side chains of Fsa2 (W45, V169, Y171, W216, Y225, and M238), and the tetramic acid moiety is hydrogen-bonded with N346 (Supplementary Fig. 14b). Therefore, MD simulations based on the crystal structures of the two enzymes suggested distinct substrate–enzyme interactions and their resultant substrate poses corresponding to the enantiomeric decalin scaffolds.

### Functional analyses of Phm7 and Fsa2

To examine the substrate–enzyme interactions predicted from the MD simulations, we first established an *in vitro* enzyme assay system using cell lysates prepared from fungal mycelia lacking the *phm7* gene^9^. The cell lysates were directly incubated with Phm7 or Fsa2, and their reaction products were analysed by liquid chromatography/electrospray ionization mass spectrometry (LC/ESI-MS). Linear polyenoyl tetramic acid **4** was less abundant in the presence of the enzymes, and the expected products of Phm7 and Fsa2, *N*-demethylphomasetin (**6**) and its derivative containing **1**-type *trans*-decalin (**7**), respectively, were formed (Fig. 4a, b). The reaction selectivity of Phm7 and Fsa2 shown in the *in vitro* assay was consistent with that observed in the producer fungus and a mutant in which *phm7* was replaced with *fsa2*^9^. Furthermore, we observed no significant difference in the amount of *cis*-decalin containing derivative **8** between the presence and absence of enzyme. The time course of the *in vitro* reaction confirmed the linear formation of product **6** for the first several minutes, and no cycloaddition in the absence of enzyme under the conditions tested (Supplementary Fig. 15). These results indicated that products **6** and **7** were exclusively formed by the action of Phm7 and Fsa2, respectively, from the linear polyene substrate found in the fungal cell lysate. Therefore, the *in vitro* assay using the fungal cell lysate allowed us to evaluate the enzyme activities by measuring the products formed.

**Fig. 4:**
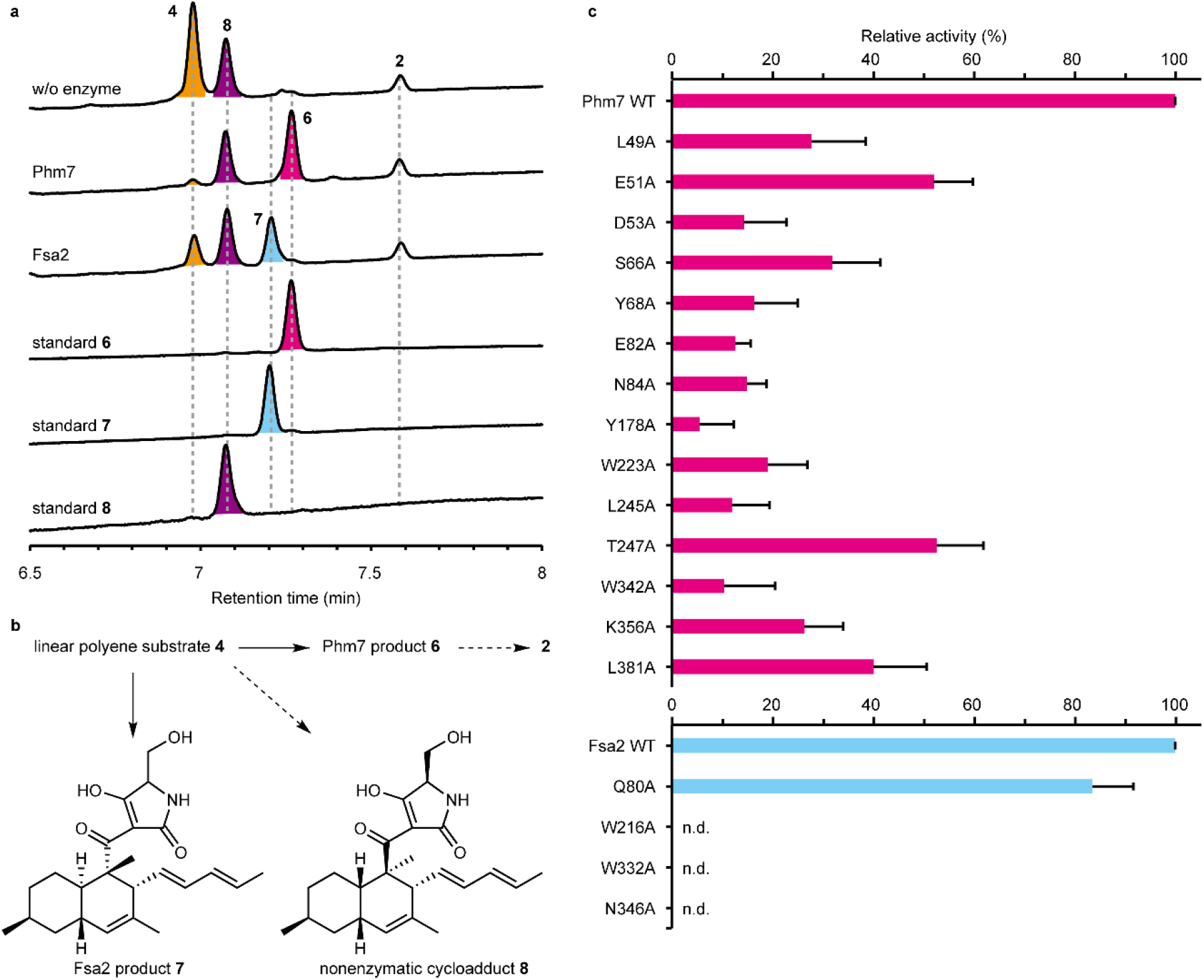
*In vitro* analysis of Ala-substituted Phm7 and Fsa2 mutants. **a**, UPLC traces of the *in vitro* Phm7 and Fsa2 reaction products. The cell lysates prepared from the Δ*phm7* mycelia were incubated with Phm7 and Fsa2 and analysed by LC/ESI-MS. **b**, Linear polyenoyl tetramic acid **4** was converted into **6** and **7** by Phm7 and Fsa2, respectively. Conversions indicated by broken arrows were not detected under the conditions tested: non-enzymatic formation of **8** from **4** was not observed *in vitro*, and sinefungin added in the reaction buffer inhibited *N*-methylation of **6. c**, Comparison of the enzyme activities of the Ala-substituted mutants with those of the wild-type Phm7 and Fsa2. Data are the mean and error bars represent the standard deviation of four independent experiments. n.d., not detected.

Using the *in vitro* assay system, we carried out site-directed mutagenesis studies of Phm7 and Fsa2 to validate the substrate binding modes predicted by the MD simulations (Supplementary Figs. 14, 16). The key amino acid residues that interact with the tetramic acid moieties (K356 in Phm7 and N346 in Fsa2), the hydrophobic U-shaped parts (W223 in Phm7 and W216 in Fsa2), and the polyene tails (W342 in Phm7 and W332 in Fsa2) were substituted with Ala, and their enzyme activities were compared with those of the wild-type enzymes. Ala substitutions of these amino acid residues significantly decreased enzyme activity *in vitro* (Fig. 4c), indicating that the pockets between the N-and C-domains of Phm7 and Fsa2 were the active sites of both enzymes, and that the proposed binding modes were reliable.

Next, we focused on Phm7 to demonstrate the binding modes. We introduced additional single Ala substitutions into the amino acid residues close to substrate **4** (Supplementary Fig. 16a,b), and performed the enzyme assay. Ala substitutions of the residues in proximity to the tetramic acid moiety of **4** (Supplementary Fig. 17) significantly decreased the enzyme activity (Fig. 4c). In particular, we observed a marked decrease in the activity of mutants harbouring substitutions in hydrophilic residues such as D53 and E82, suggesting that the hydrophilic upper wall of the pocket (Supplementary Fig. 10) was indeed responsible for trapping the tetramic acid moiety of **4**, as predicted by the MD simulations. We also expected that the hydrogen bond donations from K356 to the carbonyl oxygen at C1, as well as those of E51 and N84 to the tetramic acid moiety, would activate the dienophile to accelerate the reaction. Ala substitution at Y178 and W223 also significantly decreased enzyme activity (Fig. 4c). A230, L245, and T247 are located near the substrate but do not interact directly with the substrate in the MD model. Substitution of these residues with smaller ones (L245V and T247A) yielded mutants that retained considerable activity, whereas substitution with longer and larger ones (A230F and T247F) significantly decreased activity, probably due to steric clash (Fig. 4c and Supplementary Fig. 17). These results confirmed that the **5**-binding site formed by amino acid restudies such as Y178, W223, L245, and T247 is the reaction chamber in which the stereospecific DA reaction occurred.

To further validate the substrate–enzyme interactions in Phm7, we examined the effects of Ala substitutions of Phm7 on phomasetin (**2**) production in the producer fungus, in which the wild-type *phm7* was replaced with the mutant genes. Phm7 Y68A and W223A mutants produced significantly lower levels of **2** and suppressed production of **8** in the fungus (Supplementary Fig. 18). In addition, a new peak **9** was detected in the culture extract of the W342A mutant. Structural analyses by NMR and MS revealed that **9** was a derivative of **2** with a hydroxy group at the terminus of polyene (See Supplementary Note for structure determination of **9**, Supplementary Fig. 18). Production of **9** in this gain-of-function mutant supported the role of W342 in the interaction with the polyene tail of **4**. Taken together, our in-depth analyses of Phm7 mutants using an *in vitro* enzyme assay and *in vivo* production in fungus demonstrated that the MD simulation–based binding models were reliable, and that the predicted amino acid residues are involved in substrate binding.

### DFT calculations for the molecular mechanism in the Phm7 pocket

Given that Phm7 promotes the intramolecular DA reaction of substrate **4** in a stereoselective manner, the detailed molecular mechanism in this event with Phm7 is of great importance. Hence, we focused on how Phm7 determined the stereoselectivity of the DA reaction, as well as whether this enzyme can accelerate (i.e., catalyse) the reaction. MD simulations pointed out that the structure of **4** seemed to be convergent in the major folded conformations in the pocket of Phm7 (Supplementary Fig. 12a). The geometry between the dienophile and diene moieties in this “folded” structure is close to the configuration that would afford the decalin scaffold with the same configuration as Phm7 product **6** (Fig. 3c). We investigated the intrinsic stereoselectivity of the uncatalysed DA reaction of **4** using DFT calculations at the M06-2X/6–311+G** (scrf = CPCM, water) level of theory (Fig. 5a). Among the four transition states to the corresponding decalin derivatives, **TS**_***1a***_, which affords **8**, is located in the lowest energy state (Δ*G*^‡^ +16.1 kcal/mol) relative to the linear conformation, whereas **TS**_***1b***_, which affords **6**, requires slightly higher activation energy (Δ*G*^‡^ +16.5 kcal/mol). The reactions that afford Fsa2-type product **7** and another *cis*-decalin derivative were found to be less feasible via **TS**_***1c***_and **TS**_***1d***_, respectively. These computational results suggested that the DA reaction without any steric bias should give a mixture of **8** and **6**, whereas **7** might also be present as a minor component. Thus, the combined experimental and theoretical results indicated that efficient folding of the substrate in the Phm7 pocket plays a pivotal role as a major determinant of stereoselectivity.

**Fig. 5:**
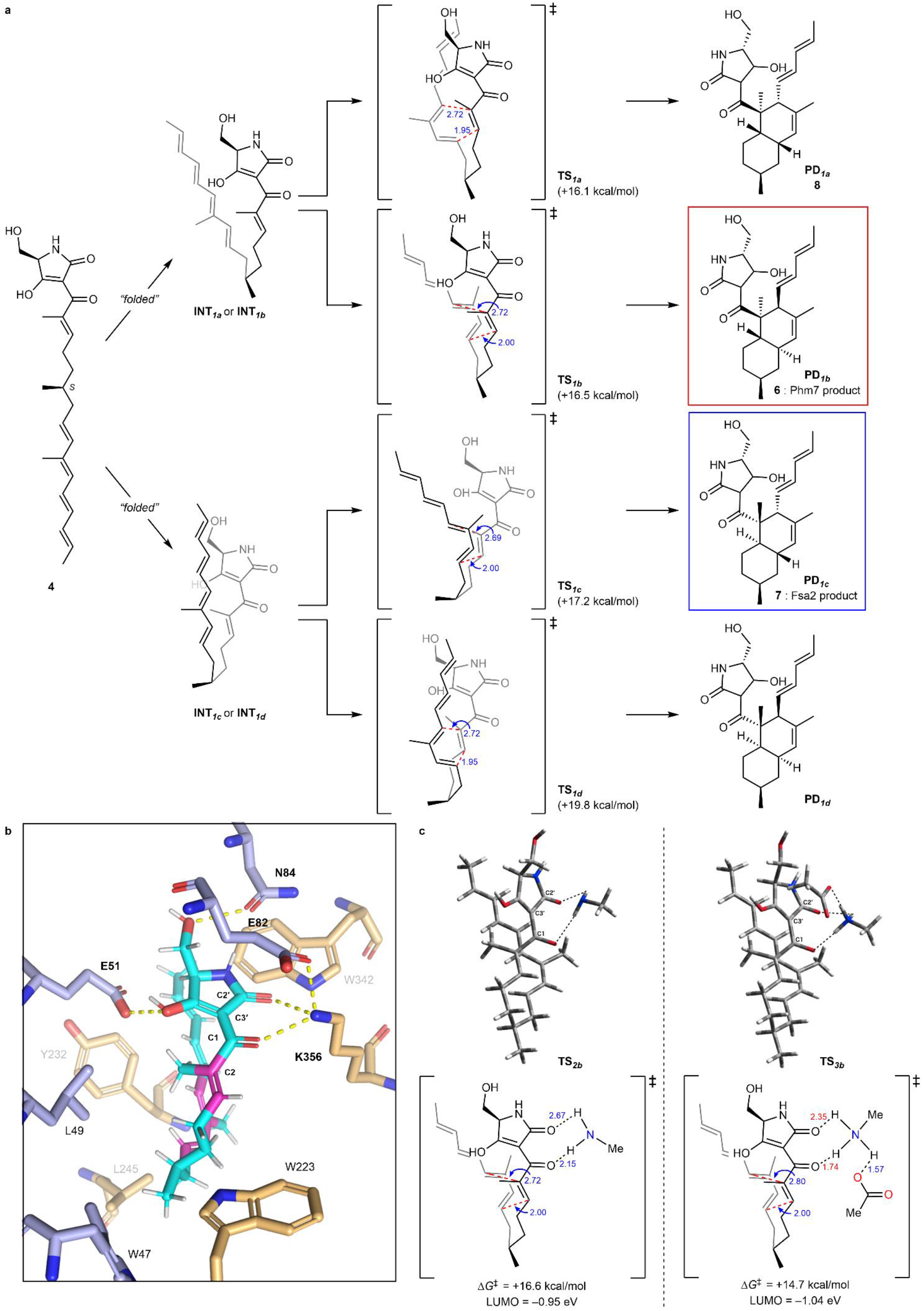
The DFT calculations for the reaction mechanism of the uncatalyzed and hydrogen bond-catalysed DA reaction of 4. **a**, Reaction pathways for the uncatalyzed DA reaction of substrate **4** to produce four types of decalin stereoisomers. The energy changes and bond lengths were calculated at the M06-2X/6-311+G** (scrf = CPCM, water) level of theory are shown in kcal/mol and Å, respectively. **b**, Illustration of the hydrogen bond network of **4** with amino acid residues in the Phm7 pocket obtained from the MD simulation (cf. Supplementary Fig. 14a). **c**, Transition state structures to give **6** with methylamine (**TS**_***2b***_) and both methylamine and acetic acid (**TS**_***3b***_) as models of the hydrogen bonding with K356 and E82 residues.

Next, to gain some insight into the rate acceleration mechanism, we investigated the amino acid residues that interact with **4** in the Phm7 pocket. In particular, we focused on the polar amino acid residues that trap the tetramic acid moiety in the upper part of the pocket. Similar to Lewis acid coordination, efficient hydrogen bond donation to the carbonyl group adjacent to the dienophile should lower the lowest unoccupied molecular orbital (LUMO) energy of the dienophile and facilitate the DA reaction^40^. From this point of view, the amino moiety of K356 is a hydrogen bond donor to the corresponding carbonyl group at the C1 of folded **4** (Fig. 5b). DFT calculations at the same level of theory were performed to track the course of the DA reaction of **4** with methylamine as a model of the K356 residue. However, no significant acceleration was observed, and the activation barrier for **TS**_***2b***_was +16.6 kcal/mol relative to the folded conformation. In fact, no efficient decrease in LUMO energy was observed (−0.90 eV for **TS**_***1b***_vs −0.95 eV for **TS**_***2b***_, Fig. 5a, c). Taking a closer look at the binding model, we discerned that the amino moiety of K356 makes another hydrogen bond with the carboxy group of E82 to form an ammonium architecture (Fig. 5b). We carried out DFT calculations by incorporating methylamine and acetic acid as models to reproduce the hydrogen bond network (i.e., **4**-K356-E82)^22^. The modelled network provides a tight hydrogen bond on the carbonyl oxygen at the C1 of **4** (1.74 Å) due to the acidified proton of the amino group. Furthermore, because the amino protons became more electron-deficient, an additional weak hydrogen bond^40^ (2.35 Å) formed with the carbonyl oxygen at C2′. Therefore, the dihedral angle of the C1 and C2′ carbonyl groups in **TS**_***3b***_(25.4°) became much smaller than that of **TS**_***2b***_(33.9°), making the conjugation between the enone dienophile and tetramic acid moiety more efficient. Thus, both electronic and structural perturbation by the hydrogen bond network decreased the LUMO energy to −1.04 eV in **TS**_***3b***_, and the DA reaction was greatly facilitated by lowering the activation barrier for **TS**_***3b***_. (+14.7 kcal/mol). These results explain the marked decrease in the enzymatic activity of the E82A mutant (Fig. 4c). In addition, the direct incorporation of the carboxy group of E82 in hydrogen bonding with **4** in the absence of K356, which was also reproduced by the DFT calculation model (Supplementary Fig. 19), could explain why the Ala substitution of K356 did not cause a complete loss of activity. Computation revealed that other hydrogen bonds between the tetramic acid moiety and the amino acids, such as E51 and N84, did not lower the LUMO energy level (Supplementary Fig. 20), and should be devoted to the appropriate positioning of substrate **4** in the Phm7 pocket.

In sum, the triad of experiments, MD simulations, and DFT calculations revealed the molecular mechanism of the Phm7 pocket as the catalyst for stereoselective DA reaction. Key findings are as follows: (i) although the intrinsic selectivity of the DA reaction of **4** prefers the formation of **8** in the absence of the enzyme, the folding structure in the pocket defines the selective formation of **6**; (ii) the sophisticated hydrogen bond network derived from K356 and E82, namely, Brønsted acid–activated hydrogen bonding catalysis, promotes the DA reaction event very efficiently; and (iii) some polar amino acid residues guide the substrate to the opportune folding structure.

## Discussion

Since the first report of direct evidence in biological DA reaction in solanapyrones biosynthesis and crystal structure of macrophomate synthase^41,42^, many DAases have been discovered, and their crystal structures have been determined.^16,18,19,21,43-47^. Their structures tell us that these enzymes were derived from ones with distinct functions other than catalysing DA reactions, implying that their active sites were subsequently converted into those for the DA reaction. By sharp contrast, DSs do not use the ligand-binding site of lipocalin family protein as an active site for the DA reaction. Fusion of two lipocalin-family proteins generated two-β-domain proteins with a pocket between the two domains. By burying their pockets in the β-domains, these proteins then acquired the ability to bind unstable linear polyenoyl tetramic acids using the relatively relaxed pocket, and eventually catalyse the DA reaction. A protein from *Nitrosomonas europaea*, NE1406, has a tertiary structure very similar to those of DSs^48^. This protein, which consists of two lipocalin-like domains and has several small pockets in and between the two β-domains (Supplementary Fig. 21), is likely to be involved in the carotenoid biosynthesis. From the standpoint of molecular evolution, nature has utilized the fusion (or duplication) of lipocalin proteins to generate new functions in a pocket between two domains.

Several amino acid residues are conserved among the DSs whose functions have been characterized (Supplementary Fig. 22). Many of them are located inside the molecule and likely play roles in maintaining enzyme structure rather than in catalysis per se. Among the amino acid residues lining the active site pocket, only Trp at the bottom of the pocket (W223 in Phm7 and W216 in Fsa2) is highly conserved, suggesting that it plays a key role in the DA reaction. Indeed, MD simulations predicted that the W223 and W216 form hydrophobic interactions with substrates **4** and **3**, respectively (Supplementary Fig. 14c). The substitutions of the conserved Trp with Ala substantially retarded the enzyme activity for both Phm7 and Fsa2. Although the sequence homology among the DSs is limited, they seem to share amino acid distributions in the pockets between the two domains (Supplementary Fig. 23). It is likely that substrates bind to the pocket in a similar way (tetramic acid–front and polyene–back orientation), allowing stereoselective DA reactions to proceed. Considering that the substrate conformation in the enzyme pocket is correlated to the product stereochemistry, we propose a model of how DSs produce decalin scaffolds with four possible configurations via the DA reaction (Fig. 6). The conformations of **TS**_***1b***_and **TS**_***1c***_are enantiomeric to each other, except for the C6 methyl group (Fig. 5a), and all the C–C–C–C dihedral angels of the carbon chains are opposite (Supplementary Fig. 13). This can be described as a difference in the manner of folding the linear substrate, i.e., the linear substrate **3** is folded clockwise and **4** counter-clockwise to yield (pseudo)enantiomeric conformations in the enzyme pocket (Fig. 3c). When the folded substrate **3** is flipped horizontally, it binds to the Fsa2 pocket in the tetramic acid– front and polyene–back orientation. A key difference between the transition state structures for *trans* and *cis*-decalin scaffolds is the rotation at the C7–C8 bond. Thus, the substrate conformation in the enzyme pocket can be predicted for *cis*-decalins, such as varicidin A (2*R*,3*S*,8*S*,11*R*, Fig. 1b) and even an unidentified metabolite (2*S*,3*R*,8*R*,11*S*). Recent phylogenetic analysis suggested that approximately 100 sequences are potentially involved in DP biosynthesis^49^. Therefore, this group of enzymes, which is widespread in Ascomycota fungi, are attractive targets for structure–function relationship studies, providing key structural features for determining the stereoselectivity of these reactions.

**Fig. 6:**
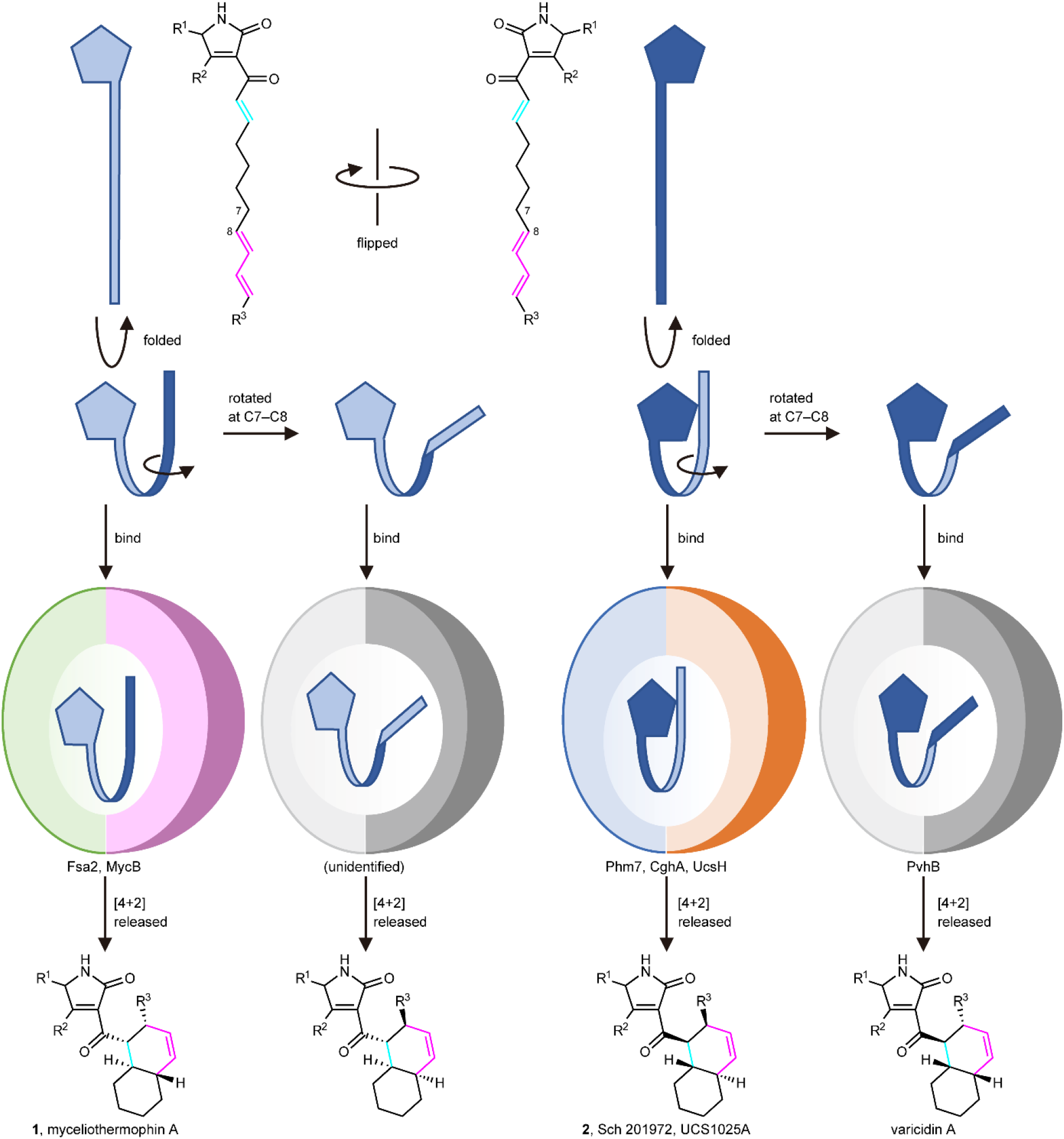
A model of the formation of four diastereomeric decalin scaffolds by decalin synthases. Linear polyenoyl tetramic acid substrates are shown schematically as a pentagon and a ribbon. Folded substrates bind to corresponding DSs in the tetramic acid–front and polyene–back manner, and are preorganized in the pocket. The substrates only with productive conformations are selected and undergo the reaction to produce respective decalin scaffolds.

The MD simulations showed that the pockets between the N- and C-domains of DSs were large enough to accommodate the substrates in various conformations. The “folded” cluster of substrate **4** contains a dominant pose (80%) with an *s*-*cis* conformation at C9–C10, explaining the expected decalin configurations (Supplementary Fig. 24). The calculated C– C–C–C dihedral angles along the carbon chain of the dominant pose takes a single orientation at all the rotational bonds, and no pose corresponding to the other possible decalin configurations was observed. This indicates that the Phm7 pocket robustly regulates the conformation of the substrate to stabilize the folded pose with a specific conformation. On the contrary, substrate **3** in the Fsa2 pocket exhibits a rather large variation in folded poses (Supplementary Fig. 12b). This is likely due to the fact that Fsa2 binds a smaller substrate (the volumes of **3** and **4** are 379 Å^3^ and 402 Å^3^, respectively) in a larger pocket relative to Phm7 (the pocket size is approximately 150 Å^3^ larger than that of Phm7, Supplementary Fig. 25). In addition, unlike Phm7, which tightly holds the tetramic acid moiety by multiple hydrogen bonds, there is no hydrophilic amino acid residue in proximity to the C1 carbonyl, and only N346 is involved in hydrogen bonding with **3**. Fsa2 likely retains the substrate affinity to the enzyme by accommodating various folded conformations and minimizing entropy loss upon binding. This enzyme still produces a reactive conformation for the DA reaction with a desired stereochemistry. Our preliminary analysis suggests that both of the hydrogen bonding of substrate **3** with N346 and the indole ring orientation of W216 play key roles in stabilizing the reactive conformation (Supplementary Fig. 26). Further investigation on the enzymatic reaction mechanism, including activation free energies, using a hybrid quantum mechanics (QM) / molecular mechanics (MM) method, should fill in the missing piece of the catalytic mechanisms. Given the structural similarity between Phm7 and Fsa2 and their opposite selectivity in the DA reaction, clarification of the detailed mechanism would be of great significance, and is the subject of ongoing research in our group.

In conclusion, our investigations combining experimental and theoretical approaches highlighted the distinct molecular mechanisms underlying Phm7- and Fsa2-catalysed DA reactions. The folding of substrate **4** in the Phm7 pocket and **3** in the Fsa2 pocket, predicted by the gREST method based on X-ray crystal structures, exhibited stereoselectivity in the construction of decalin scaffolds. The results of site-directed mutagenesis studies and DFT calculations clarified how the hydrophilic amino acid residues in the Phm7 pocket regulate and catalyse the stereoselective DA reaction. In addition, our results raise questions about the molecular tactics adopted by Fsa2, which should be addressed in future research: what determines the transition state from the flexible folded conformations in the pocket, and how dose Fsa2 accelerate the DA reaction without effective hydrogen bonding on the dienophile moiety?

## Methods

### Chemicals

All solvents and reagents were of analytical grade and purchased from commercial sources unless noted otherwise. The small molecules used in MST screening, including 3-aminomethyl-*p*-menthane (**5**), and sinefungin were purchased from Namiki Shoji and Cayman Chemical, respectively.

### Expression of Fsa2 and Phm7 in *Escherichia coli*

The ORFs of *fsa2* and *phm7* were amplified by PCR with the genomic DNA of *Fusarium* sp. FN080326^50^ and *Pyrenochaetopsis* sp. RK10-F058^51^, respectively. The amplified decalin synthase (DS) genes were cloned into the pGEM T-easy vector (Promega) and verified by sequencing. The gene fragments were excised by NdeI–XhoI digestion and ligated to the pET28b(+) vector (Novagen), resulting in pET28b(+)-*fsa2* and -*phm7*. Due to inefficient cleavage and removal of the His-tag at the N-terminus of Fsa2, modified pET28b(+)-*fsa2* lacking a thrombin site and linker peptide was used for crystallization study. The primer pairs used for the amplification are listed in Supplementary Table 1. PCR-based site-directed mutagenesis of the DS genes was performed using the primers listed in Supplementary Table 1. Whole plasmids of the mutant DSs were amplified using pET28b(+)-*fsa2* and -*phm7* as templates, which were digested with DpnI after PCR. The amplified cyclic DNAs were introduced into *E. coli* DH5α, and the mutated plasmids were verified by sequencing.

*E. coli* BL21 Star™ (DE3) (Invitrogen) cells were transformed with pET28b(+)-*fsa2* and *-phm7*, and then cultured in Terrific broth. Gene expression was induced by the addition of 0.5 mM isopropyl-β-d-thiogalactopyranoside, and further cultured at 30 °C for 6 h. For preparation of selenomethionine (SeMet)-substituted Phm7, *E. coli* B834(DE3) carrying pET28b(+)-*phm7* was cultured at 18 °C for 96 h using the Overnight Express Autoinduction System 2 (Novagen). Cells expressing the DS were harvested by centrifugation and frozen at –80 °C until use.

### Purification of Fsa2 and Phm7

The collected Fsa2 expressing cells were resuspended in buffer A (50 mM Tris-HCl, pH7.5, 200 mM NaCl, 10 % v/v glycerol) supplemented with 0.5 mg/mL lysozyme and 5 µg/mL Sm2 nuclease, and disrupted by sonication. After centrifugation at 59,800 *g* at 4 °C for 30 min, supernatants were loaded onto a Ni-NTA agarose column (Qiagen). After washing with buffer A containing 5 mM imidazole and 0.2 % v/v Tween 20, the enzyme was eluted with buffer A containing 200 mM imidazole. The eluate was 10-times diluted with buffer B1 (50 mM Tris-HCl, pH 8.0, 10 % v/v glycerol, 1 mM DTT) and loaded onto a HiTrapQ column, which was pre-equilibrated with buffer B1. The enzyme was eluted with a 0–0.5 M NaCl gradient. The concentrated fractions containing the enzyme were further purified by Superdex 75 16/600 column. The buffer of the purified enzyme was substituted with buffer C (50 mM Tris-HCl, pH7.5, 200 mM NaCl, 5 mM DTT).

The collected Phm7 expressing cells were resuspended in buffer D (50 mM Tris-HCl pH7.5, 500 mM NaCl, 10 % v/v glycerol), and disruption and centrifugation were performed as in Fsa2. The supernatants were loaded onto a Ni-NTA agarose column and washed with buffer D containing 5 mM imidazole and 0.2% v/v Tween 20. The column was further washed with buffer D containing 30 mM imidazole, and the enzyme was eluted with buffer D containing 200 mM imidazole. The collected fractions containing the enzyme were concentrated and the His-tag was removed by thrombin digestion in a dialysis tube in buffer E (50 mM Tris-HCl, pH 7.5, 200 mM NaCl, 10 % v/v glycerol, 20 mM imidazole). Undigested enzyme and thrombin were removed using Ni-NTA and Benzamidine Sepharose 6B columns. The enzyme was further purified using resource Q and superdex75 16/600 columns.

The purified Fsa2 and Phm7 were concentrated to 30 and 15 mg/mL, respectively, in crystallization buffer (50 mM Tris-HCl, pH7.5, 200 mM NaCl, 5 mM DTT) using a 10 kDa cut-off Amicon Ultra-15 concentrator (Merck), and frozen at –80 °C until use. We confirmed that Fsa2 and Phm7 were monomeric states in solution using size-exclusion chromatography.

### Phm7 ligand screening using MST assay

To bypass the unavailability of the Phm7 substrate and cofactor for structural studies, ligand screening was conducted using MST assay^38^. Fluorescent dye-labelled Phm7 and Fsa2 were prepared using the Monolith Protein labelling kit RED-NHS 2nd generation and Monolith His-Tag Labelling Kit RED-tris-NTA 2G, respectively. MST measurements were performed using a Monolith NT.115 instrument (NanoTemper). In brief, each compound was twofold serially diluted and mixed with equal amounts of the labelled Phm7 (20 nM in 50 mM Tris-HCl, pH 8.0, 0.15 M NaCl, 10 mM MgCl_2_, 0.05% v/v Tween20) or Fsa2 (100 nM in 50 mM Tris-HCl, pH 8.0, 0.2 M NaCl, 0.05% v/v Tween20). After a short incubation, the compound-enzyme mixtures were loaded into glass capillaries and measured for initial fluorescence and change in fluorescence upon thermophoresis measurement. The data were analysed using the MO Affinity software.

### Crystallization of substrate-free Phm7 and Fsa2

Crystallization of the substrate-free forms of Phm7 and Fsa2 and SeMet-substituted Phm7 was performed by mixing 1 µL of the substrate-free enzyme solution with 1 µL of the reservoir solution using the sitting drop vapor diffusion method at 20 °C. Reservoir components for Phm7 were 0.1 M Tris-HCl, pH 7.0, 1.56-1.59 M ammonium sulfate, and 15-19 % v/v glycerol. For Fsa2, reservoir solution containing 0.1 M Bis-Tris-HCl, pH 7.0, and 26-29 % w/v polyethylene glycol 3,350 was used. Initially, small Phm7 multi-crystals were obtained, and 2–6 months were required to obtain large crystals suitable for data collection. Three-dimensional rhombus-shaped single crystals of Fsa2 were obtained within a week. Crystals of SeMet-substituted Phm7 were obtained by the same procedure used for the native Phm7 crystals.

### Crystallization of inhibitor 5-bound Phm7

**5**-bound Phm7 crystals were prepared by soaking substrate-free crystals in buffer containing 20 mM Tris-HCl, pH 7.0, 50 mM NaCl, 1.6 M ammonium sulfate, 10 % v/v glycerol, 10 mM **5**, 10 % v/v ethanol, and 10 % v/v dimethyl sulfoxide) for 1 h at 4 °C.

### Data collection and structural determination of Phm7 and Fsa2

All crystals were flash frozen in a cryo-stream at 100 K, and then the crystals were transferred into liquid nitrogen. Prior to flash freezing, Fsa2 crystals were soaked cryo-buffer A; 0.77 M Bis-Tris-HCl, pH 7.0, 30 % w/v polyethylene glycol 3,350, 50 % v/v ethanol for 45 min and kept in air for 5–10 s to dehydrate. Crystals of a substrate-free and SeMet-substituted Phm7 were soaked in cryo-buffer A; 20 mM Tris-HCl, pH 7.0, 0.1 M NaCl, 2 M ammonium sulfate, 20 % v/v glycerol for a few minutes, and crystals of **5**-bound Phm7 were soaked in cryo-buffer B; 50 mM Tris-HCl, pH 7.0, 2 M ammonium sulfate, 20 % v/v glycerol, 2.5 mM **5**, 2.5 % v/v dimethyl sulfoxide, 7.5 % v/v ethanol for a few minutes.

X-ray diffraction data were collected at SPring-8 beamlines BL26B1, BL32XU, BL41XU (Hyogo, Japan), or a Photon Factory beamline BL1A (Ibaraki, Japan). The collected data were integrated and reduced using the XDS package^52^, or AIMLESS^53^ in the CCP4 package.

Single-wavelength anomalous diffraction datasets of SeMet-substituted Phm7 crystals were collected at the wavelength of the absorption peak of the selenium atom (0.97920 Å). Substructure determination, phase calculation, and auto model building were performed using Phaser SAD pipeline^54^ in the CCP4 package. Three Phm7 molecules were found in the asymmetric unit, and a model of 841 out of 1158 residues were built. Manual model building was performed using Coot^55^, and Refmac5^56^ in CCP4 and PHENIX refine^57^ were used for structure refinement. We solved the crystal structures of substrate-free Phm7, **5**-bound Phm7, and substrate-free Fsa2 by the molecular replacement method using SeMet-substituted Phm7 as a search model. The model quality was validated by MolProbity^58^ in the PHENIX package. Ramachandran plot analysis showed that the residues of Fsa2, Phm7, and inhibitor-bound Phm7 in favoured (allowed) region were 96.59 (3.34), 97.66 (2.34), and 97.91 (1.82) %, respectively. The data collection and the final refinement statistics are summarized in Supplementary Table 2. Polder maps^59^ of **5** were calculated using PHENIX (Supplementary Fig. 27).

### Substrate docking simulation

Structural models of polyenoyl tetramic acids **3** and **4** were built and energy minimized using Chem3D v.16.0 (Perkin Elmer). The terminal region of polyenes, which are not involved in DA reaction, was partially restrained by AutoDockTools (v.1.5.6) to maintain the planarity of the C–C double bond. All molecules in the asymmetric unit of the inhibitor-bound Phm7 and the substrate-free Fsa2 crystals were used as receptors, and hydrogen atoms were added to the receptors. Side chains of E82, W223, and K356 of Phm7 were treated as flexible entities to facilitate the calculation, and Fsa2 was treated as a rigid model. The docking simulations were performed using AutoDock Vina^60^ under the following conditions: calculation grid boxes for Phm7 and Fsa2 were 44×52×40 Å^3^ and 34×40×38 Å^3^, and the exhaustiveness values were 150 and 100–200, respectively.

### Molecular dynamics (MD) simulations

The initial configurations of enzyme–substrate complexes, **4**•Phm7 and **3**•Fsa2, were built based on the results of precedent docking simulations. Each system was neutralized and solvated in 150 mM NaCl solution. All simulations were performed using the GENESIS program package (version 1.4.0)^61,62^. The AMBER ff14SB force field^63^ was used for protein and ions, while the TIP3P model^64^ was used for water molecules. The substrates were parameterized with the general AMBER force field parameter set version 2.1 (GAFF2) and AM1-BCC atomic charges using the antechamber module in Amber Tools 18^65,66^.

### gREST simulations

We defined the solute region as the dihedral angle and non-bonded energy (Coulomb and Lennard-Jones) terms of both the substrate and a set of selected binding site residues: D53, S66, Y68, E82, W223, W342, and K356 for Phm7 and the corresponding residues of Fsa2 (D51, G64, T66, Q80, W216, W332, and N346). Eight replicas were employed to cover the solute temperature range of 310.0–773.9 K (T = 310.0, 350.8, 396.7, 452.8, 519.6, 592.3, 675.0, and 773.9 K). We applied a flat-bottom potential to avoid the substrate being away from the binding site.

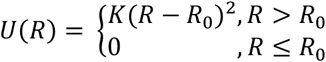

where *R, R*_0_, and *K* are the distances between the centre of masses of the substrate and binding site residues in the solute region, the flat-bottom distance, and the force constant, respectively. We set *R*_0_to 15.0 Å and *K* to 1 kcal/mol/Å^2^. The ligand feels no extra force in the region of *R* = 0–15 Å, while the harmonic restraint potential is applied beyond *R* = 15 Å to avoid substrate dissociation. Each of the **4**•Phm7 and **3**•Fsa2 systems was initially relaxed at different temperatures for 1 ns followed by production runs for 100 ns per replica (total sampling of 0.8 μs = 100 ns × 8 replicas). The trajectories at 310 K were used for the analysis. Additional details are provided in the Supplementary Notes.

### *In vitro* enzyme assay

The Δ*phm7* mutant derived from the **2**-producer fungus *Pyrenochaetopsis* sp. RK10-F058^9^ was cultured at 28 °C for 3 days in CYA medium. Mycelia were collected, washed, and frozen until use. The mycelia were ground to a fine powder under liquid nitrogen, and approximately 0.1 g of mycelial powder was suspended in 650 μL of extraction buffer (20 mM Tris-HCl, pH 7.5, 10 mM NaCl, 10 mM EDTA, 1% v/v Tween20, 45 μM tetracycline, 200 μM sinefungin). After centrifugation at 20,000 *× g* at 4 °C for 5 min, the **4**-saturated supernatant was used as substrate for the enzyme assay. Tetracycline, which was used as an external standard to normalize peak area, had no effect on the enzyme assay system. Conversion from **6** to **2**, which was mediated by intrinsic methyltransferase activity in the cell lysate, was completely inhibited by 200 μM sinefungin under the conditions tested.

Ten microliters of Phm7 or Fsa2 was added to 30 μL of the substrate and incubated at 25 °C. For evaluation of mutant Phm7 and Fsa2 activities, 100 ng of Phm7 and 10 μg of Fsa2 were used for the reactions, and incubated for 8 and 60 min, respectively. The enzyme reactions were terminated by adding 80 μL of ice-cold acetonitrile, followed by rapid quenching using liquid nitrogen. The reaction mixtures were thawed, centrifuged, and analysed by LC/ESI-MS using a Waters Acquity UPLC H-Class system fitted with a mass spectrometer (QDa, Waters). Comparison with authentic standards^9^ identified compounds corresponding to peaks **6, 7**, and **8** as *N*-demethylphomasetin and its derivatives containing **1**-type *trans*- and *cis*-decalin, respectively (Figs. 1b, 4b). An *m*/*z* value of 398.2334 [M–H]^−^, which was same as that of **6**–**8**, and a characteristic UV spectrum (λ_max_293, 305, 319 nm) for a tetraene substructure suggested the structure of linear pentanenoyl tetramic acid **4** as proposed, although it was too unstable to isolate and determine the structure by NMR.

The LC conditions were as follows: column, Waters Acquity UPLC BEH C18 (2.1 × 100 mm, 1.7 μm); flow rate, 0.25 mL/min; solvent A, water containing 0.05 % v/v aqueous formic acid; solvent B, acetonitrile containing 0.05 % v/v aqueous formic acid. After injection of the samples into a column equilibrated with 5 % solvent B, the column was developed with a linear gradient from 5 % to 60 % solvent B over 1 min and 60 % to 100 % over 4 min, followed by isocratic elution of 100 % solvent B for 6 min.

### Characterization of the mutated Phm7 in the producer fungus

The *phm7* gene on the chromosome of the **2**-producer fungus *Pyrenochaetopsis* sp. RK10-F058 was replaced with the mutated *phm7* gene under the control of a forced expression promoter (P_*tef1*_). Mutations were introduced into pBI121 containing the *ble*-P_*tef1*_-*phm7* cassette^9^ by PCR-based method as described above. Primer pairs used are listed in Supplementary Table 1. The strain RK10-F058 was transformed with the plasmid carrying the mutated *phm7* using *Agrobacterium tumefaciens*-mediated transformation, and transformants were isolated as described previously^9^. Correct replacements and introduced mutations were confirmed by PCR and direct sequencing. Metabolite production of the transformations carrying the mutated *phm7* was analysed by LC/MS as described previously^9^.

### Density functional theory (DFT) calculations

All calculations were carried out with the Gaussian 16 (revision B.01) program package^67^. The molecular structures and harmonic vibrational frequencies were obtained using the hybrid density functional method based on the M06-2X functional^68^. We used the 6-311+G** basis set^69^. The self-consistent reaction field (SCRF) method based on the conductor-like polarizable continuum model (CPCM)^70-73^ was employed to evaluate the solvent reaction field (water; *ε* = 78.39). Geometry optimization and vibrational analysis were performed at the same level. All stationary points were optimized without any symmetry assumptions and characterized by normal coordinate analysis at the same level of theory (number of imaginary frequencies, NIMAG, 0 for minima and 1 for TSs). The intrinsic reaction coordinate (IRC) method^74,75^ was used to track minimum energy paths from transition structures to the corresponding local minima.

## Data availability

Atomic coordinates and crystallographic structure factors have been deposited in the Protein Data Bank under accession codes 7E5T (Fsa2), 7E5U (Phm7), and 7E5V (inhibitor-bound Phm7). All other data are available from the corresponding author upon reasonable request.

## Acknowledgements

We acknowledge the computational resources provided by the RIKEN Advanced Center for Computing and Communication (HOKUSAI GreatWave and BigWaterfall) and the HPCI system (Project ID: hp200153) for the MD simulations and the Research Center for Computational Science (Okazaki, Japan) for the DFT calculations. This work was supported by MEXT/KAKENHI (Grant Numbers 19H04665, 20K05872 (to NK), 19K12229 (to SR), 19K06992 (to RT), 20K15273 (to KW), 19H05645 (to YS), 20H00416 (ST), 19H04658, 19H05780 (to SN)), MEXT as “Priority Issue on Post-K computer” (Building Innovative Drug Discovery Infrastructure Through Functional Control of Biomolecular Systems), the Takeda Science Foundation (to RT), and the Naito Foundation (to RT), and RIKEN Pioneering Research Projects (Dynamic Structural Biology/Glycolipidologue Initiative) (to YS).

## Author contributions

N.K., S.T., and S.N. designed the research. K.F., K.K, and T.H. performed enzyme purification and crystallography. Docking simulations were done by K.F. S.R. and Y.S. performed MD simulations. Site-directed mutagenesis and analysis of mutants were done by K.F., N.K., K.K., and T.N. Compound isolation and characterization were done by T.N. K.W. and R.T. performed DFT calculations. N.K., K.F., T.H., K.W., R.T., S.R., and S.N. wrote the manuscript. All authors discussed the data.

## References

1. Pasteur, L. Sur les relations qui peuvent exister entre la forme crystalline, la composition chimique et le sens de la polarization rotatoire. Annales Chimie Phys. 24, 442–459 (1848).

2. Clardy, J. & Walsh, C. Lessons from natural molecules. Nature 432, 829–837 (2004).

3. Nicolaou, K.C., Snyder, S.A., Montagnon, T. & Vassilikogiannakis, G. The Diels--Alder reaction in total synthesis. Angew Chem Int Ed Engl 41, 1668–98 (2002).

4. Minami, A. & Oikawa, H. Recent advances of Diels-Alderases involved in natural product biosynthesis. J Antibiot 69, 500–506 (2016).

5. Jeon, B.S., Wang, S.A., Ruszczycky, M.W. & Liu, H.W. Natural [4 + 2]-cyclases. Chem Rev 117, 5367–5388 (2017).

6. Jamieson, C.S., Ohashi, M., Liu, F., Tang, Y. & Houk, K.N. The expanding world of biosynthetic pericyclases: cooperation of experiment and theory for discovery. Nat Prod Rep 36, 698–713 (2019).

7. Lichman, B.R., O’Connor, S.E. & Kries, H. Biocatalytic strategies towards [4+2] cycloadditions. Chemistry 25, 6864–6877 (2019).

8. Kato, N. et al. A new enzyme involved in the control of the stereochemistry in the decalin formation during equisetin biosynthesis. Biochem Biophys Res Commun 460, 210–215 (2015).

9. Kato, N. et al. Control of the stereochemical course of [4+2] cycloaddition during trans-decalin formation by Fsa2-family enzymes. Angew Chem Int Ed Engl 57, 9754–9758 (2018).

10. Dickschat, J.S. Isoprenoids in three-dimensional space: the stereochemistry of terpene biosynthesis. Nat Prod Rep 28, 1917–1936 (2011).

11. Keatinge-Clay, A.T. Stereocontrol within polyketide assembly lines. Nat Prod Rep 33, 141–149 (2016).

12. Weissman, K.J. Polyketide stereocontrol: a study in chemical biology. Beilstein J Org Chem 13, 348–371 (2017).

13. Christianson, D.W. Structural and chemical biology of terpenoid cyclases. Chem Rev 117, 11570–11648 (2017).

14. Kim, H.J., Ruszczycky, M.W., Choi, S.H., Liu, Y.N. & Liu, H.W. Enzyme-catalysed [4+2] cycloaddition is a key step in the biosynthesis of spinosyn A. Nature 473, 109–112 (2011).

15. Fischbach, M.A. & Walsh, C.T. Assembly-line enzymology for polyketide and nonribosomal peptide antibiotics: logic, machinery, and mechanisms. Chem Rev 106, 3468–3496 (2006).

16. Fage, C.D. et al. The structure of SpnF, a standalone enzyme that catalyzes [4 + 2] cycloaddition. Nat Chem Biol 11, 256–258 (2015).

17. Tian, Z. et al. An enzymatic [4+2] cyclization cascade creates the pentacyclic core of pyrroindomycins. Nat Chem Biol 11, 259–265 (2015).

18. Zheng, Q. et al. Structural insights into a flavin-dependent [4 + 2] cyclase that catalyzes trans-decalin formation in pyrroindomycin biosynthesis. Cell Chem Biol 25, 718–727 e3 (2018).

19. Zheng, Q. et al. Enzyme-dependent [4 + 2] cycloaddition depends on lid-like interaction of the N-terminal sequence with the catalytic core in PyrI4. Cell Chem Biol 23, 352–360 (2016).

20. Yang, Z. et al. Influence of water and enzyme SpnF on the dynamics and energetics of the ambimodal [6+4]/[4+2] cycloaddition. Proc Natl Acad Sci U S A 115, E848–E855 (2018).

21. Dan, Q. et al. Fungal indole alkaloid biogenesis through evolution of a bifunctional reductase/Diels-Alderase. Nat Chem 11, 972–980 (2019).

22. Zou, Y. et al. Computational investigation of the mechanism of Diels-Alderase PyrI4. J Am Chem Soc 142, 20232–20239 (2020).

23. Schobert, R. & Schlenk, A. Tetramic and tetronic acids: an update on new derivatives and biological aspects. Bioorg Med Chem 16, 4203–4221 (2008).

24. Li, G., Kusari, S. & Spiteller, M. Natural products containing ‘decalin’ motif in microorganisms. Nat Prod Rep 31, 1175–1201 (2014).

25. Burmeister, H.R., Bennett, G.A., Vesonder, R.F. & Hesseltine, C.W. Antibiotic produced by Fusarium equiseti NRRL 5537. Antimicrob Agents Chemother 5, 634–639 (1974).

26. Singh, S.B. et al. Equisetin and a novel opposite stereochemical homolog phomasetin, two fungal metabolites as inhibitors of HIV-1 integrase. Tetrahedron Lett 39, 2243–2246 (1998).

27. Agatsuma, T. et al. UCS1025A and B, new antitumor antibiotics from the fungus Acremonium species. Org Lett 4, 4387–4390 (2002).

28. Nakai, R. et al. Telomerase inhibitors identified by a forward chemical genetics approach using a yeast strain with shortened telomere length. Chem Biol 13, 183–190 (2006).

29. Li, X., Zheng, Q., Yin, J., Liu, W. & Gao, S. Chemo-enzymatic synthesis of equisetin. Chem Commun 53, 4695–4697 (2017).

30. Li, L. et al. Biochemical characterization of a eukaryotic decalin-forming Diels-Alderase. J Am Chem Soc 138, 15837–15840 (2016).

31. Sato, M. et al. Involvement of lipocalin-like CghA in decalin-forming stereoselective intramolecular [4+2] cycloaddition. Chembiochem 16, 2294–2298 (2015).

32. Li, L. et al. Genome mining and assembly-line biosynthesis of the UCS1025A pyrrolizidinone family of fungal alkaloids. J Am Chem Soc 140, 2067–2071 (2018).

33. Tan, D. et al. Genome-mined Diels-Alderase catalyzes formation of the cis-octahydrodecalins of varicidin A and B. J Am Chem Soc 141, 769–773 (2019).

34. Kamiya, M. & Sugita, Y. Flexible selection of the solute region in replica exchange with solute tempering: Application to protein-folding simulations. J Chem Phys 149, 072304 (2018).

35. Re, S., Oshima, H., Kasahara, K., Kamiya, M. & Sugita, Y. Encounter complexes and hidden poses of kinase-inhibitor binding on the free-energy landscape. Proc Natl Acad Sci U S A 116, 18404–18409 (2019).

36. Niitsu, A., Re, S., Oshima, H., Kamiya, M. & Sugita, Y. De novo prediction of binders and nonbinders for T4 lysozyme by gREST simulations. J Chem Inf Model 59, 3879–3888 (2019).

37. Flower, D.R., North, A.C. & Sansom, C.E. The lipocalin protein family: structural and sequence overview. Biochim Biophys Acta 1482, 9–24 (2000).

38. Wienken, C.J., Baaske, P., Rothbauer, U., Braun, D. & Duhr, S. Protein-binding assays in biological liquids using microscale thermophoresis. Nat Commun 1, 100 (2010).

39. Sato, M. et al. Catalytic mechanism and endo-to-exo selectivity reversion of an octalinforming natural Diels–Alderase. Nat Catal (2021).

40. Taylor, M.S. & Jacobsen, E.N. Asymmetric catalysis by chiral hydrogen-bond donors. Angew Chem Int Ed Engl 45, 1520–1543 (2006).

41. Oikawa, H., Katayama, K., Suzuki, Y. & Ichihara, A. Enzymatic activity catalysing exoselective Diels–Alder reaction in solanapyrone biosynthesis. J Chem Soc, Chem Commun, 1321–1322 (1995).

42. Ose, T. et al. Insight into a natural Diels-Alder reaction from the structure of macrophomate synthase. Nature 422, 185–9 (2003).

43. Byrne, M.J. et al. The Catalytic mechanism of a natural Diels-Alderase revealed in molecular detail. J Am Chem Soc 138, 6095–6098 (2016).

44. Cogan, D.P. et al. Structural insights into enzymatic [4+2] aza-cycloaddition in thiopeptide antibiotic biosynthesis. Proc Natl Acad Sci U S A 114, 12928–12933 (2017).

45. Cai, Y. et al. Structural basis for stereoselective dehydration and hydrogen-bonding catalysis by the SAM-dependent pericyclase LepI. Nat Chem 11, 812–820 (2019).

46. Gao, L. et al. FAD-dependent enzyme-catalysed intermolecular [4+2] cycloaddition in natural product biosynthesis. Nat Chem 12, 620–628 (2020).

47. Little, R. et al. Unexpected enzyme-catalysed [4+2] cycloaddition and rearrangement in polyether antibiotic biosynthesis. Nat Catal 2, 1045–1054 (2019).

48. Chiu, H.J. et al. Structure of the first representative of Pfam family PF09410 (DUF2006) reveals a structural signature of the calycin superfamily that suggests a role in lipid metabolism. Acta Crystallogr F 66, 1153–1159 (2010).

49. Minami, A., Ugai, T., Ozaki, T. & Oikawa, H. Predicting the chemical space of fungal polyketides by phylogeny-based bioinformatics analysis of polyketide synthasenonribosomal peptide synthetase and its modification enzymes. Sci Rep 10, 13556 (2020).

50. Jang, J.H. et al. Fusarisetin A, an acinar morphogenesis inhibitor from a soil fungus, Fusarium sp. FN080326. J Am Chem Soc 133, 6865–6867 (2011).

51. Nogawa, T. et al. Wakodecalines A and B, new decaline metabolites isolated from a fungus Pyrenochaetopsis sp. RK10-F058. J Antibiot 71, 123–128 (2018).

52. Kabsch, W. XDS. Acta Crystallogr D 66, 125–132 (2010).

53. Evans, P.R. & Murshudov, G.N. How good are my data and what is the resolution? Acta Crystallogr D 69, 1204–1214 (2013).

54. McCoy, A.J. et al. Phaser crystallographic software. J Appl Crystallogr 40, 658–674 (2007).

55. Emsley, P., Lohkamp, B., Scott, W.G. & Cowtan, K. Features and development of Coot. Acta Crystallogr D 66, 486–501 (2010).

56. Murshudov, G.N. et al. REFMAC5 for the refinement of macromolecular crystal structures. Acta Crystallogr D 67, 355–367 (2011).

57. Afonine, P.V. et al. Towards automated crystallographic structure refinement with phenix.refine. Acta Crystallogr D 68, 352–367 (2012).

58. Williams, C.J. et al. MolProbity: More and better reference data for improved all-atom structure validation. Protein Sci 27, 293–315 (2018).

59. Liebschner, D. et al. Polder maps: improving OMIT maps by excluding bulk solvent. Acta Crystallogr D 73, 148–157 (2017).

60. Trott, O. & Olson, A.J. AutoDock Vina: improving the speed and accuracy of docking with a new scoring function, efficient optimization, and multithreading. J Comput Chem 31, 455–61 (2010).

61. Jung, J. et al. GENESIS: a hybrid-parallel and multi-scale molecular dynamics simulator with enhanced sampling algorithms for biomolecular and cellular simulations. Wiley Interdiscip Rev Comput Mol Sci 5, 310–323 (2015).

62. Kobayashi, C. et al. GENESIS 1.1: A hybrid-parallel molecular dynamics simulator with enhanced sampling algorithms on multiple computational platforms. J Comput Chem 38, 2193–2206 (2017).

63. Maier, J.A. et al. ff14SB: Improving the accuracy of protein side chain and backbone parameters from ff99SB. J Chem Theory Comput 11, 3696–3713 (2015).

64. Jorgensen, W.L., Chandrasekhar, J., Madura, J.D., Impey, R.W. & Klein, M.L. Comparison of simple potential functions for simulating liquid water. J Chem Phys 79, 926–935 (1983).

65. Wang, J., Wolf, R.M., Caldwell, J.W., Kollman, P.A. & Case, D.A. Development and testing of a general amber force field. J Comput Chem 25, 1157–1174 (2004).

66. Case, D.A. et al. AMBER 2018. University of California, San Francisco. (2018).

67. Frisch, M.J. et al. Gaussian 16 Rev. B.01. (Wallingford, CT, 2016).

68. Zhao, Y. & Truhlar, D.G. The M06 suite of density functionals for main group thermochemistry, thermochemical kinetics, noncovalent interactions, excited states, and transition elements: two new functionals and systematic testing of four M06-class functionals and 12 other functionals. Theor Chem Account 120, 215–241 (2008).

69. Frisch, M.J., Pople, J.A. & Binkley, J.S. Self-consistent molecular orbital methods 25. Supplementary functions for Gaussian basis sets. J Chem Phys 80, 3265–3269 (1984).

70. Klamt, A. & Schuurmann, G. COSMO: a new approach to dielectric screening in solvents with explicit expressions for the screening energy and its gradient. J Chem Soc, Perkin Trans 2, 799-805 (1993).

71. Andzelm, J., Kölmel, C. & Klamt, A. Incorporation of solvent effects into density functional calculations of molecular energies and geometries. J Chem Phys 103, 9312–9320 (1995).

72. Barone, V. & Cossi, M. Quantum Calculation of Molecular Energies and Energy Gradients in Solution by a Conductor Solvent Model. J Phys Chem A 102, 1995–2001 (1998).

73. Cossi, M., Rega, N., Scalmani, G. & Barone, V. Energies, structures, and electronic properties of molecules in solution with the C-PCM solvation model. J Comput Chem 24, 669–681 (2003).

74. Ishida, K., Morokuma, K. & Komornicki, A. The intrinsic reaction coordinate. An ab initio calculation for HNC→HCN and H−+CH4→CH4+H−?J Chem Phys 66, 2153–2156 (1977).

75. Fukui, K. The path of chemical reactions - the IRC approach. Acc Chem Res 14, 363–368 (1981).

