## Supplementary information for "Molecular basis for two stereoselective Diels-Alderases that produce decalin skeletons"

### TABLE OF CONTENTS

#### Supplementary Tables

Supplementary Table 1. Oligonucleotides used in this study. 3
Supplementary Table 2. Crystallographic statistics. 5

#### Supplementary Figures

Supplementary Fig. 1. Natural products involving enzyme-mediated Diels-Alder (DA) reaction. 6
Supplementary Fig. 2. Compounds **1**–**9** and the corresponding reactions involving DSs. 7
Supplementary Fig. 3. Structural comparison of lipocalin-folds of DSs with other proteins. 8
Supplementary Fig. 4. Phm7 ligand screening using microscale thermophoresis assay. 9
Supplementary Fig. 5. Evaluation of Phm7 inhibitory activity of **5**. 10
Supplementary Fig. 6. Evaluation of Fsa2 inhibitory activity of **5**. 11
Supplementary Fig. 7. An overall structure of inhibitor-bound Phm7. 12
Supplementary Fig. 8. Docking poses of **4** in Phm7 and **3** in Fsa2 calculated by AutoDock Vina. 13
Supplementary Fig. 9. Major bound poses sampled by the MD simulations. 14
Supplementary Fig. 10. Electrostatic potential maps of the substrate binding site of Phm7 and Fsa2. 15
Supplementary Fig. 11. Hydrophobicity maps of the substrate binding site of Phm7 and Fsa2. 16
Supplementary Fig. 12. Conformations of the substrate **4** and **3** in the Phm7 and Fsa2 pockets. 17
Supplementary Fig. 13. Distribution of the dihedral angles of the substrates in the enzyme pockets. 18
Supplementary Fig. 14. Substrate–enzyme interactions at bound states. 19
Supplementary Fig. 15. Time course of the *in vitro* Phm7 reaction. 20
Supplementary Fig. 16. Location of the amino acid residues examined in this study. 20
Supplementary Fig. 17. *In vitro* analysis of Phm7 variants substituted with F and others. 21
Supplementary Fig. 18. Evaluation of the Phm7 mutants in the producer fungus. 22
Supplementary Fig. 19. DFT calculation with a model of K356A. 23
Supplementary Fig. 20. DFT calculation of the effects of N84 and E51 residues. 23
Supplementary Fig. 21. Comparison between a structurally related bacterial protein and Phm7. 24
Supplementary Fig. 22. Multiple sequence alignment of Fsa2-family DSs. 25
Supplementary Fig. 23. The substrate binding pocket in the homology models. 26
Supplementary Fig. 24. The convergent poses of **4** in the Phm7 pocket. 27
Supplementary Fig. 25. Size and shape of binding pockets. 28
Supplementary Fig. 26. Conformational changes of **3** along the tetramic acid–N346 distance. 29
Supplementary Fig. 27. A polder map of inhibitor **5** in the Phm7 substrate binding site. 30

#### Supplementary Notes

Molecular dynamics (MD) simulations 31
Purification and structure determination of 17-hydroxyphomasetin **9** 34
Cartesian coordinates and energies obtained by DFT calculations 40
**Supplementary References** 59

**Supplementary Table 1.** Oligonucleotides used in this study.

| Oligonucleotide | Sequence (5'→3') |
|---|---|
| ***E. coli*-expression plasmid construction** | |
| NdeI_Phm7_F | CATATGTCAGAACCAACGTCCTCGTCTTCG |
| XhoI_Phm7_R | CTCGAGCTAGGGTACCGGGATCTGCAGCTG |
| NdeI_Fsa2_F | CATATGTCCAACGTCACAGTCTCTGCCTTT |
| XhoI_Fsa2_R | CTCGAGTTAGCTTAGTTCGCATTGTCCACC |
| Fsa2_modified pET_F | ATGTCCAACGTCACAGTCTCTGCC |
| Fsa2_modified pET_R | GTGATGATGATGATGATGGCTGCT |
| *Restriction sites attached are underlined. | |
| **PCR-based site-directed mutagenesis** | |
| Phm7_L49A_F | TGGGAGGCTTGGGAATTCGACACATTT |
| Phm7_L49A_R | TTCCCAAGCCTCCCAGGAGTCGACAGA |
| Phm7_E51A_F | CTTTGGGCATTCGACACATTTGACACC |
| Phm7_E51A_R | GTCGAATGCCCAAAGCTCCCAGGAGTC |
| Phm7_D53A_F | GAATTCGCCACATTTGACACCAACGGA |
| Phm7_D53A_R | AAATGTGGCGAATTCCCAAAGCTCCCA |
| Phm7_S66A_F | GGCTGTGCCTTGTACCGCGATGCCCGA |
| Phm7_S66A_R | GTACAAGGCACAGCCGAATGCGACAGA |
| Phm7_Y68A_F | TCCTTGGCCCGCGATGCCCGAGGCGTT |
| Phm7_Y68A_R | ATCGCGGGCCAAGGAACAGCCGAATGC |
| Phm7_E82A_F | CACGCCGCAGTCAATGCTCTCTGGCCC |
| Phm7_E82A_R | ATTGACTGCGGCGTGGAAGCCACCCTG |
| Phm7_N84A_F | GAAGTCGCTGCTCTCTGGCCCGACGGC |
| Phm7_N84A_R | GAGAGCAGCGACTTCGGCGTGGAAGCC |
| Phm7_Y178A_F | GTGTACGCCACATTCCCTATGGGCCCT |
| Phm7_Y178A_R | GAATGTGGCGTACACCCCGGGGCAGAG |
| Phm7_W223A_F | AGGCCTGCGCCGACATTCATGAACGAT |
| Phm7_W223A_R | TGTCGGCGCAGGCCTTGCAGACCATCC |
| Phm7_W223F_F | AGGCCTTTCCCGACATTCATGAACGAT |
| Phm7_W223F_R | TGTCGGGAAAGGCCTTGCAGACCATCC |
| Phm7_A230F_F | AACGATTTCTATTACGTGGTCGCACAA |
| Phm7_A230F_R | GTAATAGAAATCGTTCATGAATGTCGG |
| Phm7_A230S_F | AACGATTCGTATTACGTGGTCGCACAA |
| Phm7_A230S_R | GTAATACGAATCGTTCATGAATGTCGG |
| Phm7_L245A_F | CAGATAGCTCGCACTTTAGGCTCCGTG |
| Phm7_L245A_R | AGTGCGAGCTATCTGCAGCATGTACGG |
| Phm7_L245F_F | CAGATATTCCGCACTTTAGGCTCCGTG |
| Phm7_L245F_R | AGTGCGGAATATCTGCAGCATGTACGG |
| Phm7_L245V_F | CAGATAGTCCGCACTTTAGGCTCCGTG |
| Phm7_L245V_R | AGTGCGGACTATCTGCAGCATGTACGG |
| Phm7_T247A_F | CTTCGCGCTTTAGGCTCCGTGTTCGTG |
| Phm7_T247A_R | GCCTAAAGCGCGAAGTATCTGCAGCAT |
| Phm7_T247F_F | CTTCGCTTCTTAGGCTCCGTGTTCGTG |
| Phm7_T247F_R | GCCTAAGAAGCGAAGTATCTGCAGCAT |
| Phm7_W342A_F | GCAGTGGCGAGCGAGCCCACCAGCGCA |

| | |
|---|---|
| Phm7_W342A_R | CTCGCTCGCCACTGCTCGCTTATGCGA |
| Phm7_W342F_F | GCAGTGTTCAGCGAGCCCACCAGCGCA |
| Phm7_W342F_R | CTCGCTGAACACTGCTCGCTTATGCGA |
| Phm7_W342L_F | GCAGTGCTGAGCGAGCCCACCAGCGCA |
| Phm7_W342L_R | CTCGCTCAGCACTGCTCGCTTATGCGA |
| Phm7_K356A_F | ACAGGCGCGTCAGGTTGGATTGAGGCA |
| Phm7_K356A_R | ACCTGACGCGCCTGTTCCATCAGGTCC |
| Phm7_K356F_F | ACAGGCTTCTCAGGTTGGATTGAGGCA |
| Phm7_K356F_R | ACCTGAGAAGCCTGTTCCATCAGGTCC |
| Phm7_L381A_F | GGTCAGGCGCAGATCCCGGTACCCTAG |
| Phm7_L381A_R | GATCTGCGCCTGACCTCCAAATCCATG |
| Phm7_L381F_F | GGTCAGTTCCAGATCCCGGTACCCTAG |
| Phm7_L381F_R | GATCTGGAACTGACCTCCAAATCCATG |
| Fsa2_Q80A_F | GTTTCAAAGTCGCGGTTTTTGTGATCTGGGCTGATGAACG |
| Fsa2_Q80A_R | CACAAAAACCGCGACTTTGAAACCGCCGTGCTTG |
| Fsa2_W216A_F | CGTTAAGTGCGCCTCAAGTCATGACTGAGTCATACTACCTCC |
| Fsa2_W216A_R | CTTGAGGCGCACTTAACGGCGACCAAACGCGATC |
| Fsa2_W332A_F | GCATCATCGCGAACACTCCGACAAGTCGACCTGG |
| Fsa2_W332A_R | GGAGTGTTCGCGATGATGCGTTCATGACGAACCTGG |
| Fsa2_N346A_F | CCACTGGTGCCACGGGATTTGTGGAAGTACTTTGTGG |
| Fsa2_N346A_R | ATCCCGTGGCACCAGTGGCATCGGGTCCAG |
| **Genotyping of the genetically modified fungi** | |
| genotype_AF | CAGATCTAGGTGTGTCAG |
| genotype_AR | TAGGGCCGTATCGATTTC |
| genotype_BF | TTTGACGGTTGTGGATGATTTGTG |
| genotype_BR | GAGATGTTGCTGAAGTCG |
| genotype_CF | TCGTTGTTAGTCGGCAGATG |
| genotype_CR | CGCTCGAAGGCTTTAATTTGC |
| genotype_DF | TCTTTCGGGCGCTGACAA |
| genotype_DR | GCAGACAGAGACGGCGTT |

**Supplementary Table 2.** Crystallographic statistics.

| | Phm7 | | | Fsa2 |
|---|---|---|---|---|
| | SeMet derivative | Apo form | Inhibitor **5**-bond form | |
| ***Data Collection*** | | | | |
| Beam source | BL41XU (SPring-8) | BL32XU (SPring-8) | BL32XU (SPring-8) | BL32XU (SPring-8) |
| Wavelength (Å) | 0.979200 | 0.999994 | 1.000000 | 0.979400 |
| Resolution range (Å) | 49.42–2.17 (2.25–2.17) | 43.81–1.62 (1.68–1.62) | 45.18–1.61 (1.67–1.61) | 44.64–2.17 (2.25–2.17) |
| Space group | *C*2 | *C*2 | *C*2 | $P2_1$ |
| Unit cell parameters | | | | |
| a, b, c (Å) | a = 91.30, b = 151.71, c = 99.69 | a = 91.21, b = 150.48, c = 99.33 | a = 91.05, b = 149.88, c = 99.27 | a = 133.93, b = 80.23, c = 135.16 |
| α, β, γ (°) | α = γ = 90, β = 97.501 | α = γ = 90, β = 97.35 | α = γ = 90, β = 97.12 | α = γ = 90, β = 108.53 |
| Total reflections | 952909 (92401) | 583338 (59434) | 599244 (60565) | 496700 (49137) |
| Unique reflections | 70874 (7022) | 167533 (16704) | 170041 (16956) | 141327 (13856) |
| Multiplicity | 13.4 (13.2) | 3.5 (3.6) | 3.5 (3.5) | 3.5 (3.5) |
| Completeness (%) | 99.97 (99.99) | 99.76 (99.80) | 99.82 (99.78) | 97.61 (96.96) |
| Mean *I* / sigma(*I*) | 12.06 (2.66) | 6.90 (1.58) | 12.73 (2.66) | 8.48 (2.60) |
| Wilson B-factor | 24.76 | 20.29 | 21.66 | 29.05 |
| $R_{merge}$ (%) | 0.179 (0.906) | 0.107 (0.651) | 0.061 (0.447) | 0.109 (0.533) |
| $R_{meas}$ (%) | 0.185 (0.943) | 0.126 (0.769) | 0.072 (0.526) | 0.129 (0.629) |
| $CC_{1/2}$ | 0.997 (0.832) | 0.988 (0.674) | 0.996 (0.815) | 0.998 (0.780) |
| ***Refinement*** | | | | |
| Number of reflections | | 167510 (16704) | 169780 (16956) | 140531 (13855) |
| $R_{work}$ / $R_{free}$ | | 0.1894 / 0.2215 | 0.1910 / 0.2147 | 0.1798 / 0.2352 |
| Number of atoms | | | | |
| Protein | | 8647 | 8470 | 22909 |
| Inhibitor **5** | | - | 84 | - |
| Water | | 820 | 792 | 50 |
| Other ligand / Ion | | 176 | 108 | 1226 |
| RMSD | | | | |
| Bond lengths (Å) | | 0.014 | 0.035 | 0.004 |
| Bond angles (°) | | 1.26 | 1.93 | 0.70 |
| Average B-factor | | | | |
| Protein | | 25.22 | 26.97 | 31.15 |
| Inhibitor **5** | | - | 34.38 | - |
| Water | | 34.42 | 36.33 | 31.54 |
| Other ligand / Ion | | 48.00 | 45.24 | 35.98 |

※ Other molecules were containing sulfate ion, glycerol, and polyethylene glycol molecules.

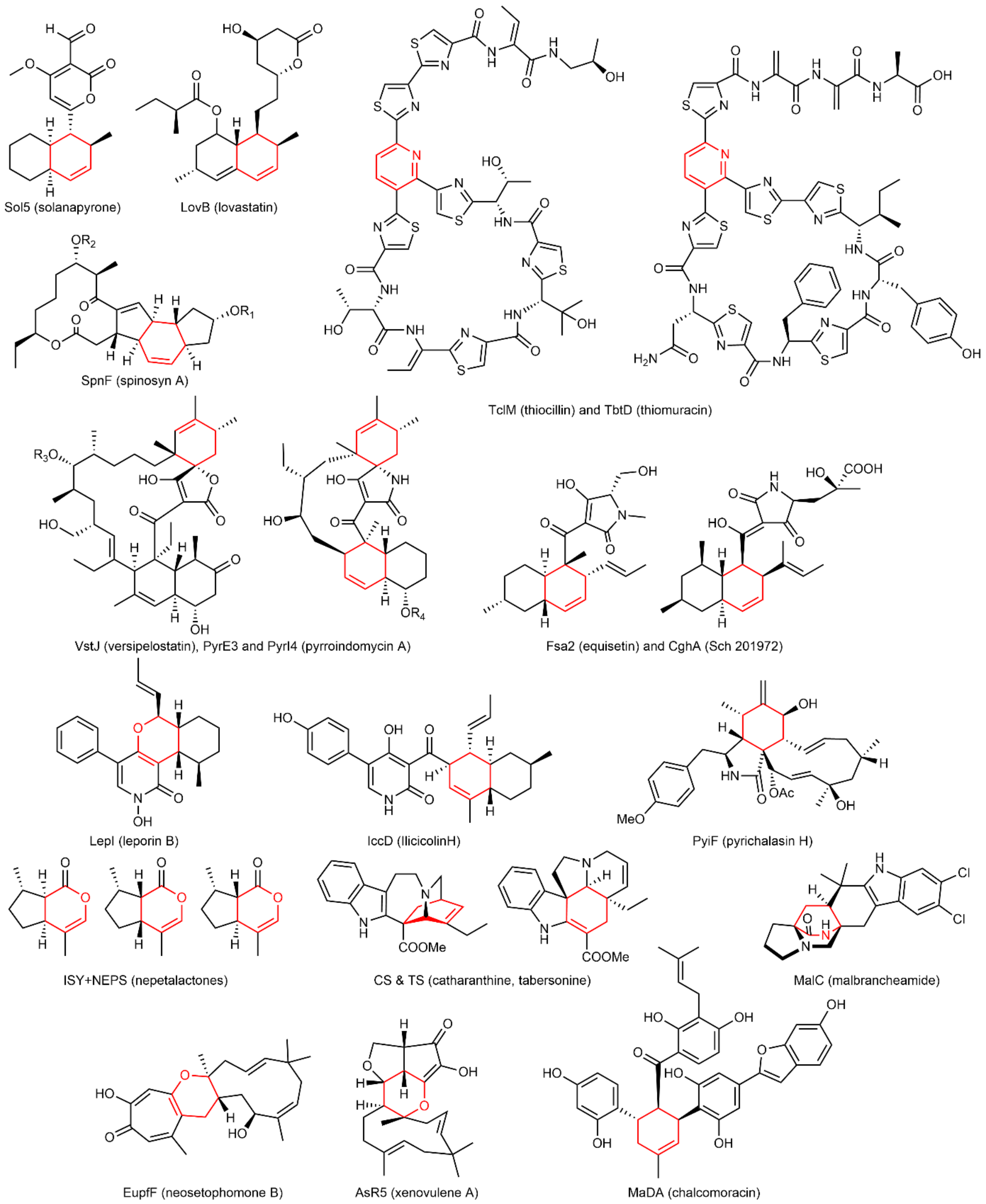


**Supplementary Fig. 1.** Natural products involving enzyme-mediated Diels-Alder (DA) reaction.

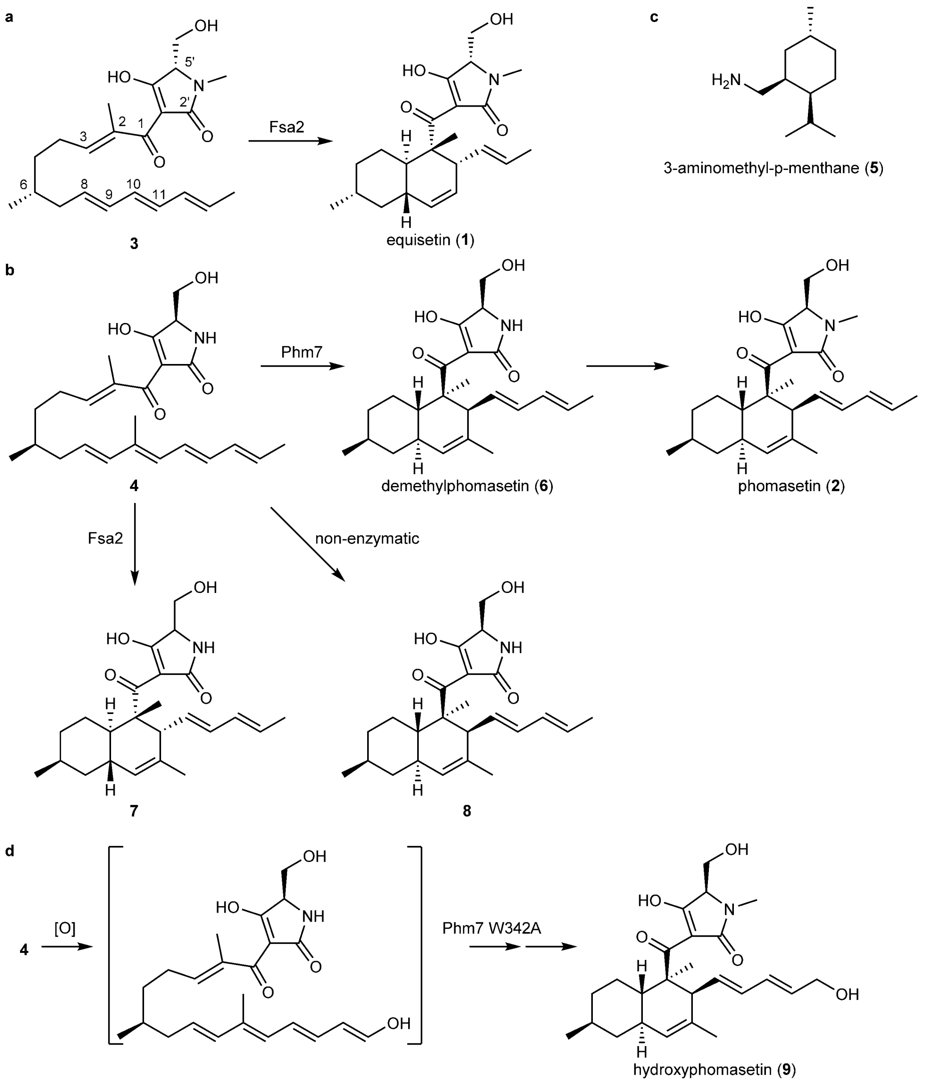


**Supplementary Fig. 2.** Compounds **1**–**9** and the corresponding reactions involving DSs. **a**, Equisetin (**1**) is formed from linear tetraenoyl tetramic acid **3** via Fsa2-mediated DA reaction, which was experimentally confirmed using a synthetic substrate[1]. **b**, pentaenoyl tetramic acid **4** is a likely Phm7 substrate to yield *N*-demethylphomasetin (**6**), which is further converted to phomasetin (**2**). Fsa2 exclusively forms a derivative of **6** containing **1**-type decalin, **7**, from **4**. A *cis*-decalin derivative **8** as well as **6** were produced in the *phm7* deletion mutant derived from a **2**-producer fungus, *Pyrenochaetopsis* sp. RK10-F058[2]. **c**, A DS inhibitor, 3-aminomethyl-*p*-menthane (**5**). **d**, A new derivative of **2** with a hydroxy group at the terminus of polyene (**9**) identified in this study, which was produced in the genetically modified fungus carrying the *phm7* mutant gene.

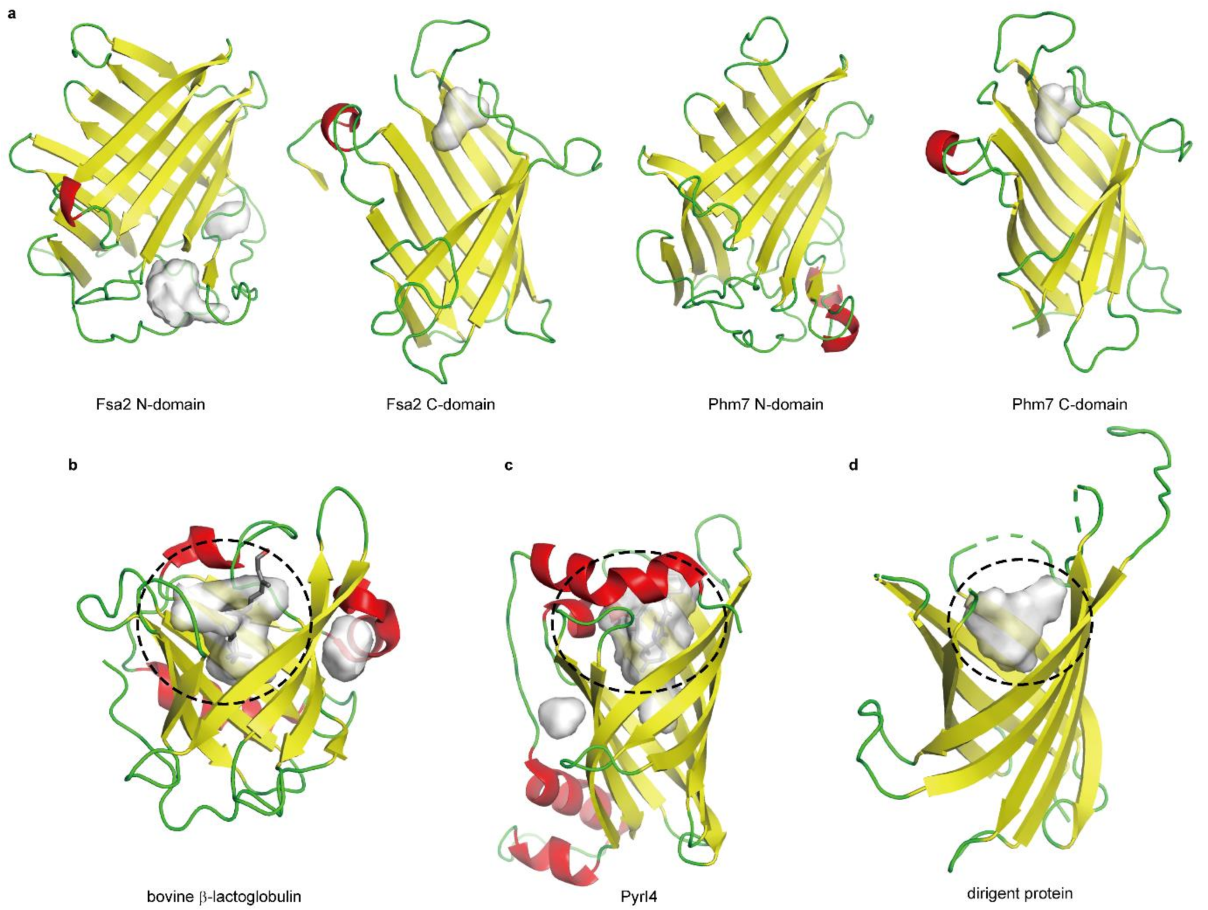


**Supplementary Fig. 3.** Structural comparison of lipocalin-folds of DSs with other proteins. **a**, N- and C-domains of Phm7 and Fsa2; **b**, bovine β-lactoglobulin (1GX8) as a typical lipocalin protein[3]; **c**, PyrI4 (5BU3) catalysing DA reaction in the pyrroindomycin biosynthesis[4,5]; **d**, dirigent protein (4REV) responsible for the stereoselective dimerization of coniferyl alcohols[6]. Secondary structures, such as α-helix, β-sheet, and loop, are indicated by red, yellow, and green, respectively. Cavities inside the β-barrel, indicated by dash line, serve as ligand-binding and active sites of these proteins.

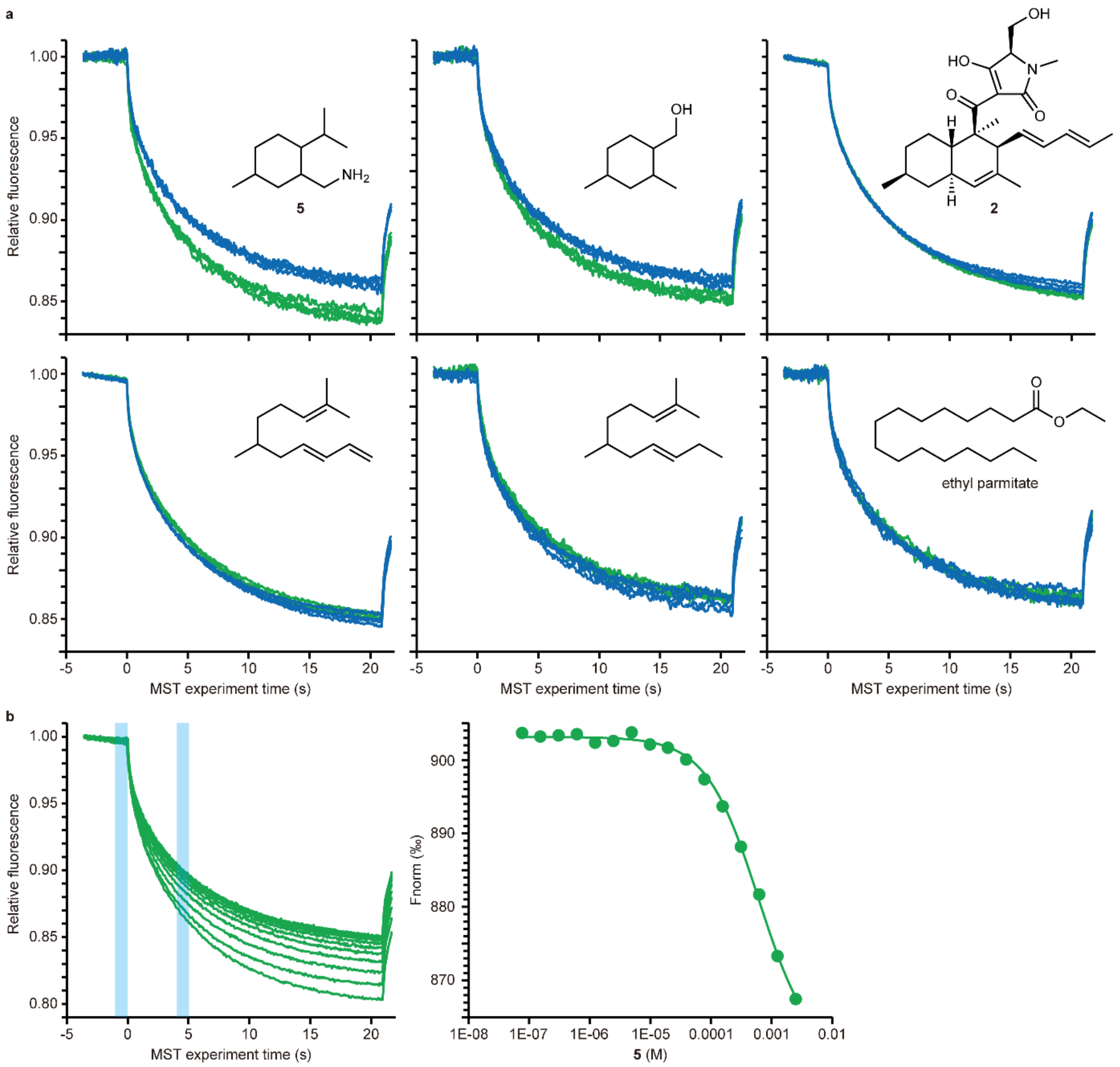


**Supplementary Fig. 4.** Phm7 ligand screening using microscale thermophoresis (MST) assay. **a**, MST traces of four independent experiments of Phm7 alone (blue) and Phm7 with 0.5 mM compound (green, except for **2**, 242 μM). The thermophoresis was monitored at 2 % excitation power and medium MST power. Commercially available small molecules possessing partial structures similar to the linear polyene substrate or cyclized product of the Phm7 reaction were screened, and some substituted cyclohexane compounds were found to bind to Phm7. **b**, Binding affinity of Phm7 for **5**. The MST traces of Phm7 with a serial dilution of **5** (2.5 mM to 76 nM, left panel). Concentration dependent change in fluorescence of the labelled Phm7 on interaction with **5** (right panel). It showed a moderate binding affinity for Phm7 with a $K_d$ value of 432 μM.

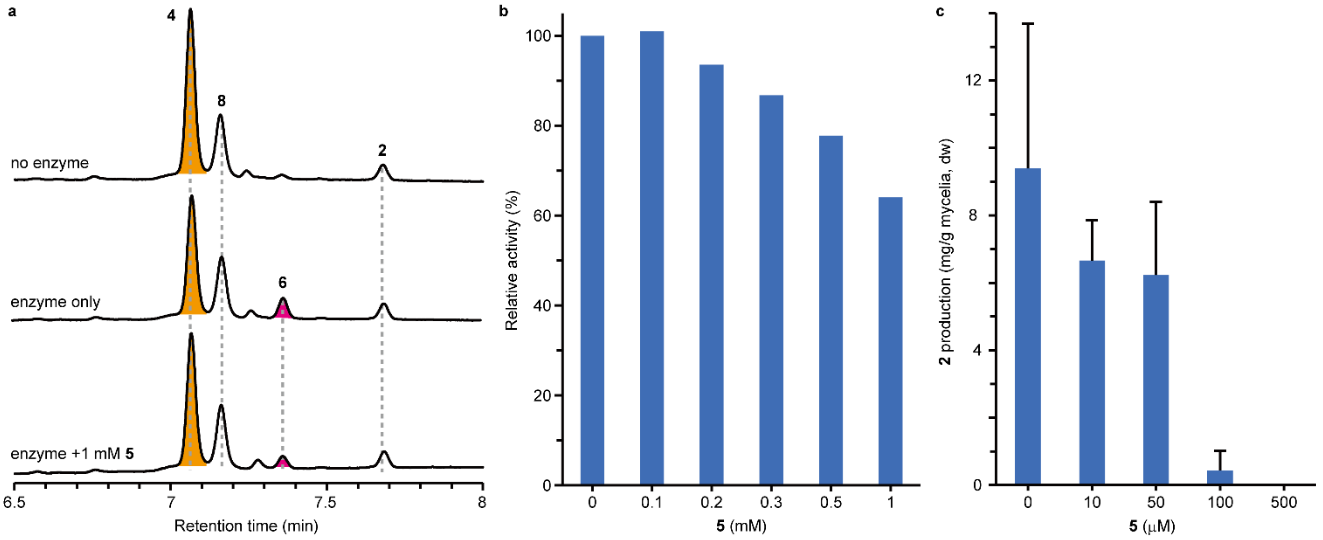


**Supplementary Fig. 5.** Evaluation of Phm7 inhibitory activity of **5**. **a**, UPLC traces of the Phm7 reaction mixtures with and without **5**. **b**, A concentration-dependent inhibition of the Phm7 activity by **5**. Phm7 (60 nM) was incubated in the Δ*phm7* cell lysate in the presence of **5** at 25 °C for 8 min. The reaction products were analysed by LC/MS. Compound **5** inhibited Phm7-mediated formation of **6** *in vitro*, although the $IC_{50}$ value was >1 mM under the conditions tested. **c**, A concentration-dependent inhibition of production of **2** in *Pyrenochaetopsis* sp. RK10-F058 by **5**. The fungus was cultured in CYA medium at 28 °C for 8 days, and their culture extracts were analysed by LC/ESI-MS. A dose-dependent inhibition of the production of **2** in the fungus was observed when **5** was added to the liquid culture medium.

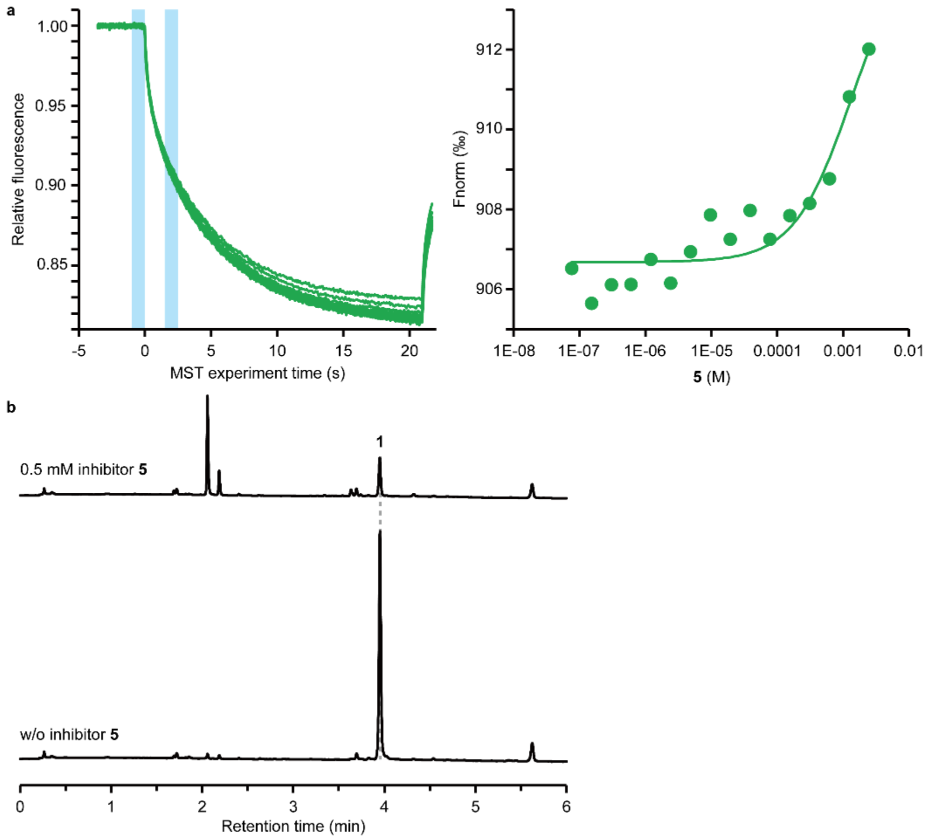


**Supplementary Fig. 6.** Evaluation of Fsa2 inhibitory activity of **5**. **a**, Binding affinity of Fsa2 for **5**. The MST traces of Fsa2 with a serial dilution of **5** (2.5 mM to 76 nM, left panel). A concentration-dependent change in fluorescence of the labelled Fsa2 on interaction with **5** (right panel). **b**, Inhibition of production of **1** in *Fusarium* sp. FN080326 by **5**. The fungus was cultured on PDB plate medium containing 0.5 mM **5** at 28 °C for 12 days, and culture extracts were analysed by LC/ESI-MS.

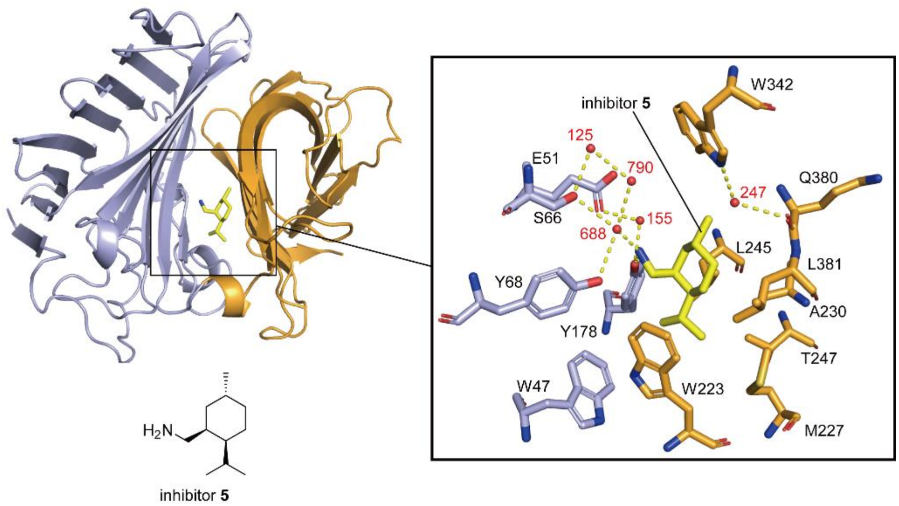


**Supplementary Fig. 7.** An overall structure of inhibitor-bound Phm7 and close-up view of the inhibitor–enzyme interaction. Dashed lines show hydrogen bonds. Red spheres represent water molecules.

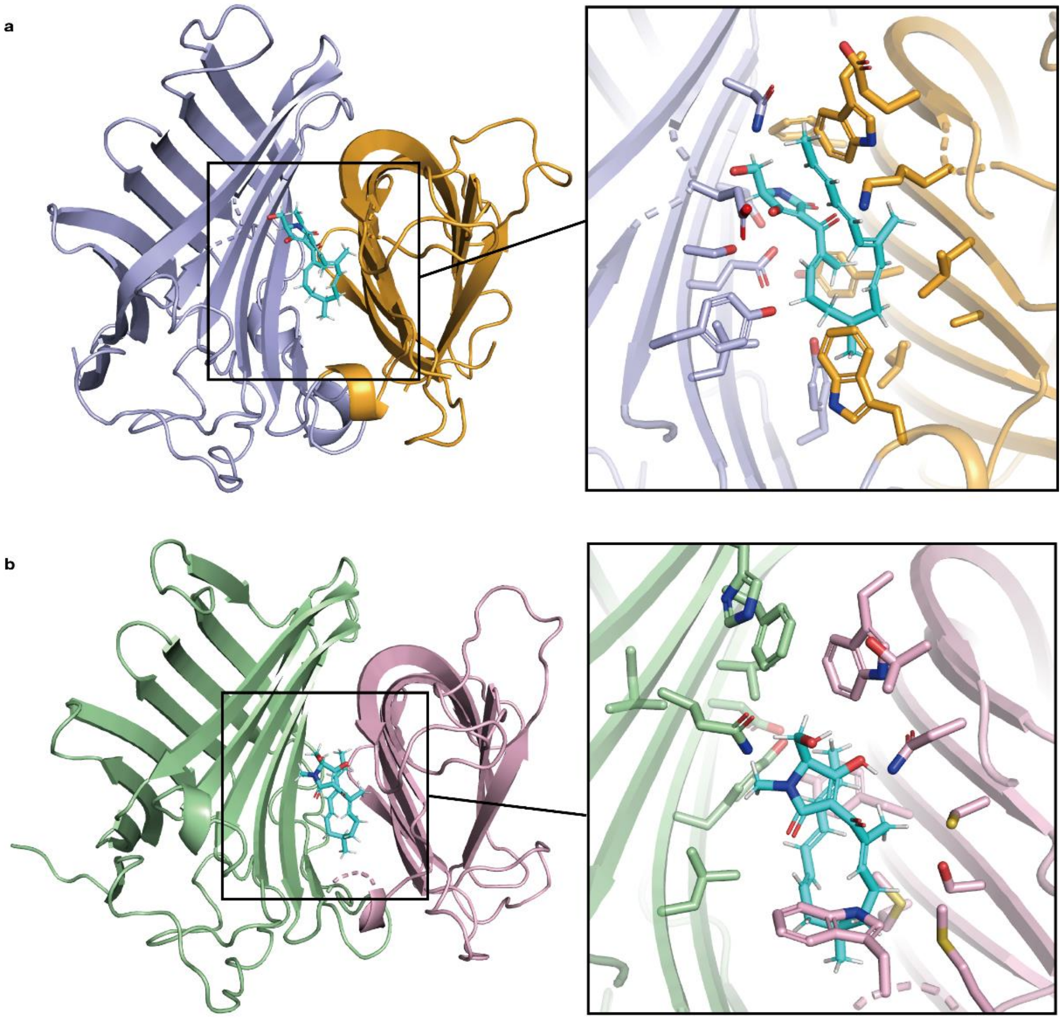


**Supplementary Fig. 8.** Docking poses of **4** in Phm7 (**a**) and **3** in Fsa2 (**b**) calculated by AutoDock Vina. Docking simulations were performed using crystal structures of inhibitor-bound form of Phm7 and substrate-free Fsa2. Substrates and surrounding amino acid residues are shown as stick models.

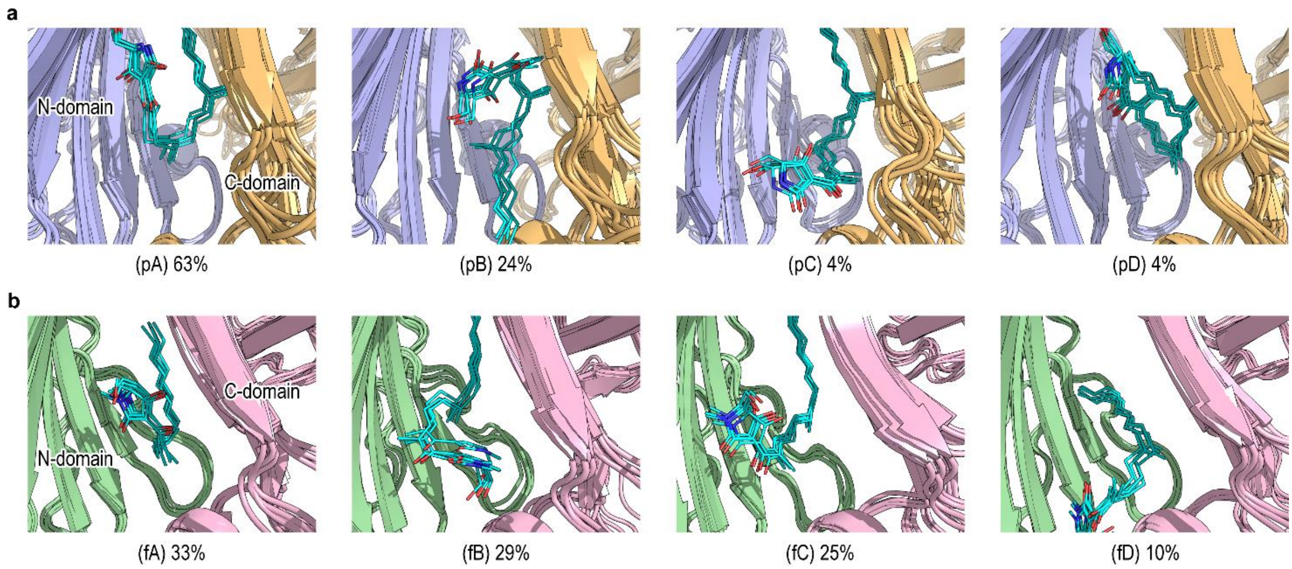


**Supplementary Fig. 9.** Major bound poses sampled by the gREST simulations. Four clusters were obtained from the clustering analysis using DBSCAN[7] for each of **4**•Phm7 (pA–pD, **a**) and **3**•Fsa2 (fA–fD, **b**). Of each cluster, the five structures proximate to the cluster centre are collectively shown. The relative population of each cluster is also given. For **4**•Phm7, the dominant pose (pA: 63%) has “folded” form, which is attainable the DA reaction. We found another “folded” form, but as a minor species (pD: 4%). The “extended” form is less populated (pB: 24% and pC: 4%) and unlikely proceeds the reaction without a large structural reorganization. In **3**•Fsa2, the “folded” form (fA: 33%) slightly dominates over the other three “extended” forms. The “extended” form (fC: 25%), in which the tetramic acid tends to head upward, can be considered as a half-folded conformation leading to the dominant “folded” form (fA: 33%). From these results, we consider that two dominant “folded” forms, pA and fA, represent the bound poses relevant to the DA reaction for **4**•Phm7 and **3**•Fsa2, respectively.

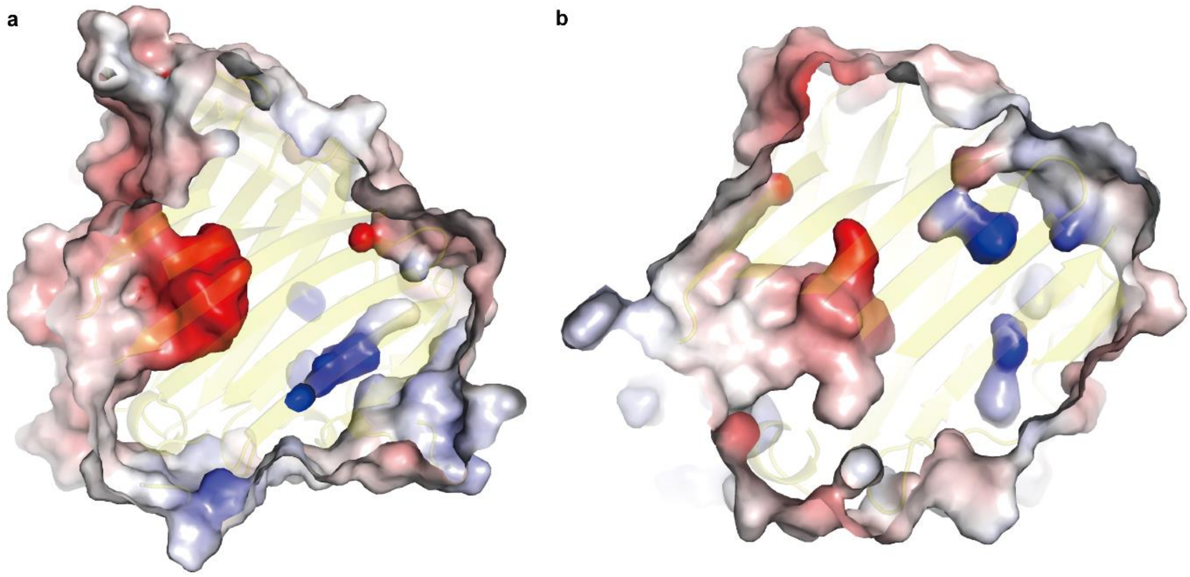


**Supplementary Fig. 10.** Electrostatic potential maps of the substrate binding site of Phm7 (**a**) and Fsa2 (**b**). Positively and negatively charge regions are indicated by blue and red, respectively.

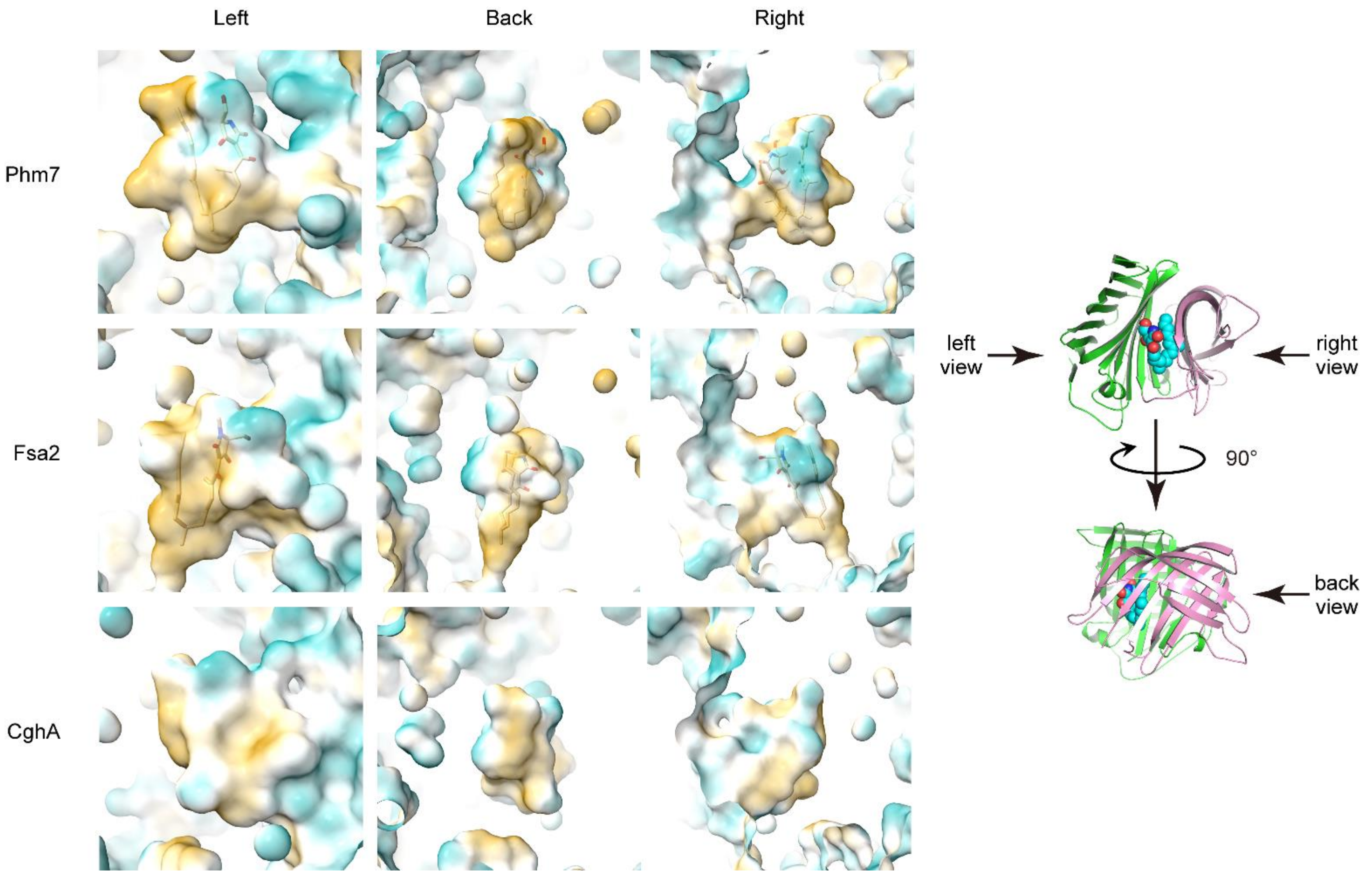


**Supplementary Fig. 11.** Surface models showing hydrophobicity maps of the substrate binding pocket in the substrate-bound Phm7 (top) and the substrate-bound Fsa2 (middle) predicted by MD simulations and crystal structure of the substrate-free CghA (PDB accession code 6KAW, bottom). The 50 % transparent molecular surfaces are displayed with colors ranging from goldenrod (hydrophobic) through white (intermediate) to cyan (hydrophilic) according to the hydrophobicity map value. The enzyme-bound substrates are shown as stick models for Phm7 and Fsa2.

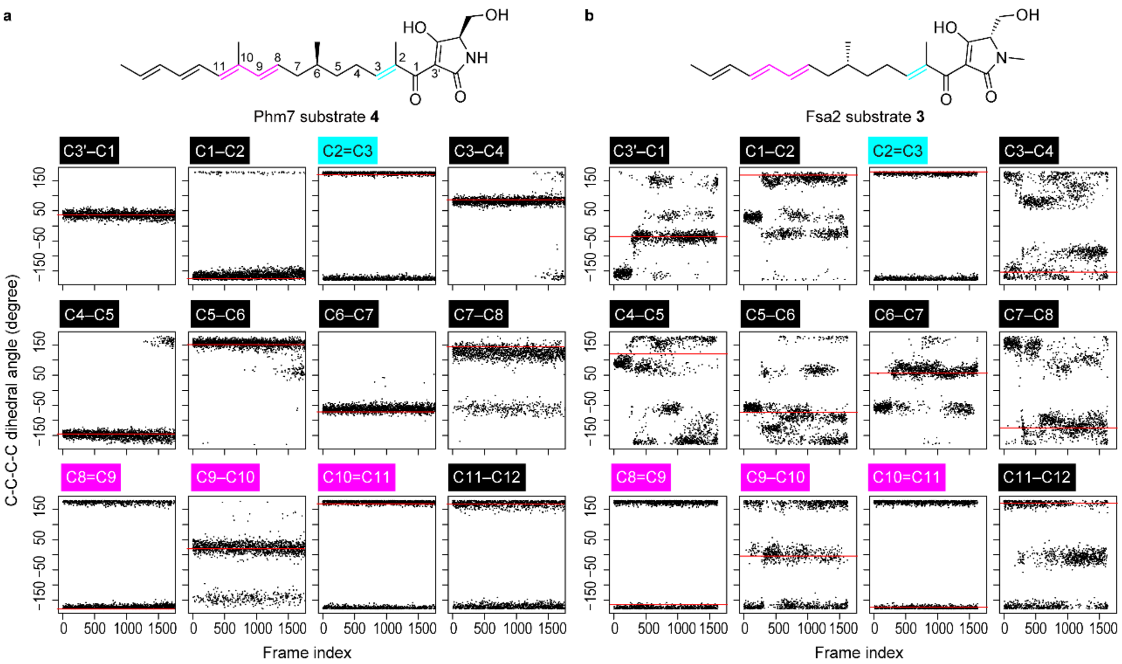


**Supplementary Fig. 12.** Conformations of the substrate **4** and **3** in the Phm7 and Fsa2 pockets. The C–C–C–C dihedral angles along the carbon chain of **4** (**a**) and **3** (**b**) were plotted for all of 3,169 and 1,629 poses in each "folded" cluster (pA and fA in Supplementary Fig. 9), respectively. The dihedral angles of the representative poses **4** and **3** are indicated by red bars. The representative pose of **3** is one that best amenable to the stereoselective DA reaction, and manually taken from the fA cluster (cf. Supplementary Fig. 26).

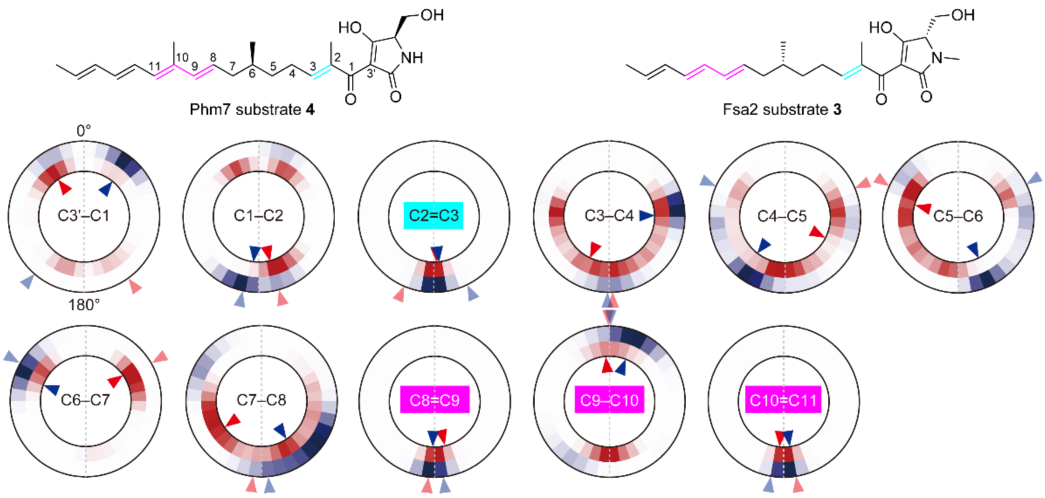


**Supplementary Fig. 13.** The distribution of the C–C–C–C dihedral angles along the carbon chain of the substrates **4** and **3** in the Phm7 and Fsa2 pockets. Circular heatmaps display the distributions of the dihedral angles of the substrates **4** (outer circle, indicated in blue) and **3** (inner, indicated in red), which were calculated from all the poses in the folded cluster. The dihedral angles of the representative poses **4** and **3** are indicated by blue and red arrow heads inside heatmaps, respectively. The angles of $\mathbf{TS}_{\mathbf{1b}}$ and $\mathbf{TS}_{\mathbf{1c}}$, which are corresponding to the decalin scaffolds of **2** and **1**, are also indicated by arrow heads outside heatmaps. Like bilaterally symmetric angles of the $\mathbf{TS}_{\mathbf{1b}}$ and $\mathbf{TS}_{\mathbf{1c}}$, all the angles of the representative poses of **4** and **3** are on either side, supporting the enantiomeric binding poses of the substrates in the enzyme pockets.

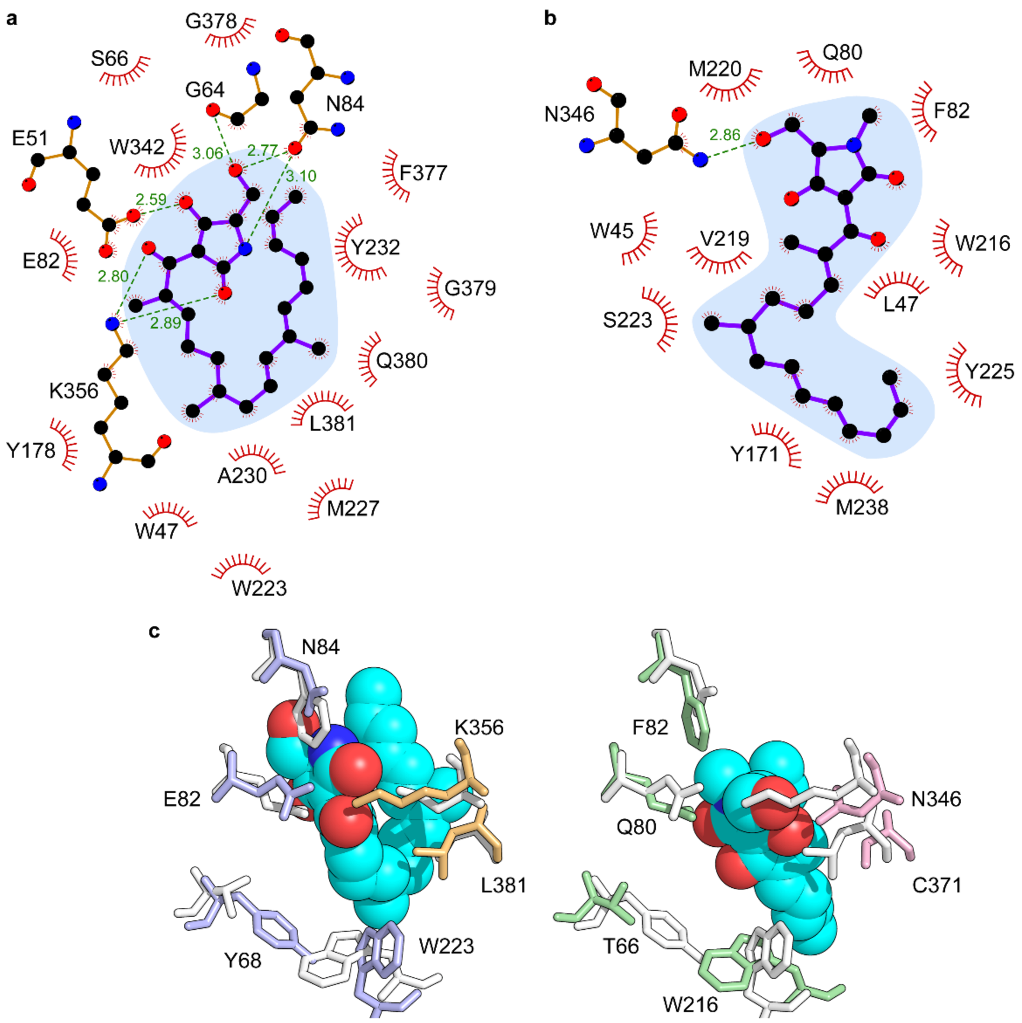


**Supplementary Fig. 14.** Substrate–enzyme interactions at bound states. **a**,**b**, Details of substrate–enzyme interactions were illustrated using Ligplot diagrams[8] for two representative poses of **4**•Phm7 (**a**) and **3**•Fsa2. (**b**) relevant to DA reaction. C, N, and O atoms are shown in ball representation with black, blue, and red colour. Hydrogen bond interactions are shown in green dashed line. Hydrophobic interactions are represented by a red arc with radiation toward the ligand. **c**, Comparison of the substrate–enzyme interactions between Phm7 (left) and Fsa2 (right) The substrate and amino acid residues are shown in sphere and stick representations, respectively. The amino acid residues are coloured according to the belonging domain. In Phm7 (Fsa2), the corresponding residues in Fsa2 (Phm7) are shown in white sticks for comparison. The bound substrates are commonly stabilized by many hydrophobic contacts. In addition, the tetramic acid of substrate **4** forms distinct hydrogen bonds with E51, G64, N84, and K356. These hydrogen bonds firmly capture the tetramic acid moiety of **4** to N-domain side, placing it to the upper part of the pocket. The residues from C-domain mainly contacts the polyene tail to stabilize the folded conformation. On the other hand, the tetramic acid of substrate **3** forms only one hydrogen bond with N346 from C-domain, placing it to the middle of pocket.

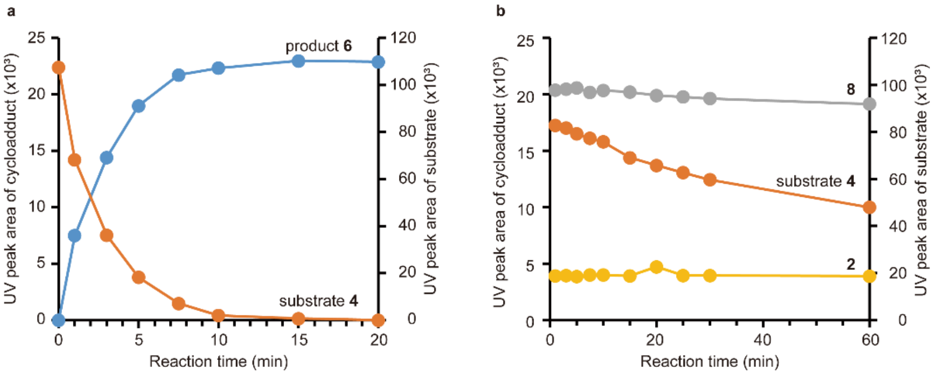


**Supplementary Fig. 15.** Time course of the *in vitro* Phm7 reaction. The cell lysates prepared from the Δ*phm7* mycelia were incubated with (**a**) and without Phm7 (**b**) at 25 °C, and the reactions were monitored by measuring UV peak areas of phomasetin (**2**), substrate **4**, Phm7 product **6**, and nonenzymatic cycloadduct **8**. **a**, In the presence of Phm7, linear formation of the product **6** with the first several minutes with complete consumption of **4** in 20 min was observed. **b**, On the other hand, **4** was gradually decreased by almost half within 60 min in the absence of enzyme. Neither **8** nor **2** was significantly changed during the reaction, and **6** was less than the detection limit. These results indicated that no cycloaddition proceeded in the absence of Phm7 under the conditions tested, and **2** and **8** detected in the reaction mixtures were present in the cell lysates from the beginning and likely to be formed during the culture.

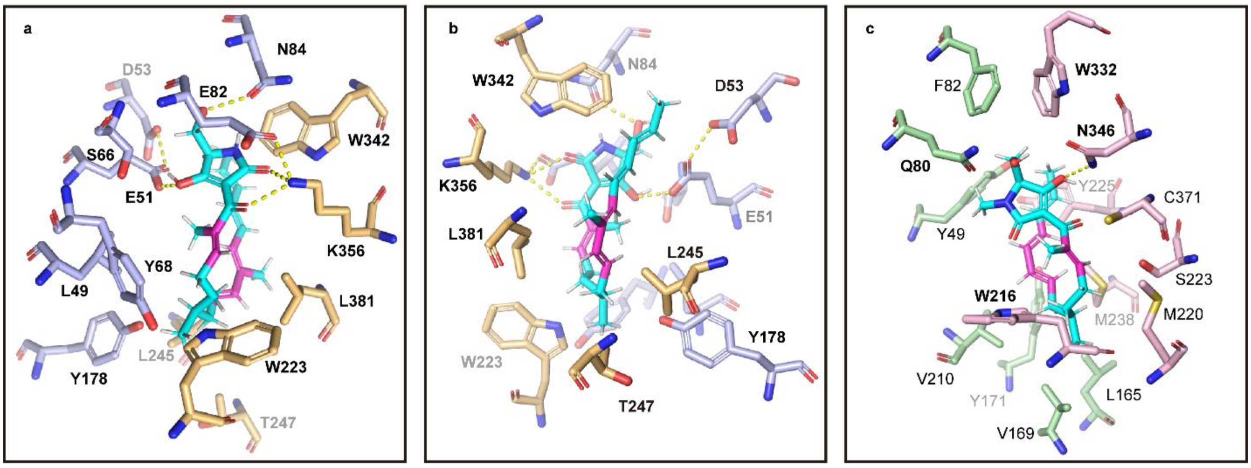


**Supplementary Fig. 16.** Location of the amino acid residues examined in this study. Based on the MD simulations, 14 amino acid residues of Phm7 (**a**,**b**) and 4 of Fsa2 (**c**, indicated in boldface) were selected for site-directed mutagenesis study.

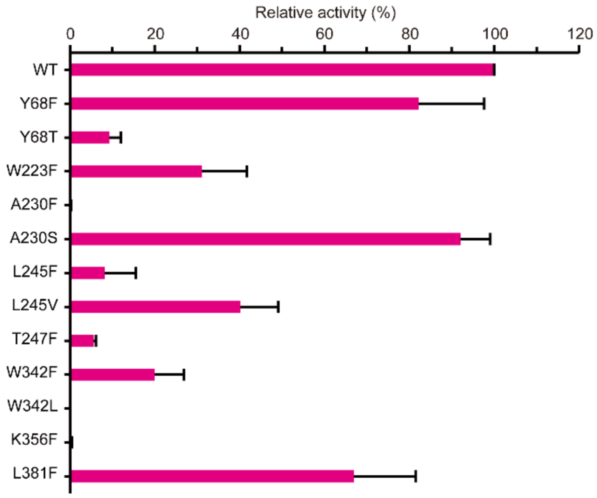


**Supplementary Fig. 17.** *In vitro* analysis of Phm7 mutants substituted with various amino acid residues. The enzyme activity of Phm7 mutants was compared with the wild-type enzyme (WT).

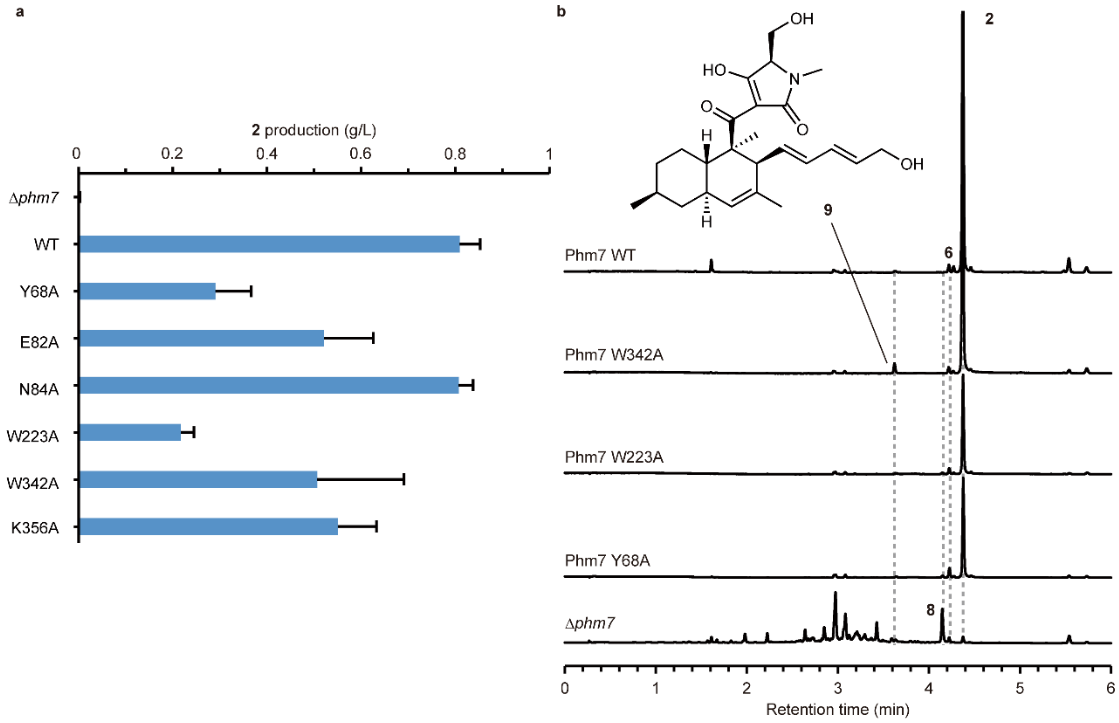


**Supplementary Fig. 18.** Evaluation of the Phm7 mutants in the producer fungus. **a**, Production of **2** in the genetically modified fungi carrying the mutated *phm7* that were derived from the **2**-producer fungus *Pyrenochaetopsis* sp. RK10-F058. The fungi were cultured in CYA medium at 28 °C for 9 days. The culture extracts were analysed by LC/MS, and amount of **2** was calculated with a standard curve[2]. Data are mean and error bars represent the standard deviation of three independent experiments. **b**, UPLC traces of the culture extracts of the genetically modified fungi containing the mutated *phm7*. A new peak **9** was detected in the culture extract of the W342A mutant. Structural analyses by NMR and MS revealed that **9** was a derivative of **2** with a hydroxy group at the terminus of polyene (See Supplementary Note for structure determination of **9**). Chemical structure of **9** is shown in an inset. Decreased turnover rate of the Phm7 W342A mutant could result in hydroxylation of the linear polyenoyl tetramic acid substrate **4** by unknown oxygenase(s) prior to the cycloaddition, which is possibly accepted to the Phm7 pocket. The replacement of W342 by Ala formed space in the pocket where the interaction between the enzyme and the polyene tail of **4** was likely to occur. The metabolite production in this gain-of-function mutant supported the role of W342 in the interaction with the polyene moiety of **4**.

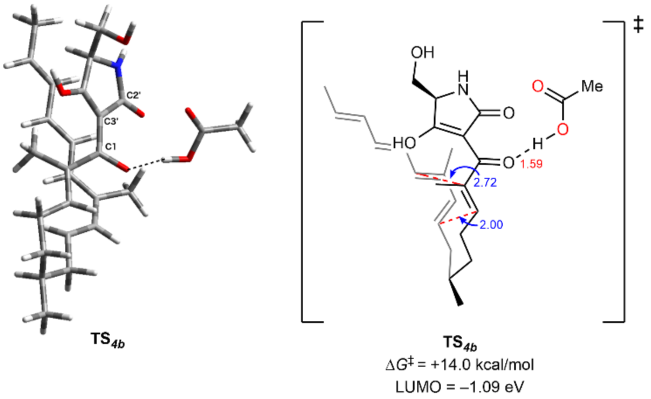


**Supplementary Fig. 19.** The DFT calculation for the DA reaction of **4** with an acetic acid having the hydrogen bonding with C1-carbonyl oxygen, as a model of hydrogen bonding of E82 residue in the Phm7 K356A mutant. A tight hydrogen bond between the carbonyl oxygen at C1 and acetic acid (1.59 Å) efficiently lowered the activation barrier and LUMO energy. These results suggest that the direct participation of E82 residue in the hydrogen bonding with **4** compensates the Ala substitution of K356, avoiding a complete loss of the activity.

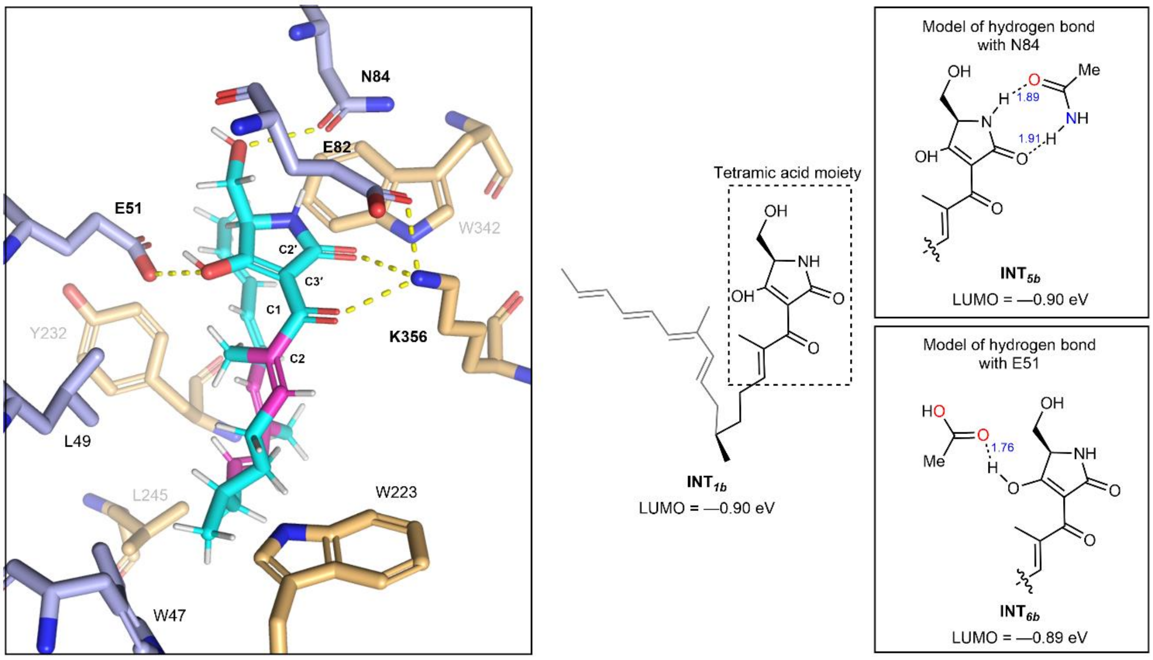


**Supplementary Fig. 20.** The DFT calculation for the investigation of the effects of amino acid residues as Brønsted acids or base on the LUMO energies of **4** coordinated with an acetamide as a model of N84 and acetic acid as a model of E51, respectively. Both models for the hydrogen bonds with N84 and E51 did not show the lowered LUMO energies, indicating that N84 and E51 residues do not accelerate the DA reaction of **4** and are likely to accommodate substrate **4** to the appropriate conformation in the pocket of Phm7.

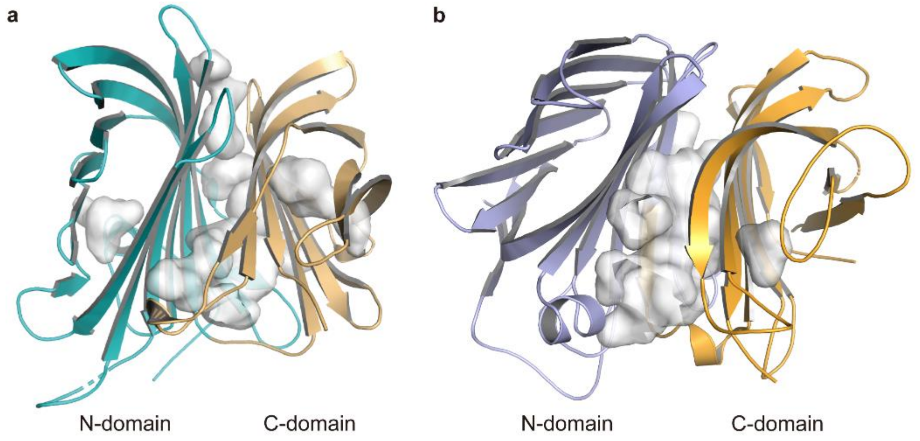


**Supplementary Fig. 21.** Comparison between a structurally related protein, NE1406, from *Nitrosomonas europaea* (2ICH, **a**)[9] and Phm7 (**b**). Structure of NE1406 is very similar to that of Phm7 (RMSD = 6.352 Å) although they showed low sequence identity of 14%. NE1406 consists of N and C-domains, which belong to PF07143 (CrtC N-terminal lipocalin domain) and PF17186 (Lipocalin_9), respectively. Bacterial homologs, belonging to the Pfam family PF171786, have been shown to have a carotenoid 1,2-hydratase activity[10], and their homology models have suggested that some catalytic residues are located at the pocket between the two domains[11].

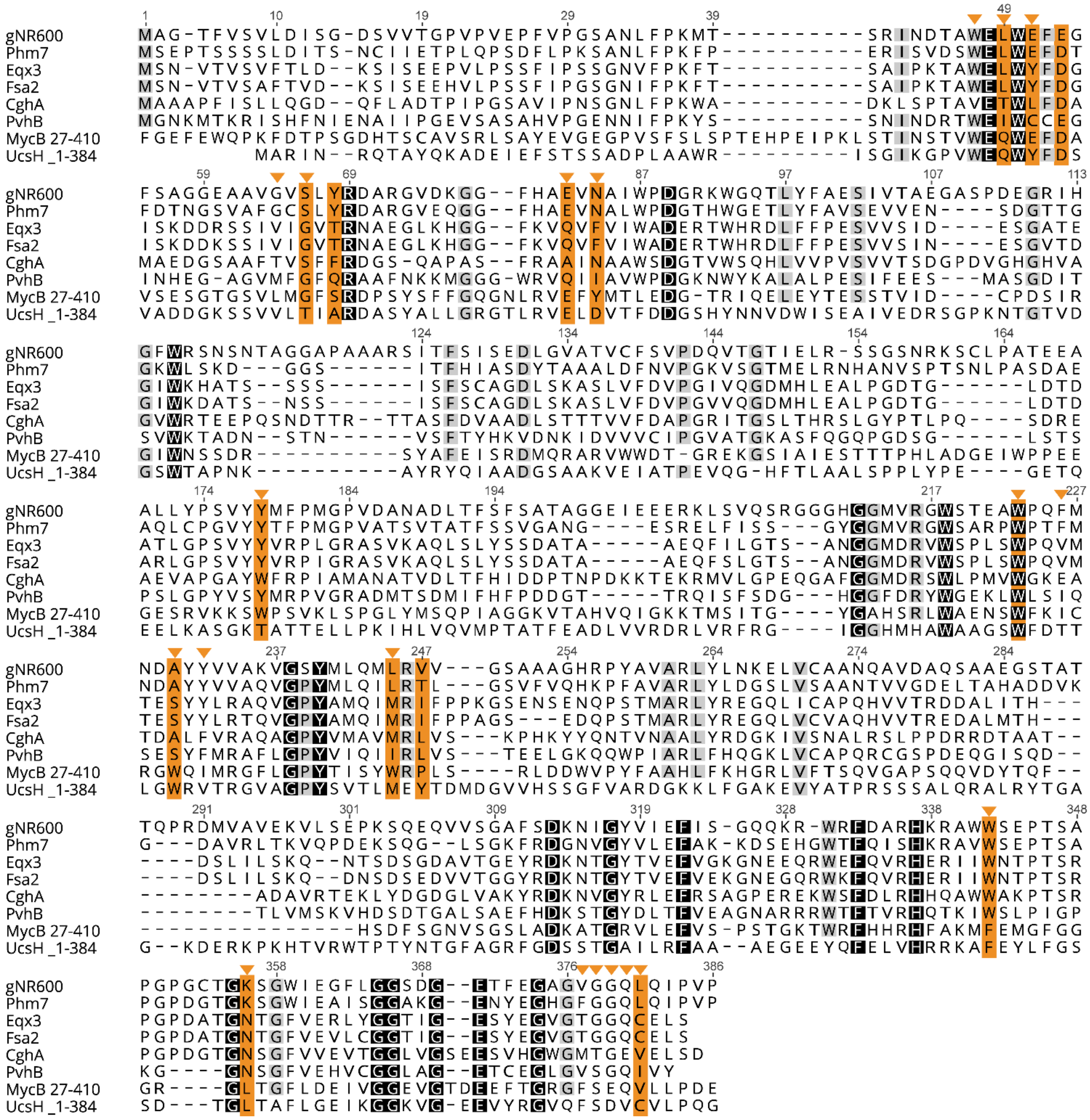


**Supplementary Fig. 22.** Multiple sequence alignment of Fsa2-family decalin synthases. Amino acid sequences of DSs whose functions have been characterized[2,12-17] were used for multiple alignment. Identical and conserved residues are shaded by black and grey, respectively. Positions in Phm7 are shown above the alignments. Amino acid residues predicted to be involved in the substrate interactions of either Phm7 or Fsa2 (cf. Supplementary Fig. 14) are indicated by arrowhead, and residues experimentally examined are highlighted by orange.

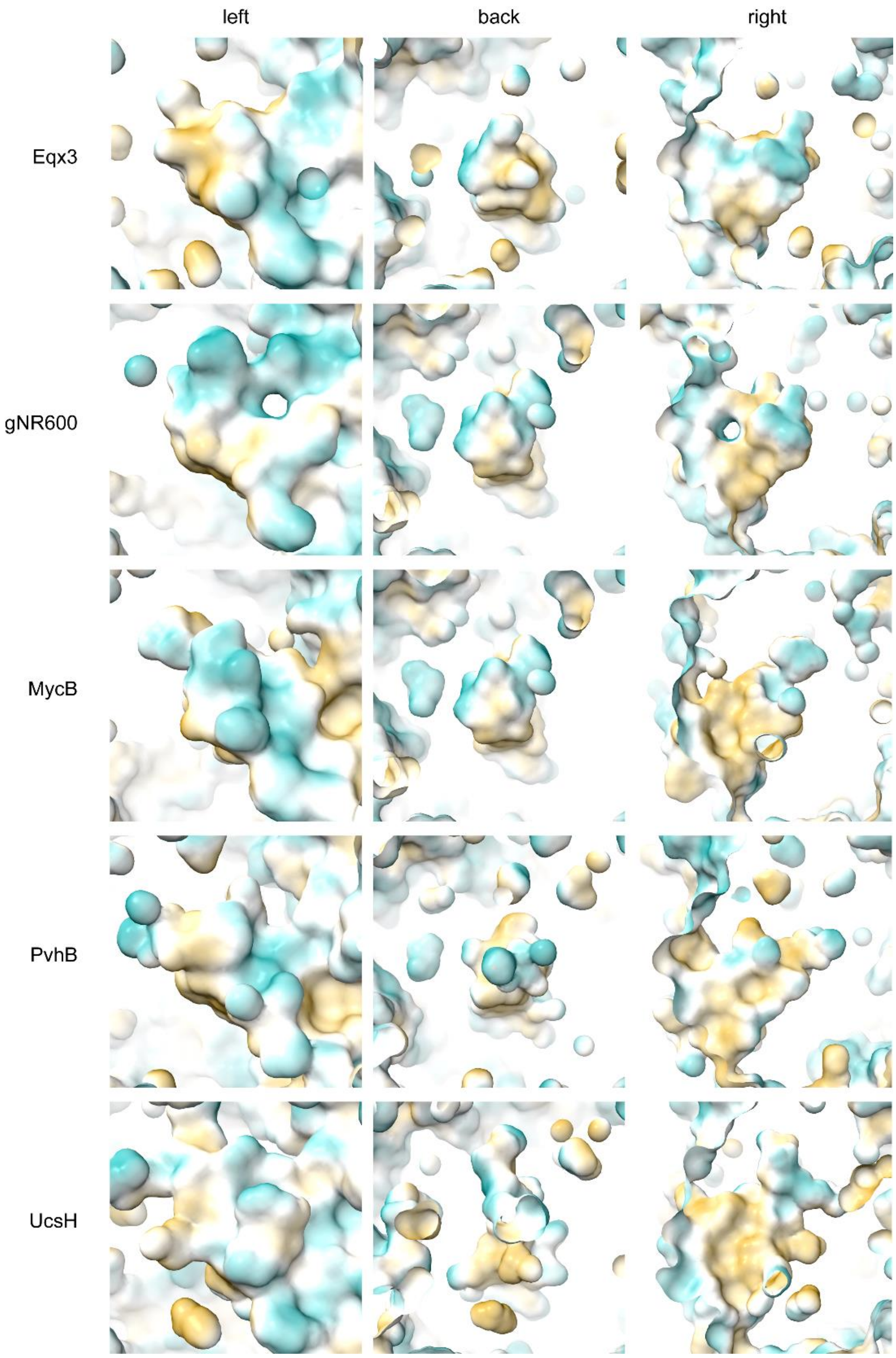


**Supplementary Fig. 23.** Surface models showing hydrophobicity maps of the substrate binding pocket in the homology models of Eqx3, gNR600, MycB, PvhB, and UcsH, which are built by SWISS-MODEL server by using the substrate-free Phm7 as a template. The 50 % transparent molecular surfaces are displayed with colours ranging from goldenrod (hydrophobic) through white (intermediate) to cyan (hydrophilic) according to the hydrophobicity map value.

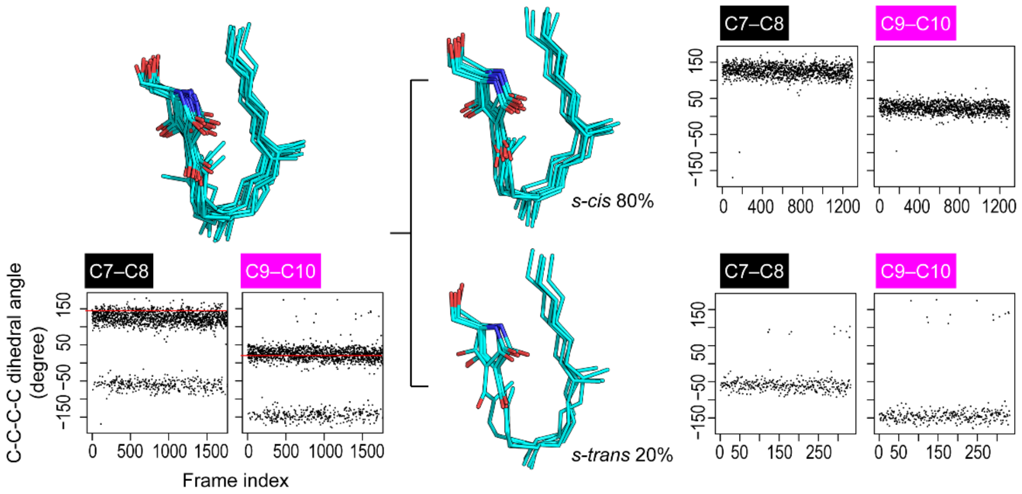


**Supplementary Fig. 24.** The convergent poses of **4** in the Phm7 pocket. A collective view of 12 representative snapshots taken from top 70% of the cluster members (RMSD from the cluster centre in ascending order) of pA, which were further divided according to the diene orientation (C9–C10) of either *s-cis* or *s-trans*. The rotation at the C9–C10 bond is highly correlated with that at the C7–C8 bond. The relative population of each conformation is also shown.

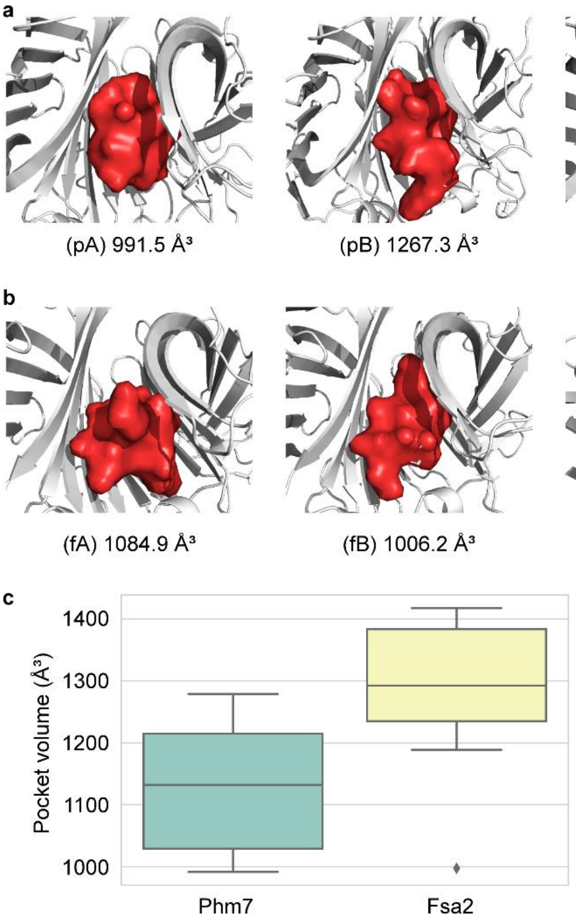


**Supplementary Fig. 25.** Size and shape of binding pockets. **a**,**b**, The binding pockets calculated for the representative structure of each cluster in **4**•Phm7 (**a**) and **3**•Fsa2 (**b**) using PyVOL plugin for PyMOL[18,19]. The *largest* mode with default settings (minimum volume of 200, minimum and maximum radii of 1.4 and 3.4, respectively.) were used for calculations. The pockets are shown in red surface representation. The calculated pocket volumes are also shown. **c**, Boxplots of pocket volumes calculated for 12 representative snapshots taken from top 70% of the cluster members of pA and fA (Fig. 3a,b). The average volume is 1129.6±105.1 and 1284.2±119.9 Å$^3$ for Phm7 and Fsa2, respectively. Mean difference is 154.6 Å$^3$ with *p*-value of 0.003, ensuring that the calculated difference in cavity volumes is statistically significant.

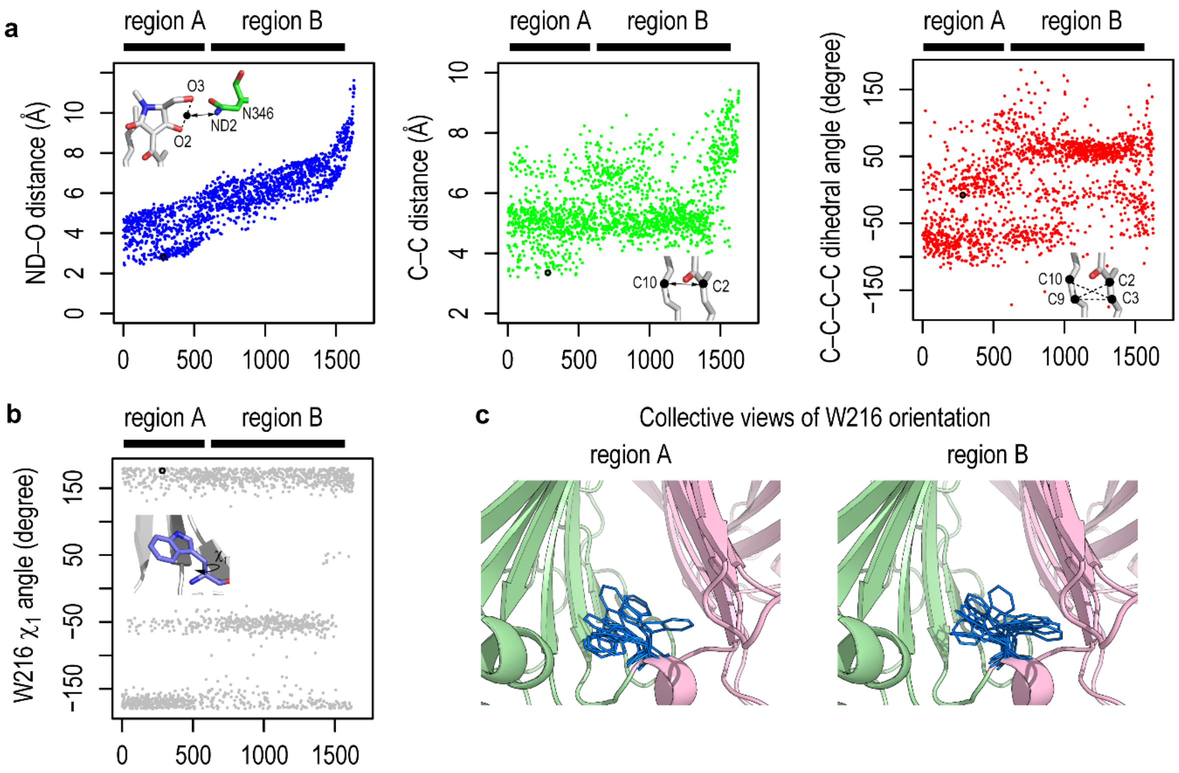


**Supplementary Fig. 26.** Conformational changes in the major cluster of **3** (fA) along the tetramic acid–N346 (N–O) distance. **a**, tetramic acid–N346 (N–O) distance, C–C distance between the diene and dienophile, and C–C–C–C dihedral representing the relative orientation of the diene and dienophile. The corresponding values of the representative structure (Fig. 3b in the main text) are shown in black circle. As the tetramic acid and N346 get closer, the distance between diene and dienophile becomes shorter (region A). **b**, $\chi_1$ angle of W216 along the tetramic acid–N346 distance. The $\chi_1$ values exhibit two distribution for distant tetramic acid–N346 distances, while one of distributions becomes dominant in the short tetramic acid–N346 distances. **c**, Ten collective W216 structures are shown for short (region A, left) and distant (region B, right) tetramic acid–N346 distances.

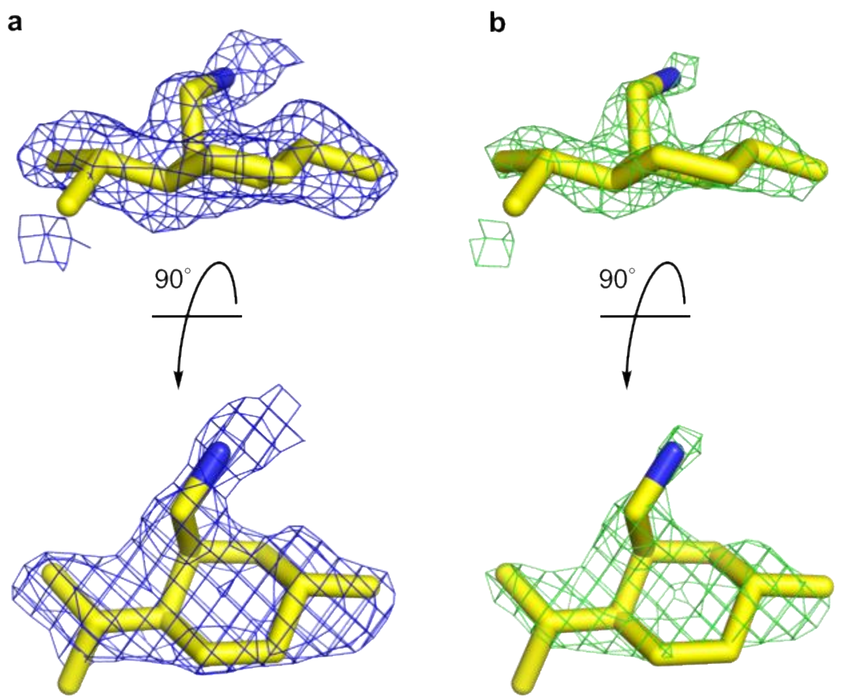


**Supplementary Fig. 27.** A polder map of inhibitor **5** in the Phm7 substrate binding site. Blue (**a**) and green (**b**) meshes show 2.0 and 3.0 σ, respectively.

### SUPPLEMENTARY NOTES

#### Molecular dynamics (MD) simulations.

The initial configurations of enzyme-substrate complexes, **4**•Phm7 and **3**•Fsa2, were built based on the results of precedent docking simulations. After manual adjustments, each system was neutralized and solvated in 150 mM NaCl solution. The final systems contain approximately 65,000 atoms (~ 20,000 $H_2O$ molecules) with box dimensions of 93×81×90 Å$^3$ and 97×79×92 Å$^3$ for **4**•Phm7 and **3**•Fsa2, respectively (Supplementary Fig. 28). All simulations were performed using the GENESIS program package (version 1.4.0)[20,21]. The AMBER ff14SB force field[22] was used for protein and ions, while the TIP3P model[23] was used for water molecules. The substrates were parameterized with the general AMBER force field parameter set version 2.1 (GAFF2) and AM1-BCC atomic charges using the antechamber module in Amber Tools 18[24,25]. The SHAKE and SETTLE algorithms[26,27] were used to constrain the covalent bonds involving hydrogen atoms and to keep water molecules rigid. Particle-mesh Ewald (PME) summation was used for long-range electrostatic interactions[28,29], while the non-bonded interactions were truncated at 8 Å with dispersion correction to account for the effect of long-range van der Waals interactions. The r-RESPA integrator was used with a time step of 2.5 fs[30], where PME calculation was performed every 2 steps. Bussi thermostat (300 K) and/or barostat (1 atm) were used for all simulations[31,32]. Each system was first minimized, and gradually heated to 300 K with the constraints applied on both the protein backbone and ligand atoms. The systems were then briefly relaxed in the NPT ensemble by gradually removing the restraints. The final configurations were used for the gREST simulations.

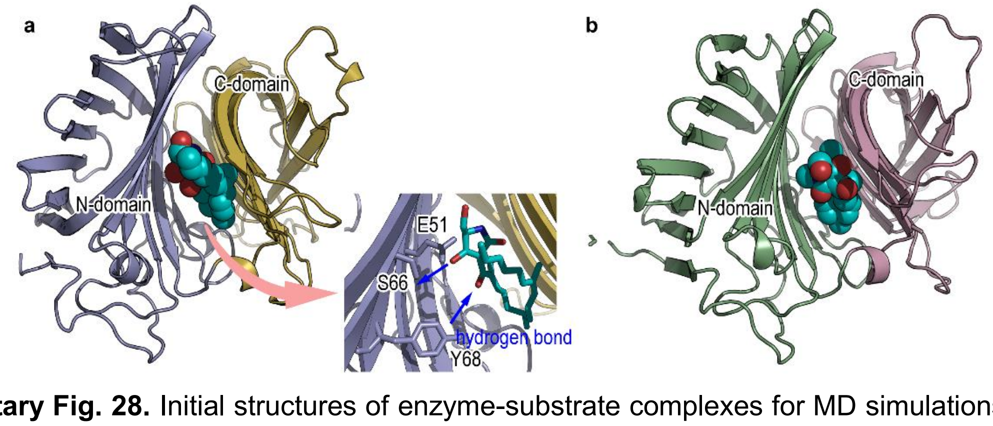


**Supplementary Fig. 28.** Initial structures of enzyme-substrate complexes for MD simulations. The initial bound forms of **4**•Phm7 (**a**) and **3**•Fsa2 (**b**) based on the docking simulations. Enzyme and ligands are show in cartoon and sphere representations, respectively. The tetramic acid moiety in **4**•Phm7 was manually reoriented to allow hydrogen bonding with polar residues in N-domain for transition state stabilization by the enzyme. The missing loop (disordered region of residues D282-K292) of Phm7 was complemented using Loop modeler tool implemented in MODELLER program[33]. D53 of Phm7 was protonated according to the p$K_a$ value (10.6) estimated from PROPKA3[34,35].

**gREST simulations.**

In the protein-ligand binding pose prediction, the sampling of all possible states is often hampered by high energy barriers associated with ligand re-orientation. Replica-exchange MD (REMD)[36], which is one of the simplest and most widely used enhanced sampling methods, uses multiple copies of the system (replicas) and exchanges the temperatures between neighbouring replicas to effectively overcome the high energy barriers. The generalized replica exchange with solute tempering (gREST)[37], which is a generalized version of REST or REST2[38-40], performs the replica exchange within a predefined subspace called as "solute" (a part of the system and/or a part of potential energy terms). The method has been successfully applied to predict protein-ligand binding poses[41,42]. In this study, *w*e defined the solute region as the dihedral angle and non-bonded energy (Coulomb and Lennard-Jones) terms of both the substrate and a set of selected binding site residues: D53, S66, Y68, E82, W223, W342, and K356 for Phm7 and the corresponding residues of Fsa2 (D51, G64, T66, Q80, W216, W332, and N346). Eight replicas were employed to cover the solute temperature range of 310.0–773.9 K (T = 310.0, 350.8, 396.7, 452.8, 519.6, 592.3, 675.0, and 773.9 K). These values were derived using an automatic tuning algorithm in GENESIS such that the resultant parameters achieve an exchange probability of 0.25. For further efficiency, we also applied a flat-bottom potential to avoid the ligand being away from the binding site.

$$U(R) = \begin{cases} K(R - R_0)^2, R > R_0 \\ 0 \qquad\qquad\quad , R \leq R_0 \end{cases}$$

where $R$, $R_0$, and $K$ are the distances between the centre of masses of the substrate and binding site residues in the solute region, the flat-bottom distance, and the force constant, respectively. We set $R_0$ to 15.0 Å and $K$ to 1 kcal/mol/Å$^2$. The substrate feels no extra force in the region of $R$ = 0–15 Å, while the harmonic restraint potential is applied beyond $R$ = 15 Å to avoid substrate dissociation. Our models for the gREST simulations are illustrated in Supplementary Fig. 29. Each of the **4**•Phm7 and **3**•Fsa2 systems was initially relaxed at different temperatures for 1 ns followed by production runs for 100 ns per replica (total sampling of 0.8 μs = 100 ns × 8 replicas). We confirmed that the replica exchanges occur sufficiently, assuring the reliability of the current gREST simulations (Supplementary Fig. 30).

**Extraction of bound poses by clustering analysis.**

The representative bound poses were obtained through a clustering analysis on the simulation trajectories at 310 K. For the clustering, pairwise heavy atoms root-mean-square deviations (RMSDs) between the bound structures (the substrate and binding site residues) were calculated over 5000 snapshots (every 20 ps) from the trajectory. Clustering was performed using the density-based spatial clustering of applications with noise (DBSCAN) implemented in Scikit-learn[7,43]. We set the minimum size of the cluster to 1 % of the examined conformations (50 snapshots) and the spatial resolution to 5 Å.

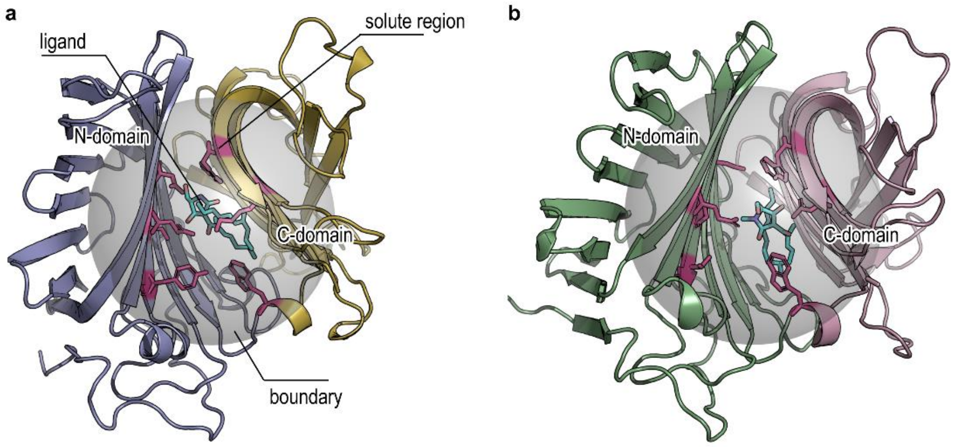


**Supplementary Fig. 29.** Models for gREST of **4**•Phm7 (**a**) and **3**•Fsa2 (**b**). N- and C-domains comprising the binding site are shown in cartoon representation. The solute region involves both selected binding site residues (sticks with magenta) and the substrate atoms (sticks with cyan). A grey sphere represents the boundary of the flat-bottom potential used to prevent the substrate dissociation from the binding site.

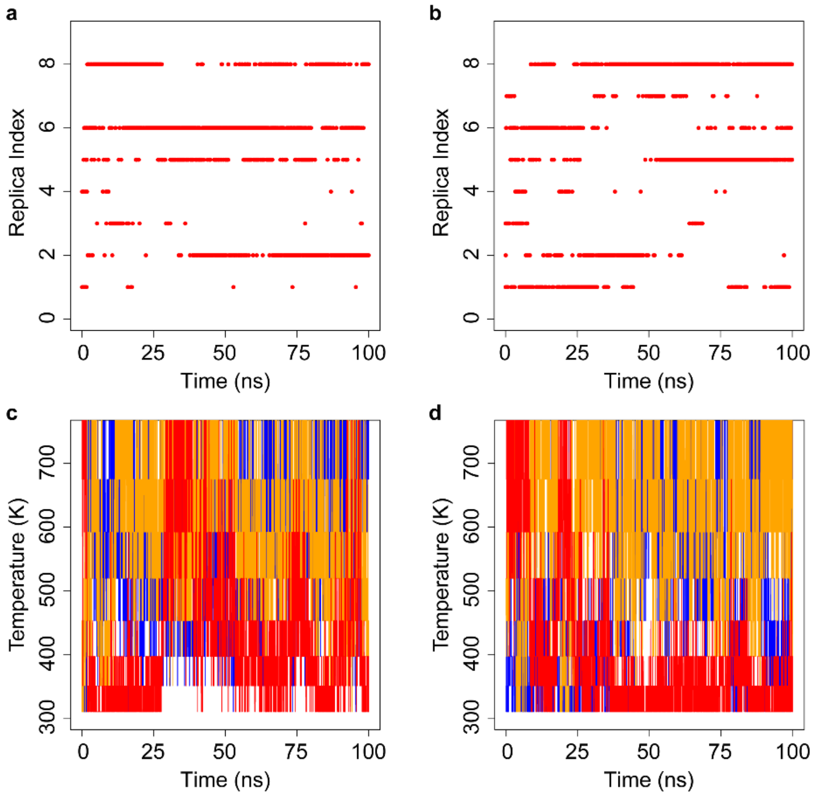


**Supplementary Fig. 30.** Replica and temperature exchanges. The time courses of the replica exchange at 310 K for **4**•Phm7 (**a**) and **3**•Fsa2 (**b**), and those of solute temperature exchanges of three arbitrary chosen replicas for **4**•Phm7 (**c**) and **3**•Fsa2 (**d**). The temperatures of three replicas are shown in blue, orange, and red. The acceptance ratios of the neighbouring replica exchanges are 24.1 % and 26.5 % for **4**•Phm7 and **3**•Fsa2, respectively.

**Purification and structure determination of 17-hydroxyphomasetin (9).**

The fungal strain containing the Phm7 K356A mutation was cultured at 28 °C for 9 days in CYA medium. The culture broth (10 L) was extracted three times with a half volume of EtOAc. It was evaporated to yield 2.23 g of crude extract and then subjected to $SiO_2$-MPLC with a $CHCl_3$/MeOH stepwise gradient to obtain seven fractions. MPLC fraction 6 was separated by C18-MPLC with a gradient elution of acetonitrile/water to yield the compound **9**-rich fraction. It was purified by C18-HPLC with acetonitrile/0.05 % v/v aqueous formic acid (55:45) to afford 3.8 mg of **9** as a colourless amorphous solid.

The chemical structure of **9** was determined to be 17-hydroxyphomasetin by NMR and MS measurements (Supplementary Figs. 31–39 and Table 3). The molecular formula was determined to be $C_{25}H_{35}NO_5$ by HRESI-TOFMS analysis (found *m*/*z*: 430.2591 $[M+H]^+$ calcd. for $C_{25}H_{36}NO_5$, 430.2588). The planar structure was determined as an analogue of **2** based on 2D NMR correlations in DQF-COSY (Supplementary Fig 34), HSQC (Supplementary Fig 35), and HMBC (Supplementary Fig 36) spectra and the comparison of NMR chemical shift values with those of **2**[2]. The geometries of $\Delta^{13}$ and $\Delta^{15}$ were assigned as *E* by NOESY correlations of H13/H15 and H14/H16, respectively. NOESY correlations of H-6/H-8, H-6/H-10ax, H-6/Me-12, and H-3/Me-12 (Supplementary Fig 37) confirmed that **9** had the same relative stereochemistry as that of **2**. The absolute configuration of **9** was deduced to be 2*R*,3S,6R,8S,11S,5'R by comparing specific rotation values and ECD spectra with those of **2**[2] (Supplementary Fig 39).

**17-hydroxyphomasetin (9)**: colourless amorphous; $[\alpha]_{589}^{26}$ +418 (*c* 0.1); UV (MeOH) $\lambda_{max}$ (log $\varepsilon$) 237 (4.50), 293 (3.93) nm; IR (ATR) $\nu_{max}$ 3315, 1646, 1596, 1446, 1375, 1224, 1079, 987 $cm^{-1}$; $^1H$ and $^{13}C$ NMR ($CD_3OD$), summarized in Supplementary Table 3; ESIMS (*m*/*z*) 430 $[M+H]^+$; HRESITOFMS (*m*/*z*) 430.2591 $[M+H]^+$ (calcd. for $C_{25}H_{36}NO_5$, 430.2588).

**Supplementary Table 3.** $^{1}H$ and $^{13}C$ NMR chemical shifts of **9** in in $CD_3OD$

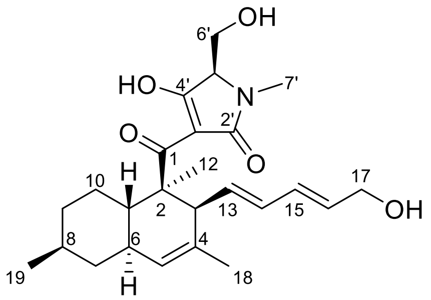

| **position** | | **δC** | **δH** | **Multi (*J* in Hz)** | **position** | **δC** | **δH** | **Multi (*J* in Hz)** |
|---|---|---|---|---|---|---|---|---|
| 1 | | 204.7 | - | | 13 | 132.3 | 6.08 | m |
| 2 | | 50.7 | - | | 14 | 133.1 | 5.87 | dd (14.9, 10.3) |
| 3 | | 50.5 | 3.40 | m | 15 | 130.6 | 5.38 | m |
| 4 | | ca. 130* | - | | 16 | 132.3 | 5.62 | m |
| 5 | | 127.7 | 5.22 | m | 17 | 63.6 | 3.75 | m |
| 6 | | 40.8 | 1.85 | m | | | 4.01 | m |
| 7 | ax | 44.1 | 0.87 | m | 18 | 22.8 | 1.59 | 3H, s |
| | eq | | 1.82 | m | 19 | 23.2 | 0.93 | 3H, d (6.3) |
| 8 | | 35.1 | 1.52 | m | 2' | 174.1 | - | |
| 9 | ax | 37.3 | 1.10 | m | 3' | 110.2 | - | |
| | eq | | 1.75 | m | 4' | 196.7 | - | |
| 10 | ax | 29.5 | 1.03 | m | 5' | 67.0 | 3.42 | m |
| | eq | | 1.98 | m | 6' | 60.5 | 3.87 | brd (10.3) |
| 11 | | 41.5 | 1.71 | m | | | 3.93 | brd (10.3) |
| 12 | | 14.9 | 1.41 | 3H, brs | N-Me | 27.6 | 3.01 | 3H, s |

*confirmed by HMBC spectrum.

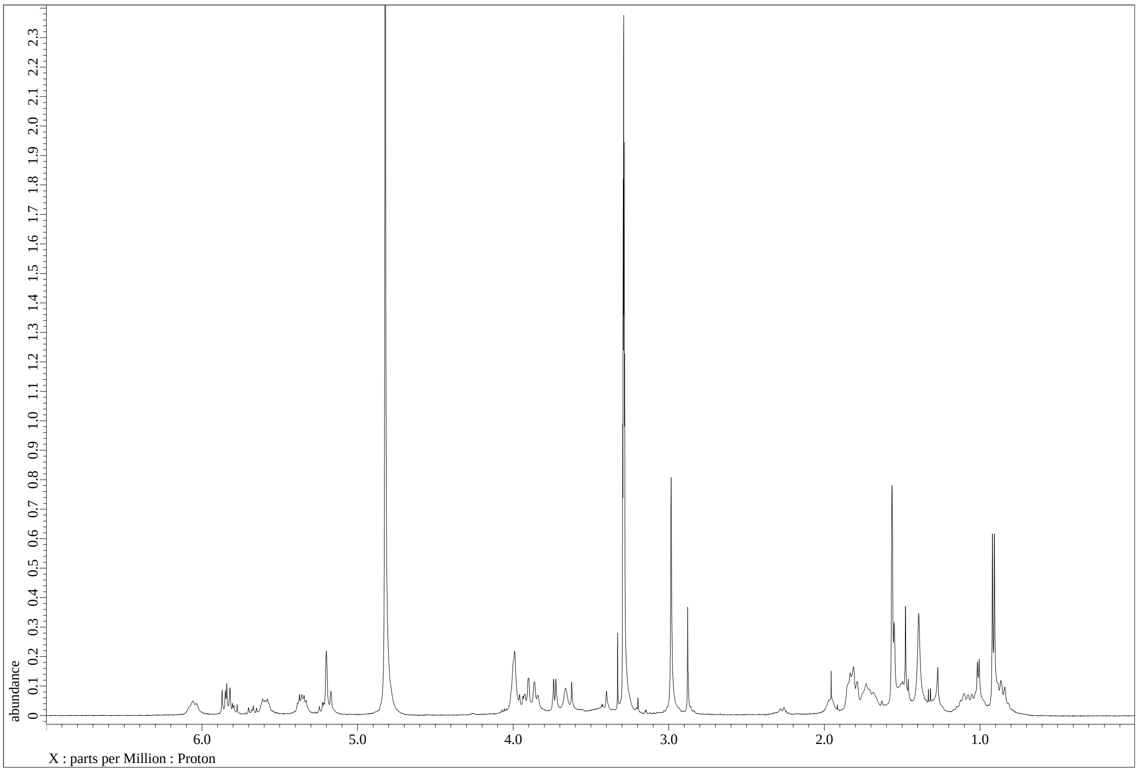


**Supplementary Fig. 31.** $^{1}H$ NMR spectrum of **9** in $CD_3OD$

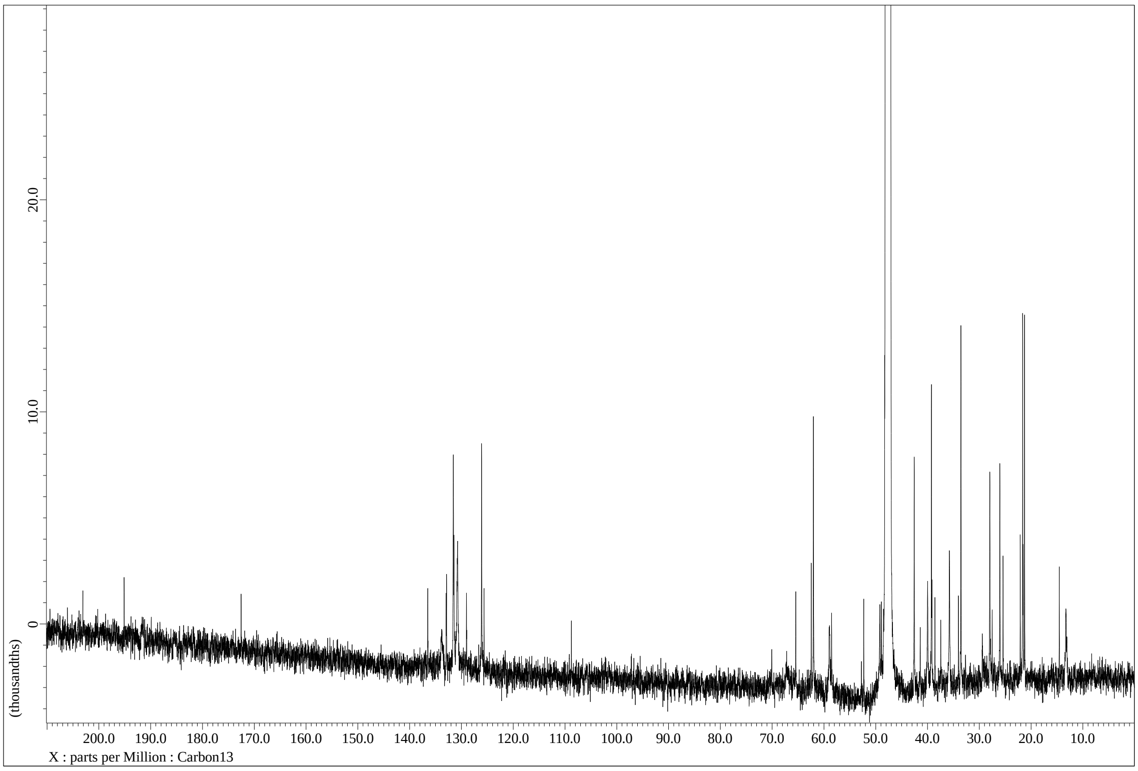


**Supplementary Fig. 32.** $^{13}C$ NMR spectrum of **9** in $CD_3OD$

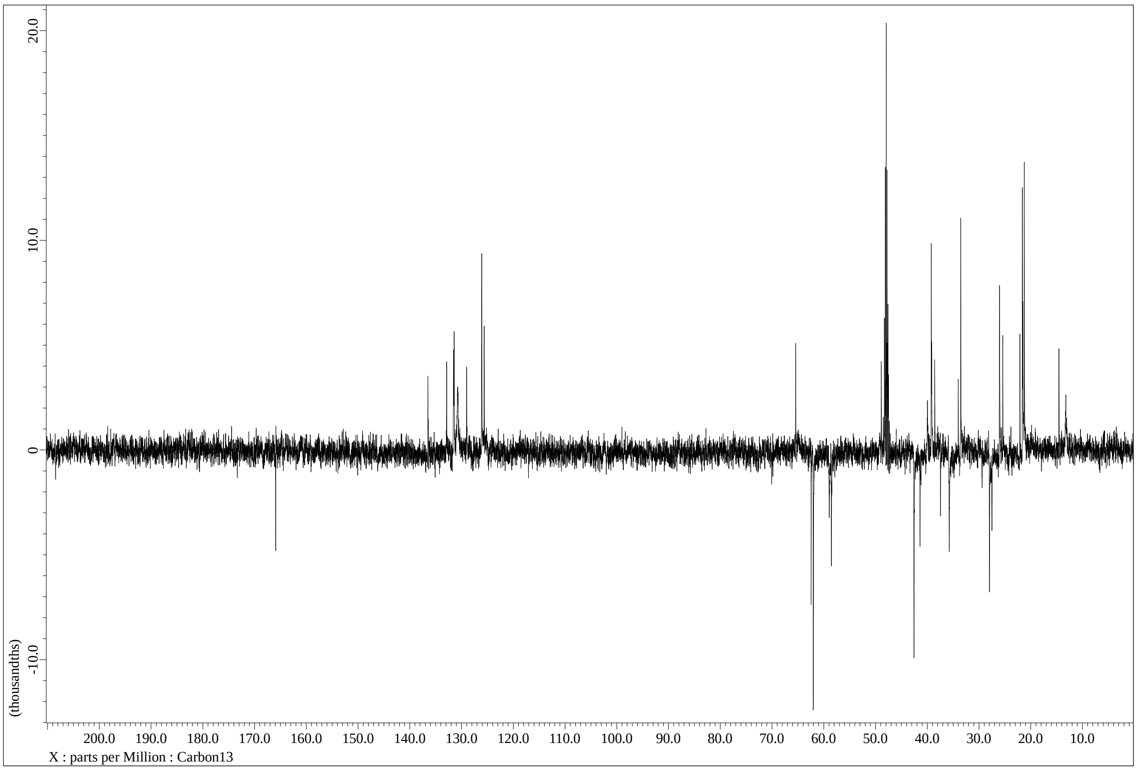


**Supplementary Fig. 33.** $^{13}C$ DEPT135 spectrum of **9** in $CD_3OD$

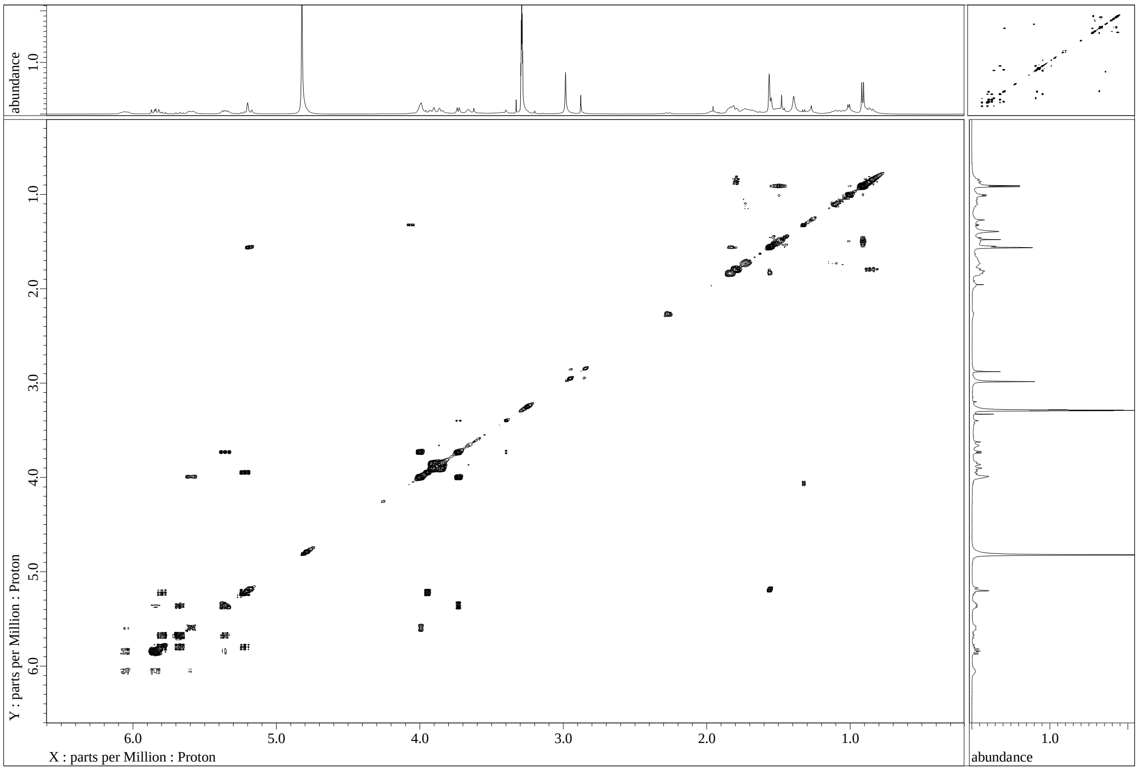


**Supplementary Fig. 34.** DQF-COSY spectrum of **9** in $CD_3OD$

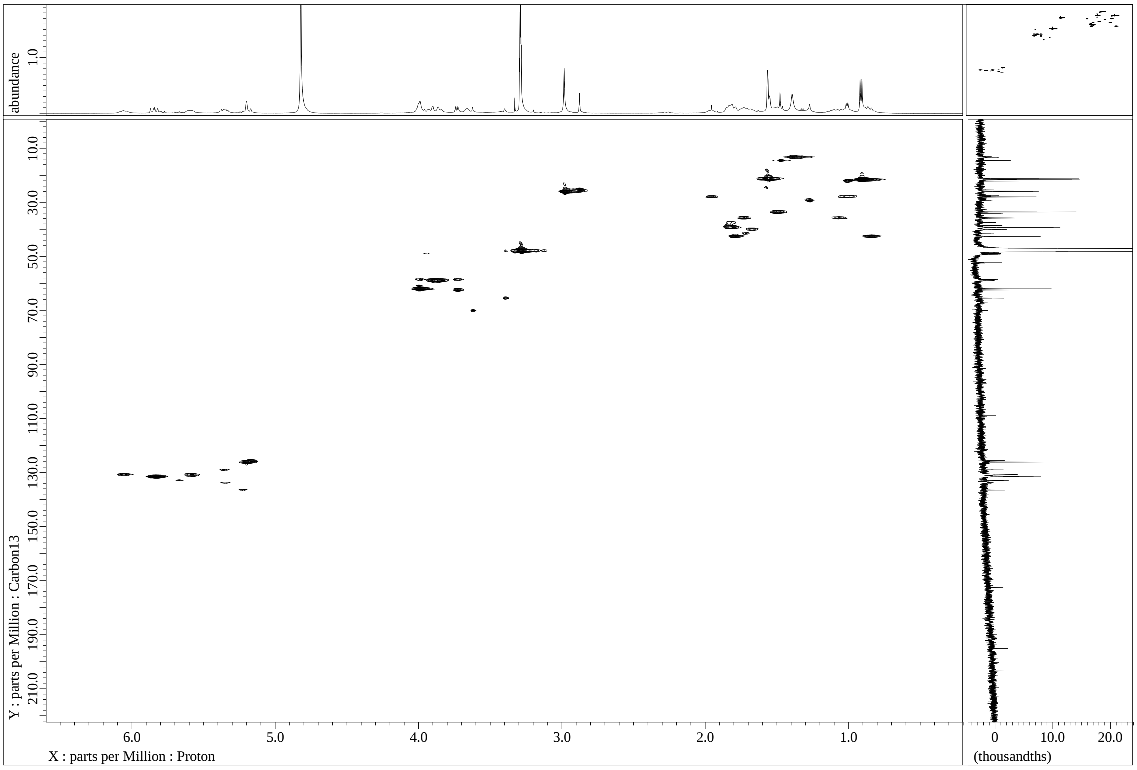


**Supplementary Fig. 35.** HSQC spectrum of **9** in $CD_3OD$

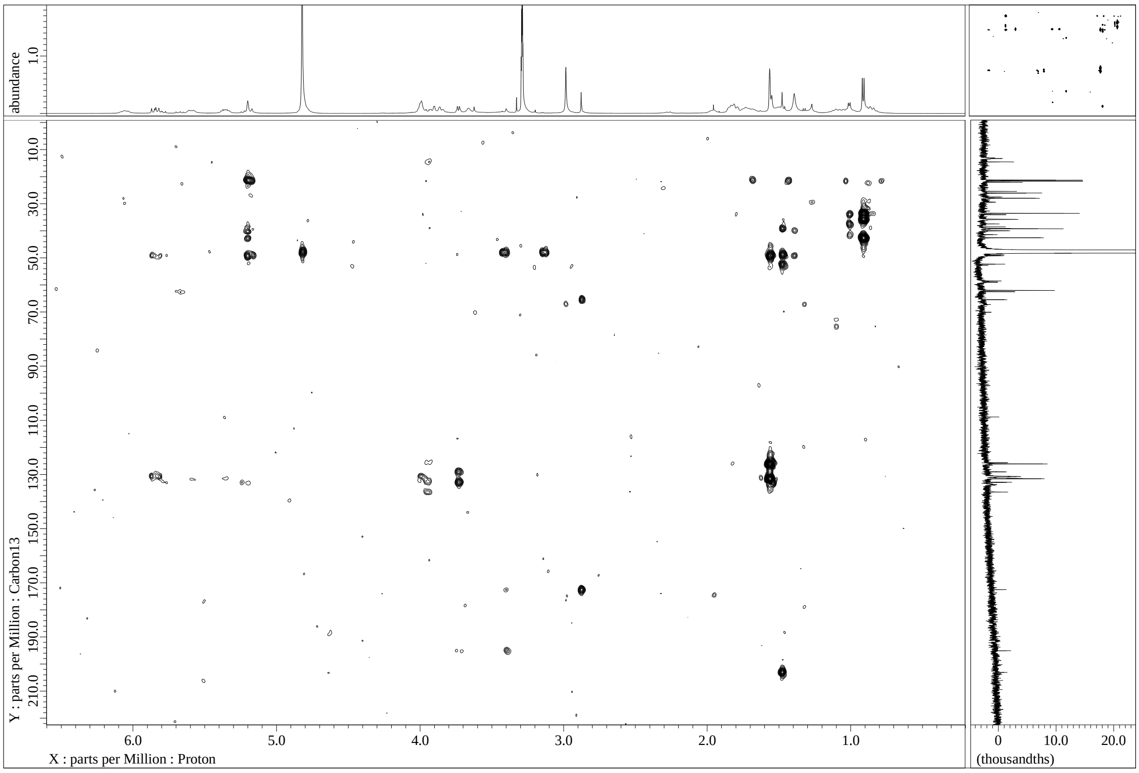


**Supplementary Fig. 36.** HMBC spectrum of **9** in $CD_3OD$

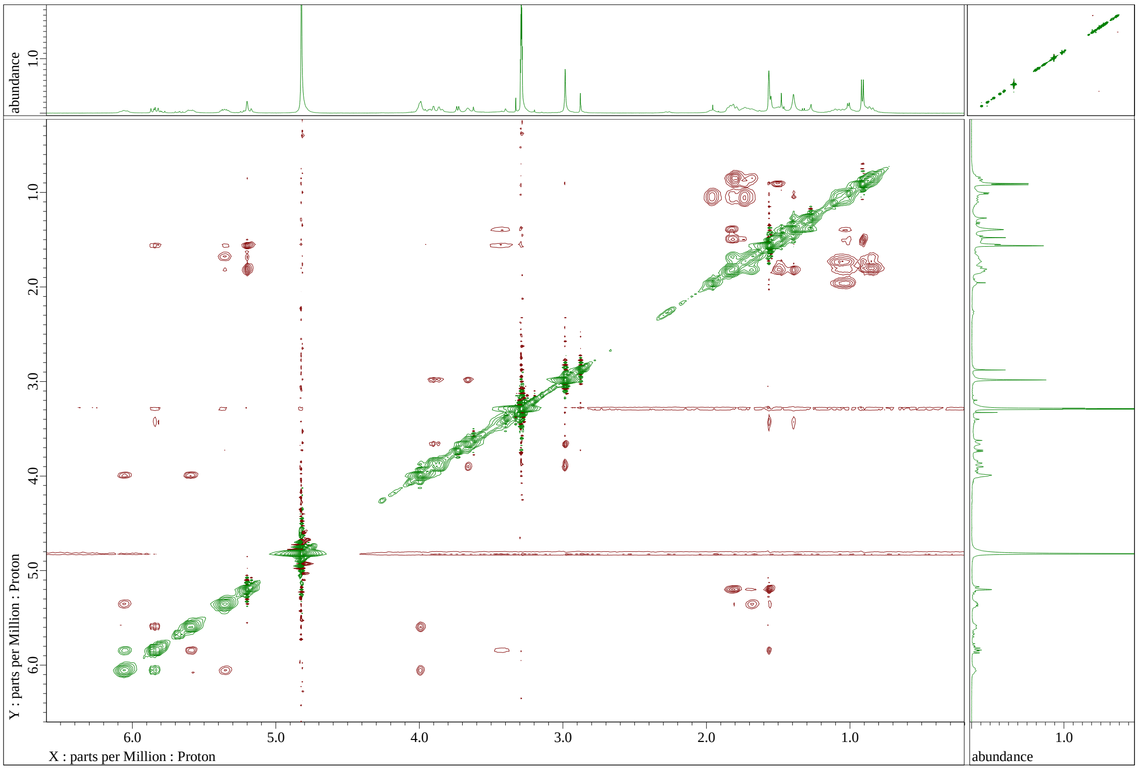


**Supplementary Fig. 37.** NOESY spectrum of **9** in $CD_3OD$

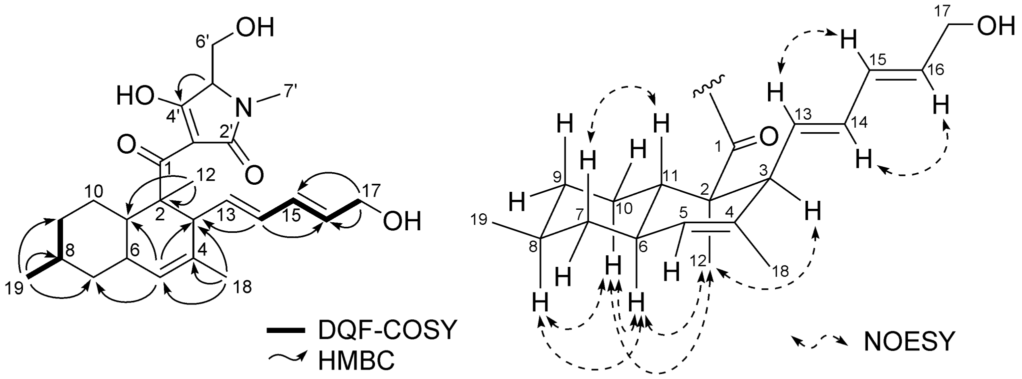


**Supplementary Fig. 38.** Key 2D NMR correlations of **9**

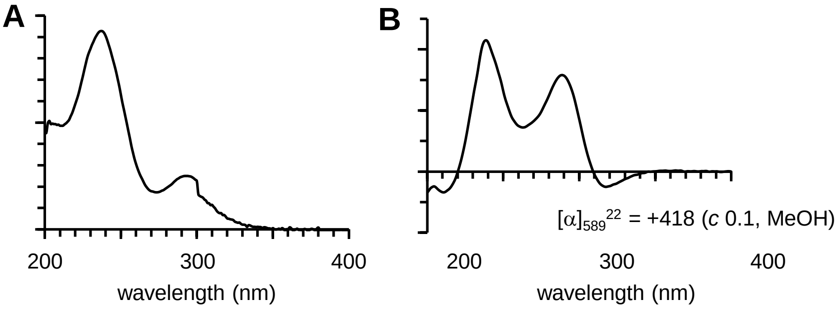


**Supplementary Fig. 39.** UV (A) and ECD (B) spectra of **9**

### Cartesian Coordinates and Energies

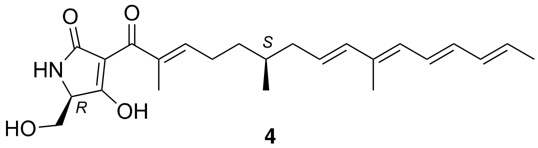


**4**

Energy (RM062X): -1289.827615 A.U.
Gibbs Free Energy: -1289.367069 A.U.

------------------------------------------------------------

| | | | |
|---|---|---|---|
| C | -0.58950300 | -1.64859600 | 1.19338400 |
| H | -0.87661400 | -0.60069100 | 1.05101000 |
| C | -1.30547400 | -2.50044500 | 0.12891700 |
| H | -1.08499200 | -3.55737600 | 0.29172900 |
| H | -0.93097700 | -2.24110100 | -0.86474800 |
| C | -2.78955100 | -2.28975500 | 0.19310500 |
| H | -3.37267100 | -3.01655800 | 0.75300000 |
| C | -3.44686500 | -1.24282800 | -0.32348800 |
| C | -2.80228700 | -0.12157100 | -1.09788200 |
| H | -2.63957600 | 0.75521100 | -0.46555400 |
| H | -1.83252400 | -0.42632000 | -1.49108500 |
| H | -3.42957200 | 0.17979500 | -1.94029400 |
| C | -4.92168700 | -1.18419000 | -0.08887500 |
| O | -5.61260300 | -2.18437300 | -0.02350400 |
| C | -5.54338600 | 0.15339200 | 0.06935200 |
| C | -5.10512200 | 1.19286200 | 0.80566100 |
| C | -6.87569300 | 0.52861400 | -0.47711800 |
| O | -3.97307000 | 1.24096600 | 1.50127700 |
| C | -6.06663300 | 2.35277500 | 0.75178400 |
| O | -7.57920500 | -0.10741500 | -1.24028200 |
| N | -7.17719900 | 1.76216200 | 0.03868900 |
| H | -3.90347900 | 2.05158700 | 2.02191300 |
| H | -6.34994100 | 2.66392500 | 1.76327100 |
| C | -5.46527800 | 3.56376600 | 0.01303200 |
| H | -7.89572600 | 2.32898000 | -0.38948100 |
| H | -6.22317400 | 4.34665700 | -0.02775700 |
| H | -4.61342900 | 3.95342600 | 0.57959900 |
| O | -5.11566500 | 3.26236300 | -1.31839100 |
| H | -4.28657700 | 2.77404600 | -1.33528300 |
| C | 0.94158300 | -1.75268100 | 1.17750900 |
| C | 1.53199400 | -1.20898000 | -0.13698900 |
| C | 1.42967600 | -3.17533900 | 1.45378300 |
| H | 1.04912200 | -0.24932900 | -0.36291800 |
| H | 1.30205600 | -1.88966600 | -0.96303400 |
| C | 3.01483000 | -1.00268600 | -0.06297300 |
| H | 0.96603300 | -3.57989600 | 2.35726700 |
| H | 1.18867400 | -3.84182300 | 0.62035300 |
| H | 2.51332300 | -3.19646400 | 1.58714800 |
| H | 3.36678800 | -0.32144600 | 0.71149900 |
| C | 3.90886100 | -1.60770700 | -0.85109600 |
| H | 3.54981900 | -2.33931500 | -1.57529600 |
| C | 5.37008200 | -1.41562600 | -0.80251000 |
| C | 5.90202200 | -0.21167200 | -0.50395500 |
| C | 6.17490600 | -2.64796500 | -1.12189200 |
| H | 5.21640000 | 0.61931200 | -0.35104400 |
| C | 7.30992300 | 0.11189400 | -0.36997400 |
| H | 7.24422100 | -2.45892300 | -1.18714600 |
| H | 6.00600100 | -3.41896900 | -0.36402400 |
| H | 5.84458200 | -3.06679200 | -2.07698400 |
| H | 8.04379500 | -0.67984600 | -0.49286200 |
| C | 7.75363700 | 1.35275700 | -0.09048800 |
| H | 7.03172700 | 2.15765900 | 0.04066000 |
| C | 9.15766400 | 1.70003000 | 0.05345200 |
| H | 9.87937100 | 0.89469100 | -0.07720900 |
| C | 9.59590700 | 2.93459300 | 0.33162700 |
| H | 8.86219900 | 3.72898300 | 0.45945200 |
| C | 11.03497800 | 3.31557600 | 0.48265700 |
| H | 11.22530000 | 3.73257900 | 1.47580400 |
| H | 11.68980600 | 2.45500000 | 0.33713400 |
| H | 11.31131100 | 4.08814000 | -0.24069100 |
| H | 1.30314400 | -1.10724800 | 1.98759200 |
| H | -0.95717100 | -1.94299700 | 2.18222600 |

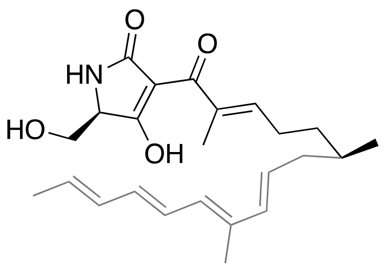


**INT$_{1a}$**

Energy (RM062X): -1289.838376 A.U.
Gibbs Free Energy: -1289.368464 A.U.

------------------------------------------------------------

| | | | |
|---|---|---|---|
| C | 4.74524000 | 0.12674700 | 1.23621000 |
| C | 5.33540200 | -0.74620800 | 0.10734900 |
| C | 4.49270500 | -1.97652600 | -0.26632700 |
| H | 5.09376200 | -2.57165200 | -0.96028500 |
| H | 4.55601400 | -0.50974100 | 2.10706600 |
| H | 5.51691500 | 0.84419200 | 1.53695400 |
| H | 4.35695000 | -2.59897800 | 0.62777800 |
| C | 3.12592900 | -1.72892900 | -0.92476500 |
| H | 3.19684600 | -0.90883200 | -1.64786600 |
| H | 2.85911800 | -2.61959000 | -1.50868500 |
| C | 2.00626100 | -1.46714300 | 0.03774200 |
| H | 2.16632100 | -1.77176200 | 1.07024700 |
| C | 3.48868500 | 0.87146400 | 0.86301400 |
| H | 2.58106700 | 0.60931300 | 1.39794800 |
| C | 3.42973300 | 1.77568900 | -0.12279800 |
| H | 4.34566800 | 2.00958600 | -0.66573200 |
| C | 2.20708100 | 2.40097400 | -0.65183900 |
| C | 1.03339800 | 2.36210800 | 0.01617800 |
| H | 1.02683100 | 1.95730000 | 1.02617300 |
| C | -0.26857600 | 2.70331000 | -0.51821300 |
| H | -0.33893400 | 3.03693700 | -1.55024900 |
| C | -1.41239400 | 2.53347300 | 0.17434600 |

| | | | |
|---|---|---|---|
| H | -1.36686700 | 2.18571800 | 1.20504500 |
| C | 0.82122500 | -0.92017900 | -0.27218100 |
| C | 0.48128400 | -0.46906800 | -1.67036000 |
| H | -0.46458500 | 0.07274800 | -1.69920600 |
| H | 1.26186900 | 0.20694100 | -2.02920400 |
| H | 0.41688000 | -1.31188600 | -2.36133200 |
| C | -0.12916000 | -0.74288100 | 0.86977900 |
| O | 0.28801800 | -0.54817300 | 2.00286600 |
| C | 6.73619600 | -1.20815300 | 0.51605000 |
| H | 7.20341900 | -1.79913100 | -0.27511700 |
| H | 7.38444300 | -0.35645000 | 0.73487000 |
| H | 6.68449800 | -1.83103400 | 1.41502500 |
| H | 5.42563300 | -0.11772300 | -0.78706400 |
| C | 2.36298000 | 2.98141200 | -2.03439400 |
| H | 1.48884500 | 3.53450500 | -2.37103000 |
| H | 3.22068300 | 3.65985600 | -2.05441800 |
| H | 2.57271100 | 2.18931600 | -2.76166500 |
| C | -2.73422400 | 2.73278400 | -0.39617800 |
| H | -2.77745600 | 3.03775400 | -1.44157000 |
| C | -3.87725300 | 2.54310700 | 0.27772600 |
| H | -3.81484300 | 2.24255100 | 1.32258200 |
| C | -5.24595700 | 2.71427700 | -0.30335000 |
| H | -5.81752400 | 3.46199700 | 0.25413600 |
| H | -5.19637800 | 3.02433500 | -1.34849400 |
| H | -5.81468000 | 1.78044200 | -0.24793300 |
| C | -1.58562700 | -0.82776100 | 0.63735700 |
| C | -2.25804900 | -1.50264600 | -0.32720000 |
| C | -2.60433000 | -0.28647100 | 1.57790000 |
| O | -1.75747400 | -2.23561500 | -1.30348800 |
| C | -3.75254500 | -1.38594100 | -0.13991500 |
| O | -2.44512200 | 0.41856800 | 2.55691500 |
| N | -3.83348100 | -0.73026200 | 1.14079700 |
| H | -4.15774200 | -0.76063200 | -0.94684600 |
| C | -4.44702900 | -2.74417600 | -0.17157600 |
| H | -4.66954400 | -0.24688300 | 1.43764600 |
| H | -4.00125400 | -3.40631800 | 0.57630600 |
| H | -5.51129900 | -2.62161900 | 0.04101300 |
| O | -4.24231100 | -3.24384200 | -1.49126000 |
| H | -2.48683000 | -2.72132900 | -1.73790300 |
| H | -4.55692500 | -4.15117900 | -1.55135800 |

------------------------------------------------------------

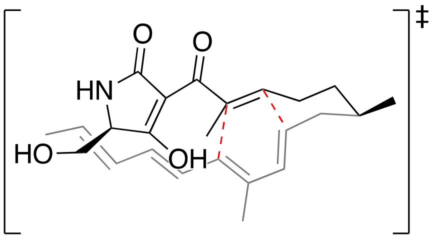


**TS$_{1a}$**

Energy (RM062X): -1289.815065 A.U.

Gibbs Free Energy: -1289.341427 A.U.

------------------------------------------------------------

| | | | |
|---|---|---|---|
| C | 4.52528000 | 0.16789900 | 1.26785400 |
| C | 5.39078000 | -0.46029300 | 0.16792000 |
| C | 4.69659800 | -1.67404700 | -0.45483700 |
| H | 5.33330300 | -2.08323800 | -1.24513300 |
| H | 4.36123400 | -0.58293700 | 2.04974400 |
| H | 5.07488700 | 0.99042900 | 1.73927100 |
| H | 4.60354700 | -2.45536200 | 0.31148300 |
| C | 3.31064700 | -1.37136800 | -1.02577600 |
| H | 3.38599200 | -0.62336300 | -1.82109900 |
| H | 2.90568700 | -2.27837100 | -1.48983200 |
| C | 2.33521300 | -0.91126300 | 0.03964600 |
| H | 2.45101400 | -1.42535000 | 0.99257600 |
| C | 3.17852600 | 0.68350000 | 0.79031300 |
| H | 2.42392100 | 0.75044300 | 1.57108700 |
| C | 3.12715800 | 1.64017100 | -0.22971800 |
| H | 4.02537200 | 1.81250300 | -0.81740800 |
| C | 1.94585000 | 2.25004500 | -0.68468700 |
| C | 0.74368700 | 2.04406200 | -0.02530300 |
| H | 0.74737300 | 1.69501800 | 1.00172300 |
| C | -0.53186100 | 2.43845100 | -0.56743900 |
| H | -0.58208300 | 2.71675100 | -1.61758600 |
| C | -1.67264300 | 2.40394100 | 0.15404900 |
| H | -1.61759700 | 2.13230300 | 1.20542100 |
| C | 0.98756800 | -0.65116700 | -0.28830000 |
| C | 0.58887700 | -0.49473900 | -1.72845000 |
| H | -0.40893300 | -0.06730600 | -1.83278600 |
| H | 1.29820400 | 0.17020500 | -2.23028500 |
| H | 0.60887800 | -1.44955800 | -2.26252300 |
| C | 0.05141400 | -0.73038100 | 0.82614700 |
| O | 0.44782500 | -0.71092800 | 2.00028400 |
| C | 6.75616200 | -0.85079500 | 0.73040000 |
| H | 7.38712600 | -1.29491900 | -0.04328400 |
| H | 7.27808800 | 0.01757200 | 1.13988000 |
| H | 6.64084400 | -1.58506000 | 1.53404100 |
| H | 5.54877100 | 0.28234700 | -0.62346500 |
| C | 1.98258600 | 3.02464200 | -1.98362500 |
| H | 1.49099300 | 3.99492300 | -1.88634200 |
| H | 3.01242900 | 3.19196800 | -2.30041800 |
| H | 1.47327300 | 2.48050700 | -2.78514500 |
| C | -2.99432600 | 2.65229400 | -0.38537200 |
| H | -3.06494000 | 2.92404600 | -1.43765900 |
| C | -4.11257200 | 2.52318100 | 0.34413400 |
| H | -4.01001200 | 2.24506500 | 1.39268300 |
| C | -5.50049600 | 2.72107400 | -0.17480100 |
| H | -6.01974800 | 3.50162000 | 0.38875100 |
| H | -5.49435300 | 2.99855000 | -1.22990300 |
| H | -6.09045000 | 1.80646700 | -0.05929500 |
| C | -1.41144600 | -0.85129400 | 0.59168500 |
| C | -2.07956900 | -1.56782900 | -0.34295300 |
| C | -2.43249400 | -0.35457800 | 1.55664200 |
| O | -1.57497300 | -2.31807800 | -1.31048100 |
| C | -3.57436100 | -1.50494300 | -0.13000700 |
| O | -2.28291800 | 0.34779700 | 2.54086100 |
| N | -3.65540900 | -0.83617600 | 1.14289800 |
| H | -4.02267100 | -0.90868900 | -0.93597900 |

| | | | |
|---|---|---|---|
| C | -4.21904600 | -2.88789600 | -0.12203800 |
| H | -4.50031400 | -0.37404600 | 1.44809400 |
| H | -3.72737900 | -3.52102100 | 0.62248000 |
| H | -5.28045500 | -2.80094400 | 0.12080800 |
| O | -4.03639200 | -3.40387500 | -1.43888700 |
| H | -2.29794900 | -2.83721600 | -1.71348100 |
| H | -4.32371400 | -4.32145200 | -1.47383300 |

--------------------------------------------------------------

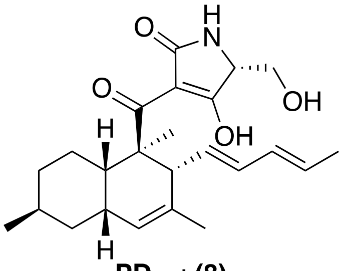


**PD$_{1a}$ : (8)**

Energy (RM062X): -1289.875388 A.U.
Gibbs Free Energy: -1289.398798 A.U.

--------------------------------------------------------------

| | | | |
|---|---|---|---|
| C | 4.59584600 | -0.46714600 | 1.03319300 |
| C | 5.42481200 | -0.09744700 | -0.19919600 |
| C | 4.77975300 | -0.69988500 | -1.44791900 |
| H | 5.35132800 | -0.41378800 | -2.33662300 |
| H | 4.65568300 | -1.55283600 | 1.18242400 |
| H | 5.02218600 | 0.00264600 | 1.92661800 |
| H | 4.83477000 | -1.79490300 | -1.37407900 |
| C | 3.31566500 | -0.28307300 | -1.61405700 |
| H | 3.25594800 | 0.79718100 | -1.78527800 |
| H | 2.91270100 | -0.77042100 | -2.50362600 |
| C | 2.49399300 | -0.67503900 | -0.37633500 |
| H | 2.57034700 | -1.76360500 | -0.28421200 |
| C | 3.12565600 | -0.06093000 | 0.89190800 |
| H | 2.57552600 | -0.45061300 | 1.75895500 |
| C | 2.95210200 | 1.43344000 | 0.83585600 |
| H | 3.79523200 | 2.07208700 | 1.09140400 |
| C | 1.79792700 | 1.97655200 | 0.45452800 |
| C | 0.59842200 | 1.06184800 | 0.21307300 |
| H | 0.22903200 | 0.84156700 | 1.22504000 |
| C | -0.52722000 | 1.76130800 | -0.50183800 |
| H | -0.34498000 | 2.08342200 | -1.52549100 |
| C | -1.72239100 | 2.00287900 | 0.04479900 |
| H | -1.90812000 | 1.69649700 | 1.07423500 |
| C | 0.97943400 | -0.34619600 | -0.45036700 |
| C | 0.46647100 | -0.44086700 | -1.89634400 |
| H | -0.61043400 | -0.29908200 | -1.95418700 |
| H | 0.93959400 | 0.32339800 | -2.51476700 |
| H | 0.70493100 | -1.41975400 | -2.31977100 |
| C | 0.21389500 | -1.38062100 | 0.39298300 |
| O | 0.76165500 | -2.23871100 | 1.04809600 |
| C | 6.87378700 | -0.55087500 | -0.04565300 |
| H | 7.46964800 | -0.27169700 | -0.91853900 |
| H | 7.33643400 | -0.10573800 | 0.83911700 |
| H | 6.92222600 | -1.63944700 | 0.06077500 |
| H | 5.41273600 | 0.99515800 | -0.30850300 |
| C | 1.58225300 | 3.46320500 | 0.39121400 |
| H | 0.77257900 | 3.76976600 | 1.06035200 |
| H | 2.49245200 | 3.99224800 | 0.67714800 |
| H | 1.29523400 | 3.78306700 | -0.61447400 |
| C | -2.83577400 | 2.63978700 | -0.65079000 |
| H | -2.65371100 | 2.98032000 | -1.66925000 |
| C | -4.04836000 | 2.79760900 | -0.10848000 |
| H | -4.20927900 | 2.44445600 | 0.90977600 |
| C | -5.21959400 | 3.42509200 | -0.79745000 |
| H | -5.57820500 | 4.29580300 | -0.24118100 |
| H | -4.95979400 | 3.74295100 | -1.80841700 |
| H | -6.05665000 | 2.72367400 | -0.85829800 |
| C | -1.27657700 | -1.24998600 | 0.50444400 |
| C | -2.27855800 | -1.52938800 | -0.34766000 |
| C | -1.88886600 | -0.80716100 | 1.77856500 |
| O | -2.19782600 | -2.03188800 | -1.57511100 |
| C | -3.62234100 | -1.21726900 | 0.27200000 |
| O | -1.31341700 | -0.42348600 | 2.78362600 |
| N | -3.25049700 | -0.88611000 | 1.62600000 |
| H | -4.05162000 | -0.35209900 | -0.25165000 |
| C | -4.59445100 | -2.38899600 | 0.18198400 |
| H | -3.86330100 | -0.37183300 | 2.24281600 |
| H | -4.14724600 | -3.27565500 | 0.64078100 |
| H | -5.52445200 | -2.14298300 | 0.69961000 |
| O | -4.82132700 | -2.58673600 | -1.21154000 |
| H | -3.09451100 | -2.28676900 | -1.86460000 |
| H | -5.34076700 | -3.38473300 | -1.34941200 |

--------------------------------------------------------------

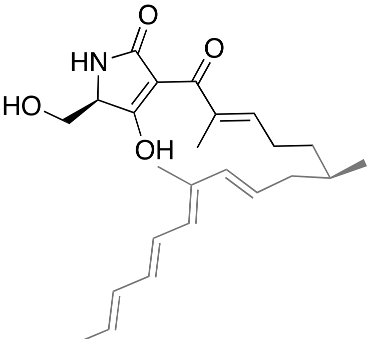


**INT$_{1b}$**

Energy (RM062X): -1289.832992 A.U.
Gibbs Free Energy: -1289.366722 A.U.

--------------------------------------------------------------

| | | | |
|---|---|---|---|
| C | 4.87692900 | 0.95414500 | 0.12084900 |
| C | 5.41888400 | -0.05179800 | -0.90551200 |
| C | 4.91021200 | -1.49046500 | -0.72792900 |
| H | 5.56688600 | -2.13775000 | -1.31610100 |
| H | 5.11134600 | 0.60646000 | 1.13393700 |
| H | 5.44070900 | 1.88607900 | -0.02093900 |
| H | 5.04120400 | -1.79283900 | 0.31869800 |
| C | 3.46626200 | -1.76702200 | -1.17523600 |
| H | 3.23829000 | -1.19030600 | -2.07625500 |

| | | | |
|---|---|---|---|
| H | 3.40217700 | -2.82051300 | -1.47960300 |
| C | 2.41319400 | -1.57463800 | -0.12907900 |
| H | 2.70284100 | -1.78935500 | 0.89697200 |
| C | 3.41726700 | 1.29930900 | 0.02670600 |
| H | 3.00016200 | 1.36277000 | -0.97840600 |
| C | 2.66752000 | 1.61839500 | 1.08902600 |
| H | 3.14454200 | 1.56460900 | 2.06763600 |
| C | 1.26365700 | 2.05792300 | 1.11790600 |
| C | 0.45662000 | 1.98202200 | 0.03525900 |
| H | 0.87363500 | 1.59161000 | -0.88891000 |
| C | -0.94708700 | 2.33445700 | -0.02139200 |
| H | -1.41684000 | 2.73579900 | 0.87109000 |
| C | -1.72604500 | 2.11500900 | -1.09941500 |
| H | -1.28662000 | 1.69877500 | -2.00494200 |
| C | 1.13393000 | -1.23683600 | -0.35423900 |
| C | 0.63339400 | -0.89850300 | -1.73745700 |
| H | 0.51731400 | -1.79194700 | -2.35489000 |
| H | -0.32670300 | -0.38710300 | -1.70522400 |
| H | 1.34287700 | -0.23801900 | -2.24287200 |
| C | 0.24292600 | -1.18061100 | 0.84798000 |
| O | 0.68123600 | -1.28922500 | 1.98261800 |
| C | 6.94877000 | -0.04758700 | -0.83709100 |
| H | 7.37912300 | -0.70472200 | -1.59590100 |
| H | 7.34900400 | 0.95756000 | -0.98848400 |
| H | 7.28593400 | -0.39984300 | 0.14329300 |
| H | 5.12032100 | 0.29087500 | -1.90515700 |
| C | 0.82173700 | 2.57611200 | 2.46077800 |
| H | 1.43082400 | 3.44085200 | 2.74305600 |
| H | -0.22729400 | 2.85538900 | 2.49223100 |
| H | 0.98366900 | 1.80662300 | 3.22112100 |
| C | -3.15844000 | 2.36067200 | -1.11461800 |
| H | -3.58950100 | 2.79438100 | -0.21237200 |
| C | -3.96215500 | 2.05829700 | -2.14320300 |
| H | -3.51857200 | 1.62021200 | -3.03639800 |
| C | -5.44469800 | 2.26344900 | -2.15529200 |
| H | -5.74529100 | 2.91104300 | -2.98397400 |
| H | -5.96767000 | 1.31216500 | -2.29429500 |
| H | -5.78869000 | 2.71268200 | -1.22212400 |
| C | -1.21795300 | -0.98902400 | 0.66244200 |
| C | -2.05292000 | -1.56594400 | -0.22266000 |
| O | -1.72576600 | -2.48380500 | -1.12763600 |
| H | -2.48565800 | -2.74158900 | -1.66398600 |
| C | -2.04668400 | -0.12233300 | 1.54505400 |
| O | -1.68453900 | 0.59043500 | 2.46277400 |
| N | -3.34482200 | -0.25429700 | 1.12193400 |
| H | -4.04087200 | 0.43152500 | 1.37582700 |
| C | -3.47590900 | -1.09124600 | -0.04670700 |
| H | -3.76608200 | -0.49979100 | -0.92621800 |
| C | -4.46838300 | -2.23506000 | 0.12831300 |
| H | -5.44427300 | -1.81190300 | 0.38409000 |
| H | -4.57742700 | -2.77238600 | -0.82188100 |
| O | -3.98607200 | -3.08772300 | 1.14656900 |
| H | -4.64064800 | -3.77105100 | 1.31593400 |

-------------------------------------------------------------

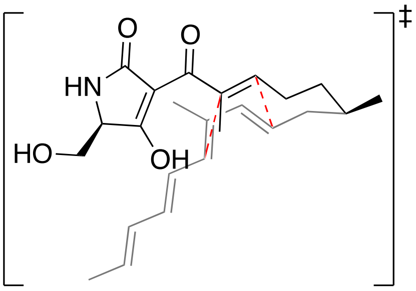


**TS$_{1b}$**

Energy (RM062X): -1289.812275 A.U.
Gibbs Free Energy: -1289.340702 A.U.

-------------------------------------------------------------

| | | | |
|---|---|---|---|
| C | 4.70371700 | 0.45470100 | 0.53384900 |
| C | 5.39152700 | 0.12767200 | -0.79094100 |
| C | 4.71524100 | -1.05342100 | -1.48321200 |
| H | 5.22092800 | -1.25475400 | -2.43249900 |
| H | 4.79619600 | -0.40347200 | 1.21191100 |
| H | 5.23764400 | 1.28509900 | 1.01031400 |
| H | 4.83765600 | -1.95011600 | -0.86173800 |
| C | 3.23162100 | -0.81380200 | -1.74398300 |
| H | 3.10875000 | 0.09244300 | -2.34609800 |
| H | 2.82901700 | -1.63556600 | -2.34760700 |
| C | 2.39141100 | -0.73172000 | -0.48043700 |
| H | 2.67684100 | -1.43133400 | 0.30091800 |
| C | 3.23891400 | 0.83958000 | 0.41920500 |
| H | 3.00637600 | 1.49187200 | -0.42150100 |
| C | 2.53794400 | 1.05331600 | 1.60942700 |
| H | 3.00507200 | 0.69568400 | 2.52522800 |
| C | 1.22953100 | 1.55248100 | 1.71988300 |
| C | 0.50805100 | 1.86140600 | 0.58073800 |
| H | 1.03168200 | 1.99286400 | -0.35815000 |
| C | -0.89352700 | 2.19416000 | 0.57335900 |
| H | -1.45419400 | 2.05560000 | 1.49444900 |
| C | -1.55210300 | 2.60730700 | -0.53213400 |
| H | -0.99924000 | 2.74461700 | -1.46036600 |
| C | 1.00354800 | -0.53728800 | -0.60312500 |
| C | 0.42353600 | -0.00767900 | -1.88443600 |
| H | 0.44104400 | -0.76306800 | -2.67624200 |
| H | -0.60827000 | 0.31874600 | -1.75490700 |
| H | 1.00405200 | 0.84719200 | -2.24599600 |
| C | 0.19191000 | -1.01177300 | 0.50524300 |
| O | 0.69675700 | -1.39829100 | 1.56647800 |
| C | 6.87689000 | -0.14650500 | -0.56684500 |
| H | 7.38265400 | -0.36292200 | -1.51087200 |
| H | 7.37210400 | 0.70998300 | -0.10277900 |
| H | 7.00908500 | -1.01053300 | 0.09208300 |
| H | 5.29345100 | 1.00383900 | -1.44626700 |
| C | 0.58109400 | 1.59227300 | 3.08167500 |
| H | 1.31370500 | 1.37245800 | 3.85921900 |
| H | 0.15295100 | 2.57712700 | 3.28599800 |
| H | -0.22379000 | 0.85619000 | 3.13886000 |
| C | -2.97835100 | 2.86773100 | -0.56984000 |
| H | -3.52769100 | 2.73531500 | 0.36147700 |

| | | | |
|---|---|---|---|
| C | -3.63671600 | 3.24382100 | -1.67548400 |
| H | -3.07137000 | 3.36846800 | -2.59772500 |
| C | -5.10698600 | 3.50811500 | -1.74421200 |
| H | -5.30249100 | 4.53360100 | -2.07102900 |
| H | -5.58662200 | 2.84965400 | -2.47425800 |
| H | -5.58291100 | 3.35632300 | -0.77427100 |
| C | -1.29130700 | -1.08677800 | 0.34434400 |
| C | -1.98765900 | -1.64862200 | -0.66021400 |
| O | -1.47813300 | -2.22111200 | -1.75192100 |
| H | -2.17426200 | -2.53179700 | -2.34380900 |
| C | -2.28198000 | -0.70481900 | 1.38573500 |
| O | -2.08040000 | -0.27451600 | 2.50873300 |
| N | -3.52802900 | -0.90180300 | 0.84398000 |
| H | -4.34027400 | -0.93257000 | 1.44353700 |
| C | -3.47490600 | -1.62381300 | -0.40661500 |
| H | -3.98113100 | -1.07606500 | -1.21046300 |
| C | -4.07618200 | -3.02532400 | -0.32561500 |
| H | -5.13241700 | -2.93785800 | -0.05316000 |
| H | -4.02224300 | -3.49812000 | -1.31341200 |
| O | -3.35101000 | -3.76153900 | 0.63700000 |
| H | -3.78511600 | -4.60873100 | 0.77079300 |

------------------------------------------------------------

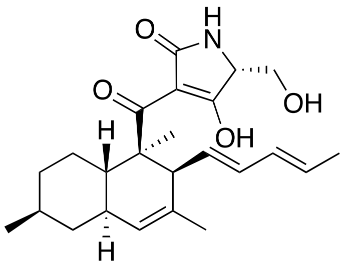

**PD$_{1b}$ : (6)**

Energy (RM062X): -1289.872928 A.U.

Gibbs Free Energy: -1289.397524 A.U.

------------------------------------

| | | | |
|---|---|---|---|
| C | 4.69847700 | 0.74623300 | 0.15876000 |
| C | 5.20152600 | -0.41531400 | -0.69726100 |
| C | 4.27537000 | -1.61959800 | -0.51500300 |
| H | 4.62146800 | -2.45374800 | -1.13336900 |
| H | 4.77684300 | 0.45705500 | 1.21656700 |
| H | 5.34050600 | 1.62375700 | 0.02416800 |
| H | 4.33031800 | -1.95140600 | 0.53040300 |
| C | 2.83181700 | -1.26783300 | -0.87255700 |
| H | 2.79368100 | -1.00487800 | -1.93500700 |
| H | 2.18097600 | -2.13840700 | -0.73847100 |
| C | 2.31620200 | -0.09812400 | -0.02557500 |
| H | 2.36981400 | -0.40415800 | 1.02461000 |
| C | 3.24403300 | 1.13249900 | -0.12908800 |
| H | 3.18142900 | 1.52704600 | -1.15645800 |
| C | 2.75802800 | 2.16034600 | 0.85679400 |
| H | 3.47915800 | 2.61900300 | 1.53048900 |
| C | 1.45837400 | 2.43771100 | 0.96136700 |
| C | 0.54064000 | 1.81188400 | -0.07413800 |
| H | 0.80095400 | 2.27655500 | -1.03355500 |
| C | -0.92174900 | 2.06652000 | 0.16550400 |
| H | -1.32773300 | 1.76916300 | 1.13055200 |
| C | -1.75929100 | 2.58206800 | -0.74048700 |
| H | -1.37960100 | 2.88333200 | -1.71664400 |
| C | 0.82817300 | 0.26227600 | -0.31534000 |
| C | 0.42540600 | -0.01467000 | -1.76695300 |
| H | 0.47198400 | -1.07096600 | -2.02087900 |
| H | -0.58890800 | 0.34351100 | -1.95500500 |
| H | 1.10184100 | 0.52497500 | -2.43495500 |
| C | -0.05546200 | -0.47818700 | 0.70481000 |
| O | 0.21131200 | -0.44563800 | 1.88768800 |
| C | 6.64762100 | -0.76417900 | -0.35654500 |
| H | 7.01276500 | -1.58499600 | -0.97910500 |
| H | 7.30750000 | 0.09476400 | -0.50343000 |
| H | 6.72693100 | -1.07455600 | 0.69029700 |
| H | 5.15282800 | -0.10991600 | -1.75163500 |
| C | 0.89451600 | 3.38452800 | 1.98188900 |
| H | 1.69345700 | 3.82198300 | 2.58227500 |
| H | 0.33129700 | 4.19145200 | 1.50393000 |
| H | 0.20101500 | 2.86929700 | 2.65326100 |
| C | -3.18880600 | 2.76016900 | -0.50346800 |
| H | -3.56384700 | 2.43117800 | 0.46470900 |
| C | -4.03241000 | 3.28056100 | -1.40116600 |
| H | -3.63528900 | 3.59742000 | -2.36452300 |
| C | -5.50230500 | 3.46484200 | -1.18484600 |
| H | -5.78172800 | 4.51779200 | -1.28274100 |
| H | -6.07866000 | 2.91534100 | -1.93461200 |
| H | -5.80237100 | 3.11791300 | -0.19473800 |
| C | -1.31829900 | -1.15405700 | 0.28694700 |
| C | -1.51127900 | -2.22628700 | -0.50014500 |
| O | -0.57045900 | -2.92054900 | -1.13850400 |
| H | -0.94872700 | -3.63405900 | -1.66773300 |
| C | -2.63510300 | -0.82160100 | 0.90179000 |
| O | -2.87446900 | -0.00085500 | 1.76908600 |
| N | -3.56692300 | -1.61129700 | 0.28134800 |
| H | -4.49054500 | -1.71352300 | 0.67718600 |
| C | -2.96252400 | -2.64388600 | -0.53022000 |
| H | -3.33636000 | -2.61828800 | -1.55997400 |
| C | -3.17138500 | -4.05057800 | 0.02733400 |
| H | -4.24566900 | -4.25496200 | 0.06345400 |
| H | -2.71398800 | -4.78018300 | -0.65091600 |
| O | -2.58790000 | -4.10463800 | 1.31167800 |
| H | -2.81016600 | -4.94689700 | 1.71840800 |

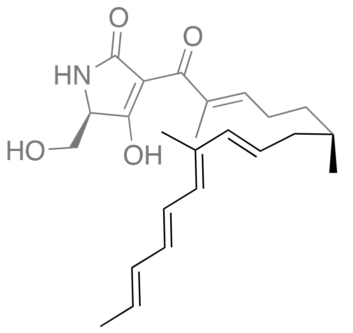


**INT$_{1c}$**

Energy (RM062X): -1289.831335 A.U.

Gibbs Free Energy: -1289.36425 A.U.

------------------------------------------------------------

| | | | |
|---|---|---|---|
| C | -5.43489400 | 0.37897000 | -0.71849900 |
| C | -4.80971800 | -0.75674300 | -1.54824900 |
| C | -4.94348700 | 0.47941900 | 0.73985300 |
| H | -4.97785100 | -1.70895100 | -1.03269900 |
| H | -5.37831100 | -0.81098000 | -2.48170300 |
| H | -5.05605500 | -0.49153100 | 1.23339700 |
| H | -5.62562900 | 1.17204300 | 1.25084100 |
| C | -3.31998600 | -0.64319400 | -1.90565100 |
| H | -3.05568300 | 0.38937100 | -2.15498700 |
| H | -3.14278600 | -1.20899500 | -2.82981800 |
| C | -2.37817500 | -1.18323200 | -0.87094800 |
| H | -2.79937800 | -1.82611400 | -0.10208000 |
| C | -3.54043600 | 0.97746700 | 0.93764100 |
| H | -3.23742900 | 1.84120500 | 0.34868600 |
| C | -2.68082000 | 0.46100900 | 1.82320000 |
| H | -3.01633500 | -0.37174400 | 2.44131900 |
| C | -1.28966000 | 0.87786200 | 2.05184700 |
| C | -0.57944000 | 1.48895000 | 1.07872600 |
| H | -1.08302600 | 1.69150500 | 0.13669000 |
| C | 0.81947400 | 1.86244000 | 1.11488800 |
| H | 1.40550600 | 1.61857700 | 1.99698200 |
| C | 1.43329600 | 2.45971600 | 0.07323900 |
| H | 0.85238000 | 2.69378200 | -0.81830600 |
| C | -1.05273400 | -0.97275400 | -0.84735800 |
| C | -0.36260900 | -0.09424700 | -1.86043000 |
| H | -0.27272100 | -0.58991800 | -2.82963600 |
| H | -0.93242900 | 0.82615800 | -2.01231100 |
| H | 0.63705600 | 0.18944000 | -1.53092100 |
| C | -0.29836500 | -1.61750500 | 0.26690600 |
| O | -0.85826600 | -2.11492200 | 1.23087100 |
| C | -5.36120300 | 1.72656800 | -1.44146300 |
| H | -5.82942300 | 1.65771300 | -2.42662000 |
| H | -5.88452200 | 2.49840700 | -0.87095700 |
| H | -4.33398200 | 2.06701500 | -1.58907000 |
| H | -6.49769600 | 0.12225600 | -0.64205800 |
| C | -0.72778500 | 0.47194600 | 3.38571900 |
| H | 0.26107700 | 0.88247300 | 3.57596500 |
| H | -1.40153500 | 0.78508400 | 4.18845200 |
| H | -0.64342300 | -0.61826600 | 3.41490300 |
| C | 2.84477800 | 2.80112700 | 0.04197300 |
| H | 3.42663700 | 2.57146300 | 0.93380300 |
| C | 3.45849000 | 3.34479500 | -1.01815300 |
| H | 2.86592400 | 3.56325100 | -1.90543300 |
| C | 4.91757100 | 3.66897100 | -1.08149900 |
| H | 5.07498300 | 4.73287800 | -1.28124600 |
| H | 5.41932300 | 3.41307400 | -0.14686200 |
| H | 5.40185800 | 3.12056200 | -1.89502100 |
| C | 1.18573800 | -1.70021000 | 0.20058000 |
| C | 1.96531800 | -2.05667200 | -0.83946900 |
| C | 2.06370400 | -1.64809700 | 1.40157000 |
| O | 1.56164400 | -2.28410200 | -2.08615500 |
| C | 3.40369900 | -2.24039600 | -0.42595800 |
| O | 1.77972800 | -1.28377900 | 2.52611600 |
| N | 3.30578200 | -2.09578800 | 1.00939500 |
| H | 2.28702700 | -2.58817400 | -2.64611800 |
| H | 3.73497200 | -3.24581300 | -0.71248000 |
| C | 4.36979500 | -1.22298100 | -1.06238100 |
| H | 4.11284300 | -1.83036600 | 1.55710300 |
| H | 5.38597300 | -1.51345400 | -0.79189500 |
| H | 4.29054100 | -1.27513300 | -2.15341300 |
| O | 4.18554300 | 0.08559200 | -0.57982900 |
| H | 3.42714500 | 0.51212000 | -0.99479700 |

------------------------------------------------------------

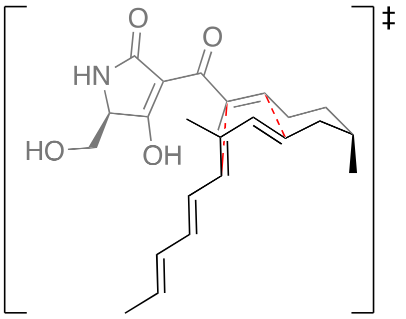


**TS$_{1c}$**

Energy (RM062X): -1289.811758 A.U.

Gibbs Free Energy: -1289.339651 A.U.

------------------------------------------------------------

| | | | |
|---|---|---|---|
| C | -5.45384000 | 0.22187300 | -0.73310300 |
| C | -4.78754100 | -0.84194400 | -1.61132200 |
| C | -4.77252200 | 0.31100200 | 0.63679700 |
| H | -4.94755600 | -1.82868800 | -1.16193900 |
| H | -5.27385900 | -0.85760800 | -2.59137500 |
| H | -4.89002400 | -0.64095100 | 1.16690500 |
| H | -5.29166100 | 1.06727100 | 1.23668100 |
| C | -3.28606600 | -0.62954900 | -1.79790500 |
| H | -3.09051600 | 0.34732200 | -2.24983600 |
| H | -2.90794100 | -1.36867600 | -2.51393400 |
| C | -2.48595500 | -0.79426500 | -0.51685300 |
| H | -2.83976100 | -1.58253700 | 0.14257600 |
| C | -3.29464500 | 0.66250900 | 0.60360700 |
| H | -3.01159300 | 1.41983700 | -0.12366500 |
| C | -2.60447600 | 0.66317800 | 1.81798900 |

| | | | |
|---|---|---|---|
| H | -3.09965600 | 0.19839100 | 2.66863100 |
| C | -1.27330600 | 1.07305000 | 2.00455900 |
| C | -0.53168300 | 1.50420900 | 0.92063800 |
| H | -1.04715100 | 1.80122700 | 0.01554700 |
| C | 0.88238300 | 1.77920300 | 0.94824100 |
| H | 1.45186500 | 1.45711800 | 1.81695100 |
| C | 1.52637300 | 2.37657200 | -0.07908700 |
| H | 0.95294100 | 2.69245700 | -0.94968100 |
| C | -1.08572700 | -0.67035000 | -0.57008400 |
| C | -0.42225400 | -0.02258700 | -1.75299100 |
| H | -0.46901800 | -0.66107400 | -2.64023500 |
| H | -0.91934100 | 0.91850800 | -2.00895700 |
| H | 0.62661400 | 0.19892100 | -1.55408700 |
| C | -0.34804600 | -1.34859100 | 0.48178400 |
| O | -0.91115100 | -1.82700100 | 1.47337300 |
| C | -5.52590300 | 1.58520200 | -1.42703700 |
| H | -6.07262300 | 1.50593900 | -2.36993400 |
| H | -6.04347800 | 2.31145800 | -0.79525500 |
| H | -4.53732100 | 1.99126100 | -1.65383600 |
| H | -6.48398700 | -0.10267500 | -0.55214600 |
| C | -0.63207500 | 0.87842400 | 3.35528600 |
| H | -0.14993200 | 1.79722800 | 3.69931400 |
| H | -1.38016800 | 0.58931300 | 4.09456000 |
| H | 0.12943400 | 0.09750200 | 3.30076900 |
| C | 2.95516300 | 2.62183100 | -0.11150000 |
| H | 3.53000000 | 2.31076100 | 0.75962600 |
| C | 3.58887100 | 3.17659000 | -1.15490000 |
| H | 2.99951500 | 3.47439500 | -2.02081100 |
| C | 5.06340800 | 3.41091600 | -1.22670300 |
| H | 5.28313400 | 4.47142300 | -1.38033900 |
| H | 5.56136800 | 3.07997000 | -0.31411600 |
| H | 5.49875100 | 2.87322800 | -2.07425900 |
| C | 1.12593500 | -1.56090700 | 0.34338600 |
| C | 1.78579700 | -2.08953600 | -0.70513400 |
| C | 2.09658700 | -1.50418900 | 1.46932100 |
| O | 1.26355600 | -2.39593800 | -1.89353800 |
| C | 3.23169800 | -2.37007900 | -0.38398900 |
| O | 1.93933200 | -1.03519300 | 2.58298100 |
| N | 3.25971500 | -2.09646200 | 1.03461400 |
| H | 1.90951000 | -2.83287000 | -2.46220600 |
| H | 3.45622100 | -3.42145600 | -0.60066400 |
| C | 4.21892700 | -1.49726200 | -1.18046500 |
| H | 4.12633800 | -1.87058700 | 1.50303600 |
| H | 5.23073200 | -1.83495100 | -0.95266900 |
| H | 4.05262000 | -1.64619500 | -2.25316300 |
| O | 4.16511800 | -0.13728900 | -0.82258500 |
| H | 3.34707700 | 0.27154100 | -1.12885100 |

-----------------------------------------------------------

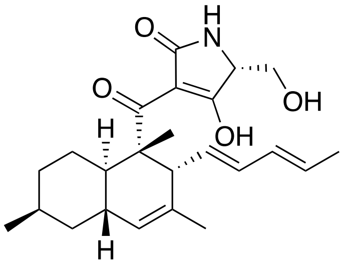

**PD$_{1c}$ : (7)**

Energy (RM062X): -1289.871318 A.U.
Gibbs Free Energy: -1289.394244 A.U.

-----------------------------------------------------------

| | | | |
|---|---|---|---|
| C | -5.42273200 | -0.47677600 | -0.57815300 |
| C | -4.57856000 | -1.72679200 | -0.29506800 |
| C | -4.84679400 | 0.70838300 | 0.20376100 |
| H | -4.67332000 | -1.98423500 | 0.76637600 |
| H | -4.96607000 | -2.57597500 | -0.86645100 |
| H | -4.95779300 | 0.50801200 | 1.27704000 |
| H | -5.41880000 | 1.61817300 | -0.00899600 |
| C | -3.10075800 | -1.50876100 | -0.62619100 |
| H | -2.99116500 | -1.35227500 | -1.70280200 |
| H | -2.53158800 | -2.41293100 | -0.38408400 |
| C | -2.52692300 | -0.31571400 | 0.14973200 |
| H | -2.64157400 | -0.54030800 | 1.21458100 |
| C | -3.36050300 | 0.96882900 | -0.07140400 |
| H | -3.23498200 | 1.29243700 | -1.11634500 |
| C | -2.82513800 | 2.02460300 | 0.85849900 |
| H | -3.52048800 | 2.55913600 | 1.50255200 |
| C | -1.51090700 | 2.22721500 | 0.95307000 |
| C | -0.64635000 | 1.47138800 | -0.03951600 |
| H | -0.92809100 | 1.83190900 | -1.03669700 |
| C | 0.82818800 | 1.70912400 | 0.12207200 |
| H | 1.25607300 | 1.54680200 | 1.11007000 |
| C | 1.64263300 | 2.07131900 | -0.87527500 |
| H | 1.23118400 | 2.24096100 | -1.87021500 |
| C | -1.00458200 | -0.08408600 | -0.09952500 |
| C | -0.56013000 | -0.56185100 | -1.48429700 |
| H | -0.69376400 | -1.63103900 | -1.62201300 |
| H | -1.14988100 | -0.04285800 | -2.24466200 |
| H | 0.49070400 | -0.32194300 | -1.65904600 |
| C | -0.20791000 | -0.72329100 | 1.05143300 |
| O | -0.62010700 | -0.67656800 | 2.19148900 |
| C | -5.53806300 | -0.17950100 | -2.07578700 |
| H | -5.96834500 | -1.03280900 | -2.60675500 |
| H | -6.18327400 | 0.68623700 | -2.24659300 |
| H | -4.56820800 | 0.03790300 | -2.52913500 |
| H | -6.43479500 | -0.66220500 | -0.20244800 |
| C | -0.88250600 | 3.17995300 | 1.92888700 |
| H | -0.26013700 | 3.91984400 | 1.41713600 |
| H | -1.64950300 | 3.70257900 | 2.50215500 |
| H | -0.23135500 | 2.64806400 | 2.62976500 |
| C | 3.08503100 | 2.23191100 | -0.72331600 |
| H | 3.49460600 | 2.05432300 | 0.27071700 |

| | | | |
|---|---|---|---|
| C | 3.90695400 | 2.54588900 | -1.73016500 |
| H | 3.47869500 | 2.71154500 | -2.71771600 |
| C | 5.39254000 | 2.67438100 | -1.60820100 |
| H | 5.72585500 | 3.67191600 | -1.90821800 |
| H | 5.72201500 | 2.49113700 | -0.58418900 |
| H | 5.89722400 | 1.96162000 | -2.26702700 |
| C | 1.15655100 | -1.30034100 | 0.84818700 |
| C | 1.59769000 | -2.32905200 | 0.09855700 |
| C | 2.28375100 | -0.91183400 | 1.74843100 |
| O | 0.89470000 | -3.06818600 | -0.75487500 |
| C | 3.05762700 | -2.61200500 | 0.34996200 |
| O | 2.31250300 | -0.00452400 | 2.55957700 |
| N | 3.31541800 | -1.78232500 | 1.50307200 |
| H | 1.41463000 | -3.80718800 | -1.09616800 |
| H | 3.20165200 | -3.67601000 | 0.56679200 |
| C | 3.93423300 | -2.23317400 | -0.86217600 |
| H | 4.25679500 | -1.51088000 | 1.75204100 |
| H | 4.95427400 | -2.56352400 | -0.66468400 |
| H | 3.57455600 | -2.76613500 | -1.74934100 |
| O | 4.00315800 | -0.84348800 | -1.07345500 |
| H | 3.13487700 | -0.48277500 | -1.29012500 |

--------------------------------------------------------------

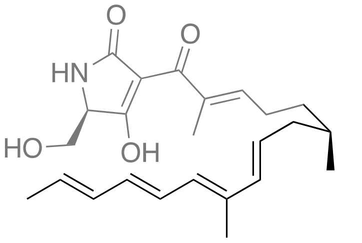

**INT$_{1d}$**

Energy (RM062X): -1289.837325 A.U.
Gibbs Free Energy: -1289.369181 A.U.

--------------------------------------------------------------

| | | | |
|---|---|---|---|
| C | -5.40304300 | -0.87606100 | 0.27662000 |
| C | -4.51458600 | -2.13548300 | 0.24400500 |
| C | -4.93376800 | 0.26131400 | 1.21982100 |
| H | -4.35014300 | -2.48443000 | 1.26979700 |
| H | -5.10939400 | -2.91151200 | -0.24662500 |
| H | -4.86284800 | -0.14056000 | 2.23534700 |
| H | -5.72081000 | 1.02250400 | 1.22416200 |
| C | -3.15546200 | -2.05917700 | -0.48400500 |
| H | -3.20142600 | -1.37713900 | -1.33724400 |
| H | -2.94871300 | -3.04906200 | -0.91066700 |
| C | -1.99476900 | -1.70964700 | 0.39829800 |
| H | -2.08676800 | -1.98552300 | 1.44736200 |
| C | -3.62733100 | 0.88858400 | 0.83296000 |
| H | -2.72532800 | 0.42515500 | 1.21910300 |
| C | -3.50448800 | 1.91947200 | -0.01195800 |
| H | -4.40893900 | 2.36218900 | -0.42967200 |
| C | -2.22934900 | 2.43047400 | -0.54296700 |
| C | -1.06786500 | 2.29546100 | 0.13240100 |
| H | -1.10164700 | 1.89959800 | 1.14591500 |
| C | 0.25571000 | 2.53850000 | -0.40453000 |
| H | 0.34224200 | 2.84205000 | -1.44477600 |
| C | 1.39415700 | 2.33122200 | 0.28625500 |
| H | 1.34398500 | 2.01302400 | 1.32692500 |
| C | -0.85837000 | -1.11540000 | 0.01083400 |
| C | -0.60506800 | -0.67150400 | -1.40878900 |
| H | -0.60950400 | -1.51408300 | -2.10226800 |
| H | -1.39118600 | 0.02741600 | -1.71009000 |
| H | 0.35108000 | -0.15484000 | -1.50029500 |
| C | 0.13942500 | -0.86209000 | 1.09961500 |
| O | -0.23479500 | -0.54333700 | 2.21803600 |
| C | -5.70607600 | -0.35511800 | -1.12868900 |
| H | -6.08161000 | -1.16155500 | -1.76431100 |
| H | -6.46541700 | 0.43057700 | -1.09175400 |
| H | -4.82009100 | 0.06844200 | -1.60704000 |
| H | -6.35441100 | -1.21416000 | 0.70353500 |
| C | -2.33159800 | 3.01107900 | -1.93055800 |
| H | -3.17168700 | 3.70943300 | -1.97785700 |
| H | -1.43458100 | 3.54170500 | -2.24298900 |
| H | -2.53844100 | 2.22031300 | -2.66055700 |
| C | 2.71637900 | 2.46312100 | -0.30453800 |
| H | 2.75571000 | 2.72006800 | -1.36309500 |
| C | 3.86033900 | 2.26262100 | 0.36290200 |
| H | 3.80426000 | 2.00234500 | 1.41810400 |
| C | 5.22492200 | 2.34105100 | -0.24467100 |
| H | 5.74896900 | 1.38579500 | -0.13386700 |
| H | 5.17480600 | 2.58676100 | -1.30709600 |
| H | 5.83862100 | 3.09410300 | 0.25832100 |
| C | 1.57994400 | -1.04069300 | 0.82830200 |
| C | 2.17560400 | -1.78526900 | -0.13445500 |
| C | 2.66686900 | -0.49549500 | 1.69277000 |
| O | 1.60063300 | -2.49361200 | -1.08980800 |
| C | 3.67883600 | -1.75854100 | -0.00172200 |
| O | 2.57682500 | 0.18628600 | 2.69798400 |
| N | 3.85278600 | -0.88650600 | 1.12768600 |
| H | 2.28092500 | -2.74935900 | -1.74134300 |
| H | 4.03990600 | -2.77486000 | 0.19699400 |
| C | 4.33893300 | -1.23859700 | -1.27784300 |
| H | 4.73230700 | -0.76466700 | 1.60723800 |
| H | 3.93158000 | -0.25242400 | -1.52447600 |
| H | 5.41893300 | -1.15974600 | -1.13030500 |
| O | 4.02758200 | -2.20135200 | -2.28054700 |
| H | 4.29209100 | -1.87484200 | -3.14596500 |

--------------------------------------------------------------

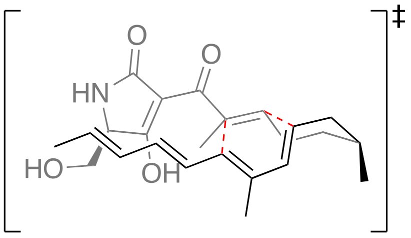

**TS$_{1d}$**

Energy (RM062X): -1289.809341 A.U.

Gibbs Free Energy: -1289.335503 A.U.

------------------------------------------------------------

| | | | |
|---|---|---|---|
| C | -5.46843100 | -0.54146500 | 0.43407800 |
| C | -4.75731600 | -1.79567000 | -0.09198100 |
| C | -4.57024400 | 0.25912000 | 1.39823300 |
| H | -4.60713400 | -2.49888400 | 0.73540400 |
| H | -5.40781000 | -2.29552700 | -0.81637500 |
| H | -4.40198600 | -0.36243300 | 2.28374300 |
| H | -5.11880600 | 1.14262000 | 1.74305700 |
| C | -3.39536200 | -1.51620200 | -0.73269500 |
| H | -3.48949500 | -0.82906500 | -1.57881900 |
| H | -2.99567000 | -2.45167800 | -1.14087400 |
| C | -2.40134000 | -0.97904900 | 0.27820200 |
| H | -2.52212000 | -1.40518600 | 1.27301200 |
| C | -3.20772900 | 0.69638100 | 0.88062900 |
| H | -2.45770300 | 0.78840200 | 1.66406000 |
| C | -3.09938600 | 1.59328400 | -0.18913700 |
| H | -3.98138400 | 1.79261500 | -0.78791100 |
| C | -1.88626100 | 2.12578500 | -0.65805200 |
| C | -0.69895900 | 1.90667300 | 0.02362000 |
| H | -0.72423700 | 1.61236400 | 1.06720800 |
| C | 0.59241300 | 2.23302200 | -0.52668900 |
| H | 0.65509700 | 2.46570800 | -1.58716100 |
| C | 1.73157200 | 2.19841600 | 0.19768300 |
| H | 1.67122800 | 1.97020500 | 1.25948200 |
| C | -1.05005700 | -0.78450900 | -0.07668100 |
| C | -0.66525000 | -0.73519000 | -1.52764200 |
| H | -0.73423000 | -1.71844100 | -2.00245700 |
| H | -1.35503100 | -0.07127200 | -2.05897200 |
| H | 0.34863200 | -0.35836100 | -1.66858800 |
| C | -0.10760600 | -0.82109400 | 1.03588800 |
| O | -0.49245600 | -0.68214500 | 2.20526000 |
| C | -6.07139500 | 0.27424600 | -0.71667800 |
| H | -6.92162600 | -0.26530200 | -1.14131200 |
| H | -6.42802200 | 1.24877900 | -0.37286800 |
| H | -5.36079500 | 0.44028400 | -1.52852500 |
| H | -6.31085000 | -0.88229000 | 1.04539700 |
| C | -1.88003000 | 2.83412100 | -1.99517400 |
| H | -2.89956000 | 3.02562800 | -2.33112500 |
| H | -1.35076300 | 3.78791300 | -1.94259300 |
| H | -1.38638100 | 2.22946000 | -2.76245000 |
| C | 3.05725400 | 2.39373600 | -0.35322200 |
| H | 3.13040500 | 2.63355400 | -1.41322000 |
| C | 4.17389900 | 2.23958600 | 0.37278500 |
| H | 4.06810100 | 1.98317600 | 1.42608200 |
| C | 5.56445900 | 2.35758500 | -0.16134500 |
| H | 6.10518200 | 1.41477900 | -0.02970300 |
| H | 5.56335000 | 2.61160100 | -1.22252600 |
| H | 6.12972200 | 3.12124600 | 0.38069600 |
| C | 1.34354500 | -1.04946800 | 0.80395600 |
| C | 1.95403200 | -1.87517400 | -0.07578800 |
| C | 2.41377200 | -0.51686600 | 1.70140700 |
| O | 1.39631400 | -2.65426800 | -0.99344200 |
| C | 3.45159500 | -1.88689000 | 0.11371000 |
| O | 2.30859200 | 0.20052100 | 2.68195600 |
| N | 3.60802500 | -0.96852300 | 1.20702100 |
| H | 2.09157400 | -2.96063600 | -1.60352900 |
| H | 3.77585600 | -2.90156600 | 0.37443000 |
| C | 4.17666300 | -1.44501500 | -1.15595800 |
| H | 4.47458600 | -0.83260700 | 1.70554300 |
| H | 3.80692700 | -0.46116100 | -1.46284200 |
| H | 5.25184700 | -1.38602400 | -0.96818000 |
| O | 3.88029900 | -2.44413200 | -2.12721700 |
| H | 4.18390500 | -2.16023700 | -2.99458500 |

------------------------------------------------------------

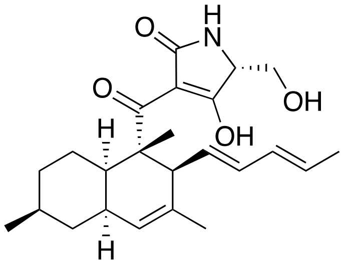

**PD*1d***

Energy (RM062X): -1289.870423 A.U.
Gibbs Free Energy: -1289.39231 A.U.

------------------------------------------------------------

| | | | |
|---|---|---|---|
| C | -5.44531500 | -0.49421100 | 0.37465200 |
| C | -4.61024500 | -1.76692100 | 0.19855100 |
| C | -4.66451400 | 0.49819200 | 1.24694800 |
| H | -4.49569700 | -2.25616800 | 1.17336100 |
| H | -5.13659900 | -2.47227800 | -0.45231700 |
| H | -4.61972400 | 0.08499500 | 2.26111400 |
| H | -5.20750400 | 1.44694500 | 1.31829600 |
| C | -3.21988400 | -1.47670800 | -0.37235800 |
| H | -3.31030300 | -1.05228200 | -1.37699800 |
| H | -2.68494000 | -2.42307700 | -0.47887100 |
| C | -2.42421800 | -0.52888400 | 0.53730600 |
| H | -2.34009900 | -1.02559700 | 1.50792800 |
| C | -3.22444700 | 0.77777500 | 0.79527100 |
| H | -2.72834200 | 1.27538600 | 1.64015100 |
| C | -3.09604600 | 1.69217000 | -0.39243400 |
| H | -3.97934100 | 2.12790900 | -0.85105000 |
| C | -1.87412000 | 1.96270600 | -0.85508500 |
| C | -0.72115500 | 1.33945900 | -0.08106300 |
| H | -0.76880500 | 1.72797900 | 0.94399000 |
| C | 0.63314600 | 1.69021300 | -0.63208300 |
| H | 0.83300400 | 1.44415300 | -1.67281900 |
| C | 1.59613900 | 2.29296800 | 0.07245600 |
| H | 1.42051300 | 2.54680000 | 1.11613800 |
| C | -0.95916100 | -0.23199500 | 0.03804100 |
| C | -0.64466100 | -0.84870000 | -1.32975200 |
| H | -0.84958300 | -1.91503300 | -1.36231200 |
| H | -1.25104700 | -0.35712400 | -2.09452500 |
| H | 0.40406800 | -0.71060300 | -1.59210200 |
| C | 0.02700500 | -0.64782100 | 1.14449500 |
| O | -0.26552800 | -0.47971200 | 2.31096200 |

| | | | |
|---|---|---|---|
| C | -5.89532500 | 0.07975800 | -0.97240800 |
| H | -6.55962300 | -0.62630700 | -1.47808000 |
| H | -6.44044600 | 1.01768200 | -0.83515200 |
| H | -5.05485700 | 0.27871400 | -1.63982500 |
| H | -6.35314300 | -0.75932800 | 0.92779200 |
| C | -1.60614900 | 2.84640700 | -2.03922000 |
| H | -2.53943200 | 3.25997500 | -2.42424400 |
| H | -0.93816300 | 3.67125900 | -1.77528700 |
| H | -1.11813400 | 2.29034300 | -2.84600600 |
| C | 2.91202700 | 2.61594400 | -0.46981200 |
| H | 3.09202000 | 2.35196000 | -1.51152900 |
| C | 3.88413100 | 3.20105200 | 0.23753200 |
| H | 3.68651000 | 3.45400400 | 1.27808000 |
| C | 5.24346700 | 3.53620300 | -0.29220400 |
| H | 6.02192500 | 3.02787000 | 0.28407500 |
| H | 5.34261700 | 3.24559100 | -1.33937000 |
| H | 5.44092600 | 4.60882000 | -0.20974700 |
| C | 1.40918100 | -1.11032300 | 0.83715600 |
| C | 1.87616000 | -2.14809700 | 0.11475500 |
| C | 2.58458800 | -0.47073500 | 1.49263200 |
| O | 1.18607100 | -3.05540900 | -0.56369100 |
| C | 3.38423000 | -2.22198900 | 0.17174200 |
| O | 2.59514000 | 0.40001000 | 2.33977700 |
| N | 3.70932400 | -1.04585200 | 0.94152700 |
| H | 1.80442500 | -3.56065300 | -1.12403600 |
| H | 3.68171800 | -3.14395700 | 0.68568200 |
| C | 3.99634700 | -2.21773700 | -1.22648700 |
| H | 4.59251000 | -0.96289300 | 1.42516600 |
| H | 3.64087400 | -1.34317300 | -1.78057100 |
| H | 5.08612200 | -2.18430600 | -1.15581200 |
| O | 3.55887300 | -3.43011500 | -1.83430200 |
| H | 3.79975700 | -3.43719500 | -2.76555400 |

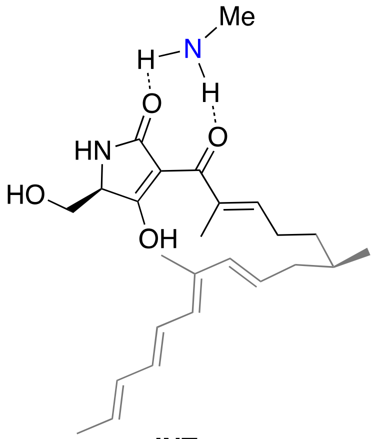


**INT$_{2b}$**

Energy (RM062X): -1385.678552 A.U.

Gibbs Free Energy: -1385.155776 A.U.

-----------------------------------------------------------

| | | | |
|---|---|---|---|
| C | 5.18754500 | 0.76639600 | 0.59788000 |
| C | 5.65289300 | 0.50687500 | -0.84228700 |
| C | 4.99302400 | -0.70673100 | -1.51788900 |
| H | 5.63278900 | -0.99417100 | -2.35664200 |
| H | 5.47439000 | -0.08030100 | 1.23274700 |
| H | 5.75123000 | 1.63371200 | 0.96705200 |
| H | 5.00129000 | -1.55645800 | -0.82390100 |
| C | 3.56935800 | -0.48985100 | -2.06952700 |
| H | 3.44016300 | 0.55171100 | -2.37630400 |
| H | 3.47072100 | -1.08055800 | -2.98982100 |
| C | 2.45175600 | -0.93317700 | -1.17800100 |
| H | 2.62591500 | -1.84061900 | -0.60435500 |
| C | 3.72510700 | 1.05277000 | 0.77189900 |
| H | 3.25705300 | 1.65099100 | -0.00861100 |
| C | 3.01213600 | 0.66578600 | 1.83711500 |
| H | 3.51833800 | 0.05220700 | 2.58185800 |
| C | 1.59236800 | 0.93287800 | 2.11604700 |
| C | 0.86831200 | 1.81501800 | 1.38929900 |
| H | 1.37375800 | 2.38213800 | 0.61103100 |
| C | -0.55464300 | 2.05701600 | 1.50221200 |
| H | -1.10754900 | 1.53406600 | 2.27636000 |
| C | -1.24753000 | 2.82175500 | 0.63481200 |
| H | -0.72058400 | 3.33404200 | -0.16939900 |
| C | 1.24024500 | -0.36393100 | -1.08182800 |
| C | 0.88529000 | 0.90982400 | -1.80974200 |
| H | 0.74720800 | 0.74121800 | -2.87955200 |
| H | -0.02864800 | 1.35307800 | -1.41651000 |
| H | 1.68545000 | 1.64482200 | -1.69217800 |
| C | 0.25901900 | -1.05149100 | -0.18719300 |
| O | 0.60327400 | -1.90947600 | 0.61382400 |
| C | 7.17349800 | 0.33069900 | -0.84355900 |
| H | 7.55888300 | 0.23961600 | -1.86140000 |
| H | 7.67018800 | 1.17762800 | -0.36408500 |
| H | 7.45100100 | -0.57590800 | -0.29602200 |
| H | 5.41127500 | 1.39586400 | -1.43949300 |
| C | 1.02358200 | 0.10969200 | 3.24047100 |
| H | 1.68774400 | 0.16077000 | 4.10811300 |
| H | 0.02635600 | 0.42043500 | 3.54095000 |
| H | 0.96025400 | -0.93626700 | 2.92544800 |
| C | -2.69464400 | 2.94873500 | 0.66381900 |
| H | -3.21062400 | 2.45267500 | 1.48516600 |
| C | -3.41332900 | 3.58768800 | -0.26912100 |
| H | -2.88708400 | 4.07118800 | -1.09088000 |
| C | -4.90791300 | 3.67172800 | -0.27810000 |
| H | -5.24485700 | 4.71221200 | -0.25457700 |
| H | -5.31732500 | 3.23024400 | -1.19211800 |
| H | -5.33712700 | 3.14927200 | 0.57880000 |
| C | -1.18103800 | -0.70452200 | -0.27732300 |
| C | -1.93587000 | -0.45320300 | -1.36460900 |
| O | -1.52629800 | -0.47054900 | -2.63069700 |
| H | -2.22565000 | -0.19751600 | -3.23744700 |
| C | -2.09158700 | -0.68660200 | 0.90554900 |
| O | -1.81878400 | -0.90828100 | 2.07196400 |
| N | -3.32629100 | -0.34331400 | 0.43590200 |
| H | -4.14144800 | -0.27422400 | 1.02596900 |
| C | -3.37444900 | -0.19353600 | -0.99565500 |
| H | -3.64831600 | 0.83296500 | -1.27682700 |
| C | -4.34374900 | -1.17392000 | -1.64402900 |

| | | | |
|---|---|---|---|
| H | -4.29304200 | -1.09114800 | -2.73536700 |
| H | -4.07228700 | -2.19305800 | -1.34986800 |
| O | -5.63094600 | -0.82427300 | -1.17037800 |
| H | -6.26341600 | -1.48386200 | -1.46958900 |
| N | -1.86970600 | -3.82246400 | 1.03308200 |
| H | -1.05750900 | -3.20915000 | 1.04009600 |
| C | -1.45092700 | -5.22005800 | 1.16334100 |
| H | -0.87765800 | -5.43687700 | 2.07287900 |
| H | -2.32805000 | -5.86976800 | 1.15550200 |
| H | -0.83451700 | -5.49329200 | 0.30489700 |
| H | -2.42503000 | -3.54563000 | 1.83601100 |

---

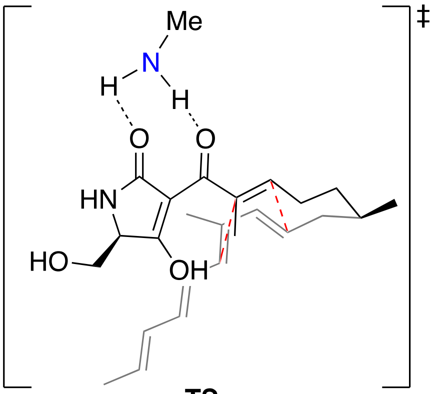

**TS*2b***

Energy (RM062X): -1385.658487 A.U.

Gibbs Free Energy: -1385.129396 A.U.

---

| | | | |
|---|---|---|---|
| C | -4.89692700 | -0.12854500 | 0.59817800 |
| C | -5.60782600 | -0.26148100 | -0.74785300 |
| C | -4.86969800 | 0.51517800 | -1.83561800 |
| H | -5.39594000 | 0.39459800 | -2.78739200 |
| H | -4.89544500 | 0.92525400 | 0.90507800 |
| H | -5.47504800 | -0.67107800 | 1.35525100 |
| H | -4.89595100 | 1.58523300 | -1.59159600 |
| C | -3.42171300 | 0.06578000 | -2.00182800 |
| H | -3.39560100 | -1.00605700 | -2.22371800 |
| H | -2.98111700 | 0.56615100 | -2.87209700 |
| C | -2.53535200 | 0.38248800 | -0.80901200 |
| H | -2.72428200 | 1.34652100 | -0.34371500 |
| C | -3.47311600 | -0.65555000 | 0.62666600 |
| H | -3.32625900 | -1.59340700 | 0.09288100 |
| C | -2.73985200 | -0.46091200 | 1.79973800 |
| H | -3.13644900 | 0.26088000 | 2.51151100 |
| C | -1.47092100 | -0.99459300 | 2.08188100 |
| C | -0.82755100 | -1.78349000 | 1.14567900 |
| H | -1.39970000 | -2.21483000 | 0.33341200 |
| C | 0.53955700 | -2.22281900 | 1.25810600 |
| H | 1.14535900 | -1.79872100 | 2.05442300 |
| C | 1.11710400 | -3.08228800 | 0.38981400 |
| H | 0.52032100 | -3.50739000 | -0.41600600 |
| C | -1.17706600 | 0.02125800 | -0.86492500 |
| C | -0.70287300 | -1.00338400 | -1.85738700 |
| H | -0.70481800 | -0.60549100 | -2.87701900 |
| H | 0.30786800 | -1.34346800 | -1.63155200 |
| H | -1.36052300 | -1.87866500 | -1.85464100 |
| C | -0.27268100 | 0.79161000 | -0.03437200 |
| O | -0.67819000 | 1.60485400 | 0.80925100 |
| C | -7.05843700 | 0.20126000 | -0.63320600 |
| H | -7.58181600 | 0.09021300 | -1.58585600 |
| H | -7.59916300 | -0.37395300 | 0.12237100 |
| H | -7.09979000 | 1.25700300 | -0.34707600 |
| H | -5.60165500 | -1.32352700 | -1.02843300 |
| C | -0.76739700 | -0.55839300 | 3.34322200 |
| H | -1.45229000 | -0.00970300 | 3.99086400 |
| H | -0.38578400 | -1.41713200 | 3.90140500 |
| H | 0.07889700 | 0.08703000 | 3.09731600 |
| C | 2.51305400 | -3.47046200 | 0.45331900 |
| H | 3.10565900 | -3.04345300 | 1.26150100 |
| C | 3.09538400 | -4.30256000 | -0.42173800 |
| H | 2.48835500 | -4.71818700 | -1.22447900 |
| C | 4.53373600 | -4.70975200 | -0.38261700 |
| H | 4.62768800 | -5.79552700 | -0.28836600 |
| H | 5.04218100 | -4.43040100 | -1.30998400 |
| H | 5.05539700 | -4.24237600 | 0.45389400 |
| C | 1.20074900 | 0.66287900 | -0.24370200 |
| C | 1.87682300 | 0.79827000 | -1.39885600 |
| O | 1.33992300 | 0.98151400 | -2.60733000 |
| H | 2.01513200 | 0.99488500 | -3.29643200 |
| C | 2.21442900 | 0.59652000 | 0.84097400 |
| O | 2.03795800 | 0.59727300 | 2.04756800 |
| N | 3.44503000 | 0.51286600 | 0.23509600 |
| H | 4.28055700 | 0.74701400 | 0.75392700 |
| C | 3.36682400 | 0.79853200 | -1.18067700 |
| H | 3.85289600 | 0.01658800 | -1.77576500 |
| C | 3.98563200 | 2.15421400 | -1.50883700 |
| H | 3.85953200 | 2.38053000 | -2.57328100 |
| H | 3.48095700 | 2.91832800 | -0.90793600 |
| O | 5.35999000 | 2.05225700 | -1.17410300 |
| H | 5.75714700 | 2.92648100 | -1.22190900 |
| N | 1.75565000 | 3.64477300 | 1.16018100 |
| H | 0.93948800 | 3.03618500 | 1.11460600 |
| C | 1.35395000 | 4.99711100 | 1.55632100 |
| H | 0.83746600 | 5.05089200 | 2.52243000 |
| H | 2.23365800 | 5.64127300 | 1.60885300 |
| H | 0.68943900 | 5.41334500 | 0.79693800 |
| H | 2.35644300 | 3.23907600 | 1.87094600 |

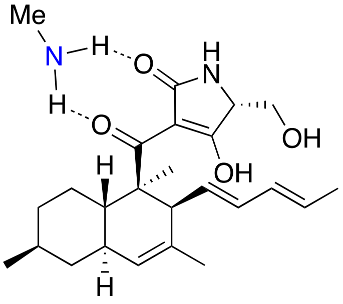

**PD*2b***

Energy (M062X): -1385.718527 A.U.
Gibbs Free Energy: -1385.185991 A.U.

------------------------------------------------------------

| | | | |
|---|---|---|---|
| C | 4.96075400 | 0.23850400 | 0.24555100 |
| C | 5.34852900 | -0.79287900 | -0.81345400 |
| C | 4.25109100 | -1.85400300 | -0.91595600 |
| H | 4.51478000 | -2.59138300 | -1.68059500 |
| H | 4.92899100 | -0.26294900 | 1.22341800 |
| H | 5.72772300 | 1.01763500 | 0.31648300 |
| H | 4.18988500 | -2.39152800 | 0.03972600 |
| C | 2.89838400 | -1.22300600 | -1.24327200 |
| H | 2.96706500 | -0.74999600 | -2.22867300 |
| H | 2.12346500 | -1.99334000 | -1.31545700 |
| C | 2.50034400 | -0.18152900 | -0.19066700 |
| H | 2.44346800 | -0.69510000 | 0.77492200 |
| C | 3.59720900 | 0.89039200 | -0.00650600 |
| H | 3.65745700 | 1.48517700 | -0.93257400 |
| C | 3.19874900 | 1.76494000 | 1.15138000 |
| H | 3.93301700 | 1.96771900 | 1.92852300 |
| C | 1.94783600 | 2.21127100 | 1.26505600 |
| C | 1.01891300 | 1.94777100 | 0.09291700 |
| H | 1.40289000 | 2.54687600 | -0.74247100 |
| C | -0.40558400 | 2.36816200 | 0.32881200 |
| H | -0.91223300 | 1.95023400 | 1.19665200 |
| C | -1.10005700 | 3.17577500 | -0.47956900 |
| H | -0.61817900 | 3.60363300 | -1.35852500 |
| C | 1.09946300 | 0.45087100 | -0.44935600 |
| C | 0.74926000 | 0.52579300 | -1.93873500 |
| H | 0.65857600 | -0.45614100 | -2.39758400 |
| H | -0.18985800 | 1.06429300 | -2.08152100 |
| H | 1.53482100 | 1.07471000 | -2.46439400 |
| C | 0.05885900 | -0.33250800 | 0.36638200 |
| O | 0.24175700 | -0.56167300 | 1.54495000 |
| C | 6.70523900 | -1.41837900 | -0.50188600 |
| H | 6.98955500 | -2.14535300 | -1.26695100 |
| H | 7.48859600 | -0.65809800 | -0.44613000 |
| H | 6.67168600 | -1.93849700 | 0.46083000 |
| H | 5.41253500 | -0.28034700 | -1.78327000 |
| C | 1.46043700 | 3.01717800 | 2.43502500 |
| H | 2.27318400 | 3.20215500 | 3.13882600 |
| H | 1.05159000 | 3.97822700 | 2.10942500 |
| H | 0.65781800 | 2.49496100 | 2.96429000 |
| C | -2.50160300 | 3.52067100 | -0.26044900 |
| H | -2.98190400 | 3.06852600 | 0.60618100 |
| C | -3.20146600 | 4.33145800 | -1.06114100 |
| H | -2.70113900 | 4.76712300 | -1.92486400 |
| C | -4.64053300 | 4.69332500 | -0.86293000 |
| H | -4.75902000 | 5.77525500 | -0.75472900 |
| H | -5.23993100 | 4.39616600 | -1.72820700 |
| H | -5.05087300 | 4.20974400 | 0.02503200 |
| C | -1.24550000 | -0.74264800 | -0.22946700 |
| C | -1.50730100 | -1.64214200 | -1.19350200 |
| O | -0.60837000 | -2.32547300 | -1.90202500 |
| H | -1.02649700 | -2.87031400 | -2.58038200 |
| C | -2.55096800 | -0.35377000 | 0.37436000 |
| O | -2.74523300 | 0.32849800 | 1.36432500 |
| N | -3.52811800 | -0.90480500 | -0.41494600 |
| H | -4.46647400 | -1.01564200 | -0.05485800 |
| C | -2.98795500 | -1.88048800 | -1.33761000 |
| H | -3.31669900 | -1.68569400 | -2.36481600 |
| C | -3.37510900 | -3.30063100 | -0.93337000 |
| H | -2.90431700 | -4.02693400 | -1.60482300 |
| H | -3.03577200 | -3.47147200 | 0.09358200 |
| O | -4.78832100 | -3.36319700 | -1.03194200 |
| H | -5.09097100 | -4.18664700 | -0.63858200 |
| N | -2.14756500 | -2.57363500 | 2.46878500 |
| H | -1.34957000 | -1.96090800 | 2.31682000 |
| C | -1.88925100 | -3.47214400 | 3.59730300 |
| H | -1.67555600 | -2.96002900 | 4.54302200 |
| H | -2.75323000 | -4.12068700 | 3.75283800 |
| H | -1.03770300 | -4.11271100 | 3.36132800 |
| H | -2.92450000 | -1.95671800 | 2.68467700 |

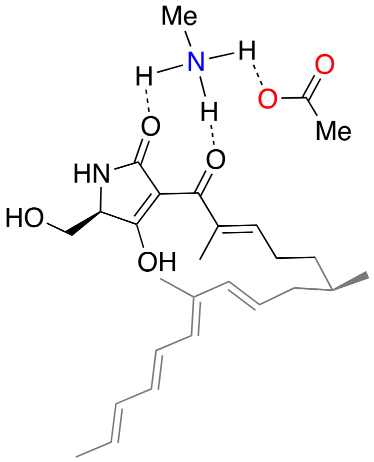


**INT$_{3b}$**

Energy (RM062X): -1614.779164 A.U.
Gibbs Free Energy: -1614.194361 A.U.

------------------------------------------------------------

| | | | |
|---|---|---|---|
| C | 5.41788800 | 0.71826500 | 0.40274700 |
| C | 5.83882200 | 0.28951200 | -1.01057200 |
| C | 5.09468800 | -0.93797600 | -1.56107700 |
| H | 5.68849200 | -1.32567700 | -2.39346400 |
| H | 5.60538500 | -0.10185300 | 1.10594200 |
| H | 6.08712500 | 1.53679600 | 0.70030400 |
| H | 5.08563600 | -1.72772600 | -0.79939900 |
| C | 3.66669000 | -0.69838400 | -2.08498500 |
| H | 3.59998400 | 0.28686100 | -2.55460000 |
| H | 3.47847700 | -1.42223200 | -2.88899600 |
| C | 2.57148900 | -0.89907400 | -1.08584000 |
| H | 2.71133300 | -1.71274500 | -0.37809300 |
| C | 4.00557400 | 1.20419900 | 0.55625100 |
| H | 3.60050900 | 1.77047000 | -0.28156700 |
| C | 3.28014900 | 1.03787400 | 1.66961500 |
| H | 3.72937800 | 0.45708300 | 2.47475500 |
| C | 1.91704400 | 1.51771500 | 1.94483500 |

| | | | |
|---|---|---|---|
| C | 1.24390800 | 2.31859100 | 1.08655400 |
| H | 1.75518100 | 2.66107000 | 0.19021700 |
| C | -0.13451200 | 2.74556700 | 1.20678500 |
| H | -0.69859700 | 2.43662800 | 2.08235400 |
| C | -0.78176400 | 3.43596300 | 0.24634400 |
| H | -0.24353900 | 3.73701000 | -0.65183800 |
| C | 1.41401800 | -0.22119500 | -1.02965700 |
| C | 1.12106400 | 0.94716000 | -1.93970500 |
| H | 0.91369600 | 0.62591800 | -2.96208200 |
| H | 0.26911100 | 1.52675500 | -1.58693200 |
| H | 1.98390400 | 1.61730200 | -1.97251000 |
| C | 0.45456700 | -0.65955800 | 0.02617700 |
| O | 0.82671700 | -1.34666300 | 0.97226000 |
| C | 7.34432500 | 0.01018800 | -1.01073400 |
| H | 7.70294500 | -0.21731300 | -2.01691100 |
| H | 7.90615400 | 0.86771100 | -0.63306600 |
| H | 7.57092600 | -0.84867700 | -0.37053500 |
| H | 5.64704600 | 1.13134200 | -1.68897700 |
| C | 1.33678200 | 1.00048000 | 3.23343900 |
| H | 2.05892900 | 1.13563000 | 4.04371600 |
| H | 0.40397800 | 1.48455900 | 3.51116100 |
| H | 1.13525100 | -0.07040800 | 3.13397300 |
| C | -2.19576700 | 3.76403300 | 0.31008200 |
| H | -2.72287800 | 3.47560700 | 1.21868000 |
| C | -2.87661600 | 4.35000800 | -0.68396100 |
| H | -2.33987400 | 4.62354500 | -1.59124600 |
| C | -4.34415900 | 4.64242100 | -0.65092100 |
| H | -4.53706800 | 5.70980200 | -0.79247400 |
| H | -4.86261300 | 4.11938600 | -1.46057000 |
| H | -4.78672100 | 4.33286400 | 0.29752300 |
| C | -0.96291900 | -0.24197500 | -0.05260900 |
| C | -1.74955300 | -0.04687000 | -1.13167600 |
| O | -1.42511000 | -0.28655900 | -2.39808800 |
| H | -2.10342100 | 0.03231600 | -3.00726100 |
| C | -1.80122100 | 0.03894500 | 1.14904400 |
| O | -1.50766500 | -0.11164700 | 2.32480800 |
| N | -2.98900900 | 0.52582900 | 0.69228500 |
| H | -3.78106600 | 0.68506800 | 1.29716000 |
| C | -3.11022100 | 0.47054500 | -0.74326700 |
| H | -3.24381500 | 1.47814100 | -1.16245400 |
| C | -4.26487000 | -0.41858600 | -1.18723700 |
| H | -4.26778400 | -0.52039100 | -2.27913000 |
| H | -4.14888000 | -1.40398700 | -0.72754500 |
| O | -5.44479200 | 0.23033400 | -0.74012000 |
| H | -6.18732500 | -0.37132600 | -0.84525400 |
| N | -1.12671700 | -2.90343300 | 2.29775700 |
| H | -0.39083200 | -2.28743900 | 1.91079700 |
| C | -0.61133600 | -4.26100100 | 2.57106400 |
| H | 0.21378500 | -4.21458500 | 3.27857800 |
| H | -1.41247100 | -4.87321600 | 2.98013800 |
| H | -0.27550600 | -4.67830500 | 1.62437300 |
| H | -1.49373900 | -2.44404900 | 3.12865800 |
| H | -1.90065400 | -2.95392300 | 1.53435300 |
| O | -2.87075500 | -3.11946300 | 0.35662200 |

| | | | |
|---|---|---|---|
| C | -2.15977300 | -3.42568200 | -0.65610900 |
| O | -0.93037000 | -3.60831200 | -0.62671700 |
| C | -2.88153900 | -3.55513600 | -1.99316000 |
| H | -2.53046300 | -4.44432900 | -2.51786900 |
| H | -3.96230700 | -3.59791900 | -1.86689600 |
| H | -2.62834700 | -2.69021600 | -2.61200100 |

------------------------------------------------------------

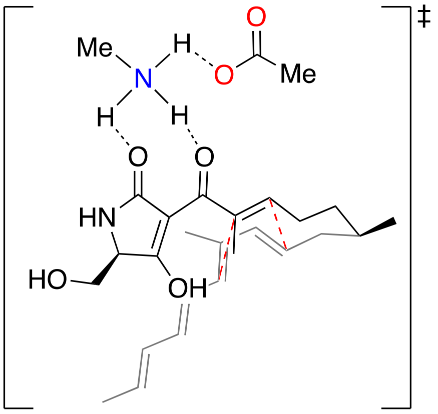


**TS**$_{3b}$

Energy (RM062X): -1614.759336 A.U.

Gibbs Free Energy: -1614.170988 A.U.

------------------------------------------------------------

| | | | |
|---|---|---|---|
| C | -5.17706100 | -0.26848900 | 0.38156400 |
| C | -5.82982600 | -0.25007800 | -0.99914700 |
| C | -5.00417900 | 0.57802000 | -1.97987600 |
| H | -5.49025200 | 0.57227400 | -2.96009800 |
| H | -5.14488800 | 0.75391100 | 0.77941200 |
| H | -5.81359100 | -0.84588300 | 1.06201600 |
| H | -4.98393500 | 1.62183900 | -1.64058600 |
| C | -3.57842800 | 0.05792100 | -2.12380900 |
| H | -3.60456600 | -0.99076700 | -2.43562400 |
| H | -3.06841800 | 0.60134400 | -2.92777100 |
| C | -2.72693400 | 0.22313300 | -0.87602100 |
| H | -2.86763400 | 1.16487000 | -0.35238000 |
| C | -3.77872800 | -0.86203600 | 0.43506900 |
| H | -3.64555400 | -1.77292500 | -0.14762700 |
| C | -3.10751000 | -0.77482100 | 1.65760000 |
| H | -3.51026300 | -0.06828200 | 2.38130800 |
| C | -1.90032200 | -1.40724900 | 2.00761700 |
| C | -1.23421100 | -2.19770400 | 1.09058400 |
| H | -1.75726800 | -2.54073900 | 0.20605900 |
| C | 0.10897500 | -2.68114400 | 1.26530100 |
| H | 0.65929700 | -2.34674800 | 2.14080100 |
| C | 0.75806200 | -3.42293000 | 0.33959600 |
| H | 0.22284100 | -3.75871800 | -0.54711800 |
| C | -1.39555400 | -0.23586100 | -0.90753700 |
| C | -0.99031400 | -1.27808400 | -1.91159800 |
| H | -0.89857800 | -0.85850800 | -2.91805500 |
| H | -0.03843700 | -1.74034900 | -1.65480700 |
| H | -1.74455600 | -2.07019000 | -1.96327300 |
| C | -0.48029700 | 0.39569200 | 0.01139200 |

| | | | |
|---|---|---|---|
| O | -0.88991800 | 1.18635100 | 0.88871200 |
| C | -7.26015100 | 0.27593800 | -0.90403600 |
| H | -7.74314600 | 0.27642200 | -1.88405500 |
| H | -7.86256300 | -0.33450900 | -0.22686700 |
| H | -7.26369700 | 1.30314000 | -0.52586800 |
| H | -5.86168300 | -1.28258900 | -1.37256800 |
| C | -1.27678300 | -1.06494600 | 3.33852900 |
| H | -2.00354900 | -0.56657400 | 3.98107800 |
| H | -0.92342900 | -1.95999400 | 3.85564400 |
| H | -0.42243200 | -0.39968600 | 3.19151100 |
| C | 2.16721900 | -3.75021800 | 0.42346600 |
| H | 2.69476900 | -3.42932800 | 1.32026700 |
| C | 2.84411900 | -4.36835200 | -0.55535800 |
| H | 2.30254600 | -4.66960100 | -1.45074400 |
| C | 4.30913900 | -4.66475100 | -0.51830700 |
| H | 4.49346500 | -5.73780300 | -0.62399100 |
| H | 4.82367200 | -4.17476900 | -1.35050800 |
| H | 4.75852000 | -4.32436100 | 0.41588100 |
| C | 0.97940100 | 0.11592600 | -0.08494600 |
| C | 1.76551700 | 0.01494500 | -1.17583800 |
| O | 1.39679200 | 0.23402300 | -2.43884600 |
| H | 2.10261400 | 0.01370000 | -3.05910600 |
| C | 1.87091600 | -0.06620200 | 1.09478400 |
| O | 1.60091200 | 0.08817200 | 2.27786600 |
| N | 3.08717900 | -0.45811000 | 0.62046800 |
| H | 3.91088500 | -0.47091800 | 1.20445300 |
| C | 3.17629400 | -0.37649400 | -0.81711000 |
| H | 3.39827600 | -1.36217600 | -1.25082200 |
| C | 4.23684500 | 0.62190900 | -1.26099400 |
| H | 4.22873700 | 0.72525900 | -2.35263300 |
| H | 4.02479600 | 1.58946400 | -0.79695000 |
| O | 5.47679300 | 0.09113600 | -0.81795300 |
| H | 6.15481700 | 0.76550500 | -0.91640200 |
| N | 0.91318100 | 2.81317700 | 2.23797700 |
| H | 0.22135600 | 2.17951800 | 1.78329900 |
| C | 0.29827200 | 4.09256400 | 2.64788700 |
| H | -0.52962300 | 3.90634700 | 3.32887700 |
| H | 1.04646700 | 4.71045900 | 3.14055100 |
| H | -0.05308300 | 4.59074500 | 1.74750200 |
| H | 1.30686700 | 2.29438300 | 3.02072600 |
| H | 1.68896500 | 2.98337800 | 1.50689600 |
| O | 2.71172300 | 3.22439200 | 0.34171400 |
| C | 2.03869000 | 3.68845200 | -0.63343700 |
| O | 0.85805700 | 4.07637200 | -0.55883400 |
| C | 2.73405600 | 3.73482500 | -1.98984500 |
| H | 2.37023300 | 4.57830900 | -2.57576200 |
| H | 3.81696000 | 3.78698600 | -1.88277400 |
| H | 2.48440900 | 2.81874800 | -2.53346600 |

-------------------------------------------------------------

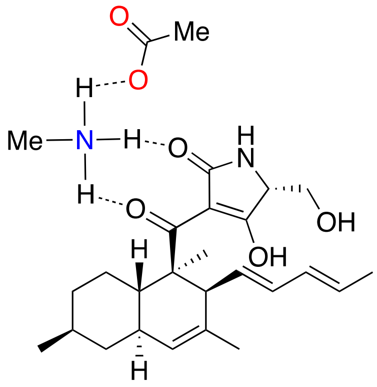

**PD$_{3b}$**

Energy (RM062X): -1614.815821 A.U.

Gibbs Free Energy: -1614.22358 A.U.

-------------------------------------------------------------

| | | | |
|---|---|---|---|
| C | 5.14748100 | 0.19707500 | 0.06126700 |
| C | 5.43312000 | -0.79037700 | -1.06969600 |
| C | 4.24468800 | -1.74119100 | -1.22797500 |
| H | 4.43555300 | -2.44522400 | -2.04386600 |
| H | 5.07897400 | -0.36512600 | 1.00356500 |
| H | 5.98155000 | 0.89847600 | 0.17344400 |
| H | 4.14244400 | -2.33454000 | -0.30959800 |
| C | 2.95226400 | -0.97073300 | -1.49412300 |
| H | 3.05682900 | -0.43933100 | -2.44592200 |
| H | 2.11007900 | -1.66226200 | -1.60784900 |
| C | 2.65762100 | 0.02695700 | -0.36779500 |
| H | 2.56240800 | -0.54490400 | 0.56115700 |
| C | 3.84531100 | 0.98309300 | -0.12615800 |
| H | 3.95100000 | 1.63477200 | -1.00868500 |
| C | 3.53695300 | 1.80382000 | 1.09712800 |
| H | 4.30004600 | 1.89484100 | 1.86758300 |
| C | 2.32852000 | 2.33792100 | 1.27452500 |
| C | 1.35358100 | 2.23950900 | 0.11471900 |
| H | 1.75429000 | 2.88667700 | -0.67520300 |
| C | -0.03373600 | 2.72795500 | 0.43177700 |
| H | -0.53174800 | 2.28767700 | 1.29361500 |
| C | -0.71128200 | 3.61681500 | -0.30240900 |
| H | -0.23840400 | 4.07113200 | -1.17282800 |
| C | 1.31751100 | 0.79517400 | -0.56632600 |
| C | 0.99365900 | 1.04249000 | -2.04195300 |
| H | 0.84502100 | 0.12287900 | -2.60060200 |
| H | 0.09609000 | 1.65775100 | -2.13748700 |
| H | 1.82380900 | 1.58597800 | -2.50053400 |
| C | 0.21378000 | 0.03317100 | 0.18556800 |
| O | 0.39106600 | -0.27942300 | 1.34858800 |
| C | 6.73090100 | -1.55324900 | -0.81908500 |
| H | 6.94371200 | -2.24821700 | -1.63537000 |
| H | 7.57926000 | -0.87084600 | -0.72234400 |
| H | 6.65788500 | -2.13319400 | 0.10653000 |
| H | 5.53502200 | -0.22088700 | -2.00381700 |
| C | 1.93561900 | 3.09606200 | 2.51049200 |
| H | 2.77796100 | 3.16892300 | 3.19981800 |
| H | 1.59565900 | 4.10598300 | 2.26262000 |

| | | | |
|---|---|---|---|
| H | 1.10816100 | 2.60189500 | 3.02826200 |
| C | -2.08516000 | 4.01656300 | -0.01269900 |
| H | -2.55784500 | 3.54225400 | 0.84646300 |
| C | -2.77119700 | 4.90173000 | -0.74352500 |
| H | -2.27975300 | 5.35912900 | -1.60105500 |
| C | -4.18296200 | 5.31981800 | -0.47362800 |
| H | -4.24378500 | 6.39832700 | -0.30206600 |
| H | -4.82494700 | 5.10108600 | -1.33167900 |
| H | -4.58663000 | 4.80685500 | 0.40078000 |
| C | -1.13014900 | -0.18972300 | -0.41274500 |
| C | -1.52760100 | -0.81336200 | -1.53959000 |
| O | -0.75490100 | -1.46284200 | -2.40652000 |
| H | -1.25563800 | -1.77092600 | -3.17283300 |
| C | -2.36731000 | 0.24167200 | 0.30293800 |
| O | -2.45279200 | 0.71314400 | 1.42592100 |
| N | -3.41037200 | 0.02410600 | -0.54835200 |
| H | -4.36783200 | 0.03848400 | -0.22567600 |
| C | -3.02347500 | -0.74101600 | -1.71180300 |
| H | -3.25797200 | -0.19840000 | -2.63654300 |
| C | -3.70445900 | -2.10439900 | -1.74544300 |
| H | -3.33167100 | -2.69117000 | -2.59354300 |
| H | -3.48995100 | -2.62922400 | -0.81018700 |
| O | -5.09045400 | -1.83722700 | -1.88962100 |
| H | -5.58020200 | -2.64893700 | -1.72965300 |
| N | -1.66104900 | -1.49681000 | 3.01503000 |
| H | -0.85868000 | -1.06293000 | 2.53832600 |
| C | -1.25633700 | -2.23770000 | 4.22797800 |
| H | -0.75714300 | -1.57011000 | 4.92683400 |
| H | -2.14184800 | -2.66335700 | 4.69522200 |
| H | -0.58419100 | -3.03387500 | 3.91714100 |
| H | -2.32282000 | -0.74673200 | 3.20742500 |
| H | -2.08569400 | -2.17932800 | 2.27391100 |
| O | -2.47553300 | -3.14687200 | 1.17500500 |
| C | -1.39284000 | -3.72143200 | 0.81911000 |
| O | -0.31873800 | -3.64316700 | 1.43909700 |
| C | -1.43970000 | -4.52282200 | -0.47540100 |
| H | -0.73515100 | -5.35269700 | -0.43580500 |
| H | -2.44527700 | -4.88817300 | -0.68101700 |
| H | -1.13662700 | -3.86562600 | -1.29593500 |

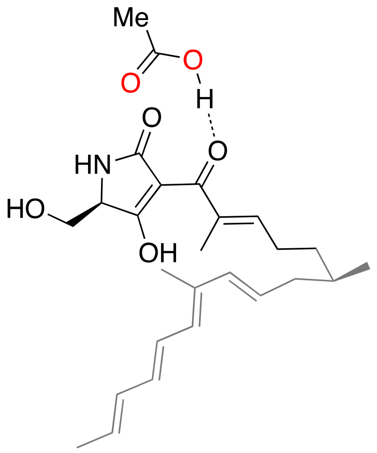

**INT$_{4b}$**

Energy (RM062X): -1518.921987 A.U.

Gibbs Free Energy: -1518.399435 A.U.

---

| | | | |
|---|---|---|---|
| C | 5.14774800 | -0.35763800 | -0.84981400 |
| C | 5.71292600 | -0.68711900 | 0.53960600 |
| C | 5.08041800 | 0.10513400 | 1.69477300 |
| H | 5.76123500 | 0.02938300 | 2.54712400 |
| H | 5.30191800 | 0.70636600 | -1.06481400 |
| H | 5.75462900 | -0.90998800 | -1.57994000 |
| H | 5.04329300 | 1.16864900 | 1.42749800 |
| C | 3.69396600 | -0.36499000 | 2.16931700 |
| H | 3.62624100 | -1.45459600 | 2.10824300 |
| H | 3.60444700 | -0.12287600 | 3.23679800 |
| C | 2.52559700 | 0.28945700 | 1.50332500 |
| H | 2.63817500 | 1.34363900 | 1.26152300 |
| C | 3.71077000 | -0.71948200 | -1.09038700 |
| H | 3.37158900 | -1.65559700 | -0.64769800 |
| C | 2.88923900 | -0.00386400 | -1.86920500 |
| H | 3.28314000 | 0.92253500 | -2.28637500 |
| C | 1.49490500 | -0.29461500 | -2.23636100 |
| C | 0.84534800 | -1.40274400 | -1.81012600 |
| H | 1.39071200 | -2.12145500 | -1.20353900 |
| C | -0.54751000 | -1.72326100 | -2.04231300 |
| H | -1.13210500 | -1.05981500 | -2.67163000 |
| C | -1.18407400 | -2.75156400 | -1.44666800 |
| H | -0.62962900 | -3.42080000 | -0.78986700 |
| C | 1.33607200 | -0.28193000 | 1.25345800 |
| C | 1.07641200 | -1.74528000 | 1.51870400 |
| H | 0.98447300 | -1.95722100 | 2.58550100 |
| H | 0.16689600 | -2.08679600 | 1.02681600 |
| H | 1.90510900 | -2.34413600 | 1.13213900 |
| C | 0.29976600 | 0.60135400 | 0.65032300 |
| O | 0.60533800 | 1.67471600 | 0.13281600 |
| C | 7.22423300 | -0.44122700 | 0.52550200 |
| H | 7.68071500 | -0.73810400 | 1.47220000 |
| H | 7.70880900 | -1.00067100 | -0.27811500 |
| H | 7.43431400 | 0.62194600 | 0.36938100 |
| H | 5.54162200 | -1.75491300 | 0.72960700 |

| | | | |
|---|---|---|---|
| C | 0.85778800 | 0.77556700 | -3.08089200 |
| H | 1.51188000 | 1.02203700 | -3.92226200 |
| H | -0.12160100 | 0.50100500 | -3.46280000 |
| H | 0.73253600 | 1.68009200 | -2.47749800 |
| C | -2.61014700 | -2.99465400 | -1.58637400 |
| H | -3.15269300 | -2.33766300 | -2.26607600 |
| C | -3.28217100 | -3.93854700 | -0.91342400 |
| H | -2.72842500 | -4.58444800 | -0.23338300 |
| C | -4.75810300 | -4.16646100 | -1.01237900 |
| H | -4.97763400 | -5.18899200 | -1.33286300 |
| H | -5.23835500 | -4.03283800 | -0.03801900 |
| H | -5.21762200 | -3.47505600 | -1.72077600 |
| C | -1.11665100 | 0.19030400 | 0.66537500 |
| C | -1.81309000 | -0.49528600 | 1.59638800 |
| O | -1.36514700 | -0.90419700 | 2.77507300 |
| H | -2.04381200 | -1.38394600 | 3.26695100 |
| C | -2.06754100 | 0.53565800 | -0.42603200 |
| O | -1.83302100 | 1.12460100 | -1.46443400 |
| N | -3.29066300 | 0.03773700 | -0.05853100 |
| H | -4.01974500 | -0.09123200 | -0.74503800 |
| C | -3.24330300 | -0.72444900 | 1.16707100 |
| H | -3.37880600 | -1.79724400 | 0.96876100 |
| C | -4.26324900 | -0.28604100 | 2.21016900 |
| H | -5.26271000 | -0.36383500 | 1.77207100 |
| H | -4.22247900 | -0.96952000 | 3.06733200 |
| O | -3.96319400 | 1.03821200 | 2.59756900 |
| H | -4.64488600 | 1.34761300 | 3.20066600 |
| O | -2.18461500 | 3.64538100 | 0.50688000 |
| C | -1.69073500 | 4.19458800 | -0.45381600 |
| O | -0.59139500 | 3.75620000 | -1.05188600 |
| C | -2.24405800 | 5.43277500 | -1.10156800 |
| H | -0.30871300 | 2.90089000 | -0.63869600 |
| H | -3.07387500 | 5.82124800 | -0.51743800 |
| H | -1.46000600 | 6.18488600 | -1.19380300 |
| H | -2.58503600 | 5.18388200 | -2.10831900 |

------------------------------------------------------------

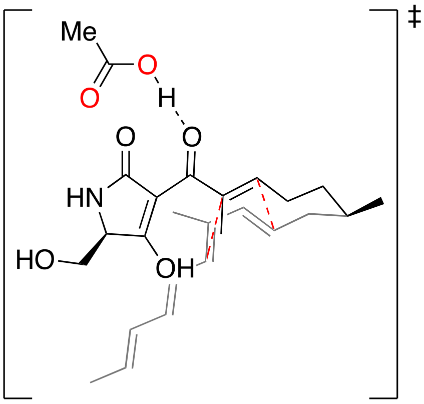


**TS$_{4b}$**

Energy (RM062X): -1518.903084 A.U.
Gibbs Free Energy: -1518.377177 A.U.

------------------------------------------------------------

| | | | |
|---|---|---|---|
| C | 4.94626400 | 0.04079400 | -0.58330300 |
| C | 5.70597700 | -0.68542000 | 0.52457900 |
| C | 5.00783500 | -0.50317400 | 1.86917200 |
| H | 5.56816700 | -1.03289200 | 2.64538900 |
| H | 4.94161400 | 1.11762600 | -0.37132000 |
| H | 5.49296500 | -0.08712400 | -1.52472300 |
| H | 5.01827300 | 0.56090000 | 2.13871000 |
| C | 3.57219700 | -1.01473300 | 1.85063000 |
| H | 3.56669000 | -2.06794000 | 1.55371600 |
| H | 3.15759800 | -0.98301900 | 2.86501000 |
| C | 2.63037500 | -0.20832700 | 0.97186900 |
| H | 2.77099800 | 0.86787100 | 1.02274300 |
| C | 3.51604800 | -0.42195000 | -0.81067900 |
| H | 3.37612800 | -1.50247500 | -0.80217300 |
| C | 2.75823100 | 0.32145100 | -1.71852100 |
| H | 3.14052500 | 1.30947900 | -1.96936800 |
| C | 1.49940500 | -0.01229300 | -2.25319900 |
| C | 0.86046600 | -1.17553400 | -1.86982900 |
| H | 1.41934400 | -1.94607500 | -1.35260500 |
| C | -0.51530400 | -1.46559800 | -2.16832900 |
| H | -1.08403100 | -0.71152600 | -2.70321000 |
| C | -1.17289700 | -2.54630700 | -1.69159600 |
| H | -0.62732900 | -3.30789600 | -1.13722400 |
| C | 1.28665100 | -0.62332600 | 0.87960400 |
| C | 0.93089100 | -2.05123900 | 1.18593100 |
| H | 0.96750700 | -2.25879600 | 2.25950700 |
| H | -0.06628200 | -2.30497600 | 0.83117800 |
| H | 1.64239200 | -2.73101000 | 0.70513100 |
| C | 0.32197300 | 0.38611100 | 0.53196000 |
| O | 0.69423700 | 1.52994600 | 0.17548900 |
| C | 7.15188900 | -0.19841100 | 0.58451700 |
| H | 7.71078300 | -0.72595700 | 1.36105200 |
| H | 7.66216800 | -0.35442700 | -0.36909500 |
| H | 7.18415700 | 0.87143900 | 0.81361400 |
| H | 5.70837500 | -1.75797300 | 0.28756000 |
| C | 0.80071800 | 1.01370300 | -3.11176800 |
| H | 1.50305200 | 1.78632000 | -3.42689200 |
| H | 0.37076100 | 0.55961100 | -4.00749500 |
| H | -0.00932200 | 1.48269900 | -2.54749200 |
| C | -2.60819400 | -2.71195000 | -1.80791700 |
| H | -3.14451000 | -1.95990700 | -2.38573000 |
| C | -3.29511900 | -3.69406400 | -1.20566700 |
| H | -2.74366200 | -4.43140300 | -0.62417000 |
| C | -4.78133300 | -3.84856300 | -1.26077800 |
| H | -5.05718700 | -4.81667500 | -1.68887600 |
| H | -5.21159900 | -3.81708100 | -0.25514900 |
| H | -5.23957100 | -3.05970100 | -1.85929400 |
| C | -1.13042600 | 0.09201800 | 0.62238100 |
| C | -1.80480700 | -0.58146600 | 1.57637700 |
| O | -1.29547000 | -1.08468800 | 2.69851900 |
| H | -1.97141500 | -1.52552300 | 3.22782600 |
| C | -2.14185800 | 0.58025600 | -0.35207900 |
| O | -1.95875300 | 1.21060500 | -1.37837000 |
| N | -3.37157000 | 0.17283200 | 0.10153300 |

| | | | |
|---|---|---|---|
| H | -4.15384900 | 0.12605000 | -0.53543000 |
| C | -3.27865200 | -0.68362900 | 1.26081200 |
| H | -3.50522300 | -1.72650700 | 0.99541200 |
| C | -4.18520300 | -0.26996800 | 2.41222700 |
| H | -5.21540200 | -0.23348600 | 2.04537600 |
| H | -4.14277300 | -1.03177300 | 3.20067200 |
| O | -3.75641300 | 0.98731700 | 2.89216400 |
| H | -4.37204600 | 1.28897000 | 3.56602700 |
| O | -1.96499800 | 3.56447400 | 0.77614700 |
| C | -1.48692600 | 4.08931900 | -0.20878600 |
| O | -0.39272400 | 3.65440300 | -0.80641700 |
| C | -2.07573400 | 5.29816900 | -0.88394900 |
| H | -0.09248800 | 2.78250800 | -0.40003900 |
| H | -2.88164700 | 5.71009100 | -0.28229500 |
| H | -1.30205500 | 6.04855200 | -1.04820800 |
| H | -2.46197200 | 5.00053300 | -1.86105900 |

-------------------------------------------------------------

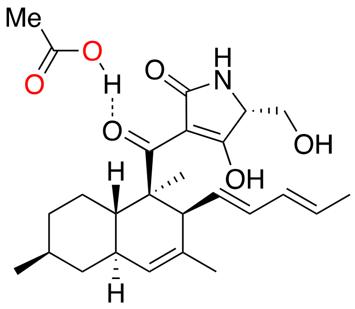


**PD$_{4b}$**

Energy (RM062X): -1518.957857 A.U.
Gibbs Free Energy: -1518.428537 A.U.

-------------------------------------------------------------

| | | | |
|---|---|---|---|
| C | 4.93279800 | -0.29553100 | -0.67861800 |
| C | 5.50854000 | 0.56523300 | 0.44505100 |
| C | 4.47569700 | 1.61395400 | 0.86235500 |
| H | 4.87405300 | 2.22750900 | 1.67642800 |
| H | 4.77402600 | 0.34327400 | -1.55915200 |
| H | 5.65316900 | -1.06599000 | -0.97472100 |
| H | 4.29352500 | 2.28684800 | 0.01381300 |
| C | 3.16403700 | 0.96054700 | 1.29607600 |
| H | 3.35990500 | 0.34639800 | 2.18144300 |
| H | 2.43716700 | 1.72311900 | 1.59380600 |
| C | 2.57886500 | 0.09027500 | 0.17743700 |
| H | 2.40211600 | 0.73996900 | -0.68651100 |
| C | 3.60018600 | -0.95981800 | -0.31792900 |
| H | 3.77514000 | -1.67910900 | 0.49863000 |
| C | 3.00615600 | -1.65589200 | -1.51232400 |
| H | 3.60247400 | -1.73868000 | -2.41871000 |
| C | 1.74091000 | -2.07443000 | -1.48818100 |
| C | 1.01952100 | -1.97571100 | -0.15602600 |
| H | 1.52874400 | -2.67048000 | 0.52313600 |
| C | -0.42846400 | -2.38189600 | -0.20768200 |
| H | -1.07151300 | -1.85717000 | -0.91238300 |
| C | -0.97695600 | -3.31599400 | 0.57640000 |
| H | -0.35421800 | -3.85160400 | 1.29266100 |
| C | 1.20766000 | -0.55784900 | 0.55056200 |
| C | 1.06181700 | -0.81714600 | 2.05245400 |
| H | 1.09074700 | 0.09439600 | 2.64291600 |
| H | 0.12468000 | -1.33801600 | 2.25935000 |
| H | 1.88196600 | -1.45919800 | 2.38379900 |
| C | 0.08765800 | 0.31201300 | -0.03305200 |
| O | 0.09106500 | 0.55789800 | -1.22839000 |
| C | 6.82484000 | 1.21237400 | 0.02367600 |
| H | 7.24449100 | 1.81561200 | 0.83276300 |
| H | 7.56435000 | 0.45855200 | -0.25852200 |
| H | 6.66682600 | 1.86835700 | -0.83839800 |
| H | 5.69777800 | -0.08409100 | 1.31098900 |
| C | 1.04373500 | -2.67829500 | -2.67278500 |
| H | 1.72592800 | -2.75768400 | -3.52039500 |
| H | 0.65382100 | -3.67345500 | -2.44000200 |
| H | 0.18895000 | -2.06420000 | -2.97307700 |
| C | -2.39270800 | -3.67011100 | 0.53960200 |
| H | -3.01242500 | -3.12506100 | -0.17119600 |
| C | -2.94665300 | -4.60215600 | 1.32226200 |
| H | -2.30921200 | -5.13350000 | 2.02749400 |
| C | -4.39522600 | -4.97924100 | 1.30725600 |
| H | -4.51880000 | -6.04090600 | 1.07530400 |
| H | -4.85024300 | -4.81643500 | 2.28843200 |
| H | -4.94782100 | -4.39682000 | 0.56829500 |
| C | -1.07873700 | 0.75244200 | 0.76604100 |
| C | -1.16666000 | 1.47065700 | 1.90268700 |
| O | -0.15866600 | 1.97565000 | 2.60263200 |
| H | -0.46873100 | 2.47965700 | 3.36570600 |
| C | -2.46726100 | 0.54934900 | 0.26111000 |
| O | -2.81075300 | -0.06072800 | -0.73529100 |
| N | -3.30198700 | 1.16451700 | 1.14997000 |
| H | -4.29514000 | 0.98412000 | 1.15061000 |
| C | -2.60728700 | 1.73106000 | 2.28295800 |
| H | -2.83527400 | 1.18185500 | 3.20522800 |
| C | -2.90299500 | 3.20820000 | 2.51741500 |
| H | -3.98433700 | 3.33175800 | 2.62599000 |
| H | -2.43766900 | 3.52598100 | 3.45885700 |
| O | -2.39779300 | 3.94066300 | 1.42209300 |
| H | -2.66854200 | 4.85901100 | 1.50888900 |
| O | -2.48875300 | 2.80823200 | -2.17782300 |
| C | -2.46809600 | 2.09717900 | -3.15695000 |
| O | -1.65677700 | 1.05017600 | -3.27158100 |
| C | -3.33698300 | 2.28297600 | -4.36769200 |
| H | -1.16391500 | 0.92571000 | -2.42931700 |
| H | -3.91700600 | 3.19683800 | -4.27346600 |
| H | -2.71765600 | 2.31777600 | -5.26453700 |
| H | -4.00641500 | 1.42577600 | -4.45968900 |

-------------------------------------------------------------

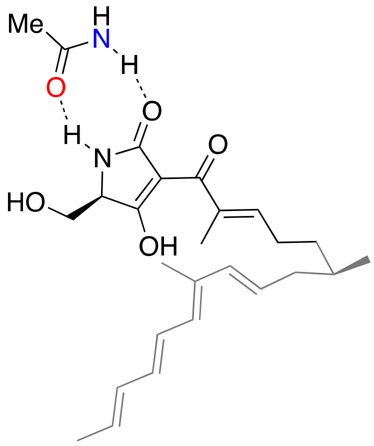


**INT$_{5b}$**

Energy (RM062X): -1499.051390 A.U.
Gibbs Free Energy: -1498.516692 A.U.

-------------------------------------------------------------

| | | | |
|---|---|---|---|
| C | 5.34675000 | -1.64461500 | 0.18384600 |
| C | 6.14253700 | -0.41318500 | 0.63998200 |
| C | 5.81071700 | 0.88201600 | -0.11916400 |
| H | 6.64474700 | 1.57054900 | 0.04198000 |
| H | 5.57198700 | -1.85660700 | -0.86806400 |
| H | 5.72503900 | -2.50097200 | 0.75833400 |
| H | 5.79053300 | 0.67515200 | -1.19650300 |
| C | 4.51921200 | 1.60969300 | 0.30136600 |
| H | 4.33521800 | 1.45948300 | 1.36874800 |
| H | 4.68681400 | 2.68820500 | 0.18131000 |
| C | 3.30094900 | 1.29158100 | -0.50808700 |
| H | 3.45742400 | 1.14438000 | -1.57435100 |
| C | 3.85995600 | -1.58158000 | 0.37858900 |
| H | 3.51590400 | -1.08844300 | 1.28688600 |
| C | 2.97820700 | -2.14836200 | -0.45466700 |
| H | 3.37481300 | -2.61788300 | -1.35434000 |
| C | 1.51318500 | -2.19853300 | -0.32756400 |
| C | 0.86561900 | -1.75926200 | 0.77675600 |
| H | 1.45467500 | -1.39733900 | 1.61587500 |
| C | -0.56945900 | -1.67174300 | 0.94132800 |
| H | -1.20188200 | -2.05650400 | 0.14767900 |
| C | -1.17507300 | -1.05878000 | 1.97827700 |
| H | -0.57666900 | -0.63738500 | 2.78513200 |
| C | 2.03836100 | 1.24698300 | -0.05699600 |
| C | 1.69966000 | 1.41145100 | 1.40509900 |
| H | 1.80359000 | 2.44784400 | 1.73234600 |
| H | 0.68076000 | 1.09112200 | 1.61839600 |
| H | 2.37140600 | 0.80360700 | 2.01636100 |
| C | 0.97870400 | 0.99420600 | -1.08532500 |
| O | 1.24892200 | 0.58007500 | -2.20205300 |
| C | 7.63708300 | -0.71583300 | 0.50274400 |
| H | 8.24267100 | 0.10161200 | 0.90005300 |
| H | 7.90623700 | -1.63115000 | 1.03520900 |
| H | 7.90214000 | -0.84957900 | -0.55113800 |
| H | 5.92411500 | -0.24539600 | 1.70282500 |
| C | 0.80868100 | -2.73026800 | -1.54788200 |
| H | 1.33385700 | -3.61128100 | -1.92620200 |
| H | -0.22990600 | -2.99586800 | -1.36380700 |
| H | 0.81645800 | -1.96520200 | -2.33039900 |
| C | -2.61428900 | -0.86907000 | 2.04083200 |
| H | -3.19330900 | -1.29978000 | 1.22442300 |
| C | -3.25344100 | -0.17082300 | 2.98885700 |
| H | -2.67098300 | 0.26991700 | 3.79677400 |
| C | -4.73192200 | 0.06584400 | 3.00313200 |
| H | -5.18785100 | -0.33400700 | 3.91375800 |
| H | -4.95487000 | 1.13745400 | 2.98550000 |
| H | -5.20851300 | -0.39946300 | 2.13787100 |
| C | -0.43941200 | 1.26332400 | -0.74187900 |
| C | -0.97426000 | 2.26651800 | -0.01672600 |
| O | -0.31721800 | 3.28820200 | 0.52088300 |
| H | -0.91055800 | 3.87451500 | 1.00689800 |
| C | -1.57493100 | 0.43417100 | -1.22391800 |
| O | -1.52618100 | -0.58472600 | -1.90611500 |
| N | -2.72141300 | 0.99142100 | -0.74931900 |
| H | -3.59459300 | 0.45947100 | -0.73608000 |
| C | -2.47538500 | 2.14028200 | 0.08684900 |
| H | -2.74214500 | 1.92880700 | 1.13187500 |
| C | -3.21190900 | 3.39616100 | -0.36469400 |
| H | -4.28445700 | 3.18150600 | -0.38289100 |
| H | -3.04410000 | 4.19901100 | 0.36391100 |
| O | -2.72952100 | 3.75616400 | -1.64308900 |
| H | -3.25203400 | 4.49075300 | -1.97674800 |
| C | -4.78951100 | -2.02576500 | -0.80194100 |
| C | -5.81550600 | -3.02773400 | -0.33737100 |
| H | -6.79689900 | -2.71721900 | -0.69394100 |
| H | -5.60272800 | -4.04136100 | -0.67281900 |
| H | -5.83279800 | -3.00975600 | 0.75481000 |
| O | -4.99366200 | -0.81392900 | -0.68599000 |
| N | -3.64325000 | -2.50939500 | -1.30586800 |
| H | -2.91146400 | -1.86688900 | -1.61621800 |
| H | -3.49585900 | -3.50185500 | -1.39489900 |

-------------------------------------------------------------

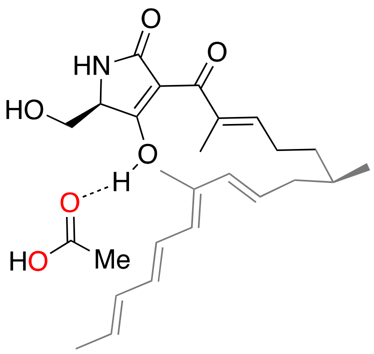


**INT$_{6b}$**

Energy (RM062X): -1518.918413 A.U.
Gibbs Free Energy: -1518.392945 A.U.

-------------------------------------------------------------

| | | | |
|---|---|---|---|
| C | 5.48173900 | 0.48983700 | 0.91481700 |
| C | 5.64577700 | 1.65423200 | -0.07348800 |

| | | | |
|---|---|---|---|
| C | 5.02529000 | 1.40344000 | -1.45921500 |
| H | 5.51902500 | 2.08164300 | -2.16020900 |
| H | 6.05219300 | -0.37539400 | 0.55677700 |
| H | 5.94812900 | 0.79845500 | 1.85994500 |
| H | 5.27900500 | 0.38846900 | -1.78927000 |
| C | 3.50163400 | 1.63231500 | -1.57446300 |
| H | 3.16781000 | 2.33732900 | -0.80847100 |
| H | 3.30574500 | 2.12941600 | -2.53340300 |
| C | 2.65851900 | 0.39443200 | -1.57207000 |
| H | 3.06815200 | -0.46341700 | -2.10033900 |
| C | 4.06832700 | 0.07566800 | 1.19639500 |
| H | 3.33463400 | 0.87789800 | 1.25559600 |
| C | 3.68572100 | -1.18933700 | 1.40931100 |
| H | 4.43942300 | -1.96995000 | 1.30923400 |
| C | 2.32406100 | -1.66405400 | 1.70227300 |
| C | 1.35175100 | -0.82392900 | 2.12409700 |
| H | 1.61613000 | 0.21243400 | 2.32229000 |
| C | -0.05391000 | -1.13928200 | 2.26430300 |
| H | -0.37769200 | -2.15846900 | 2.07296200 |
| C | -0.99524900 | -0.20278100 | 2.49682700 |
| H | -0.69454000 | 0.83190900 | 2.65737600 |
| C | 1.43026500 | 0.26815300 | -1.04703500 |
| C | 0.76614600 | 1.38099000 | -0.27424700 |
| H | 0.42683400 | 2.18374100 | -0.93168800 |
| H | -0.09432700 | 1.02005000 | 0.28581400 |
| H | 1.46968500 | 1.81378800 | 0.44119400 |
| C | 0.76001100 | -1.05969400 | -1.24886400 |
| O | 1.38925200 | -2.03952400 | -1.62021400 |
| C | 7.13683900 | 1.96568500 | -0.22289400 |
| H | 7.29422000 | 2.85591900 | -0.83572100 |
| H | 7.60548800 | 2.13250600 | 0.75005900 |
| H | 7.65263000 | 1.12907900 | -0.70535200 |
| H | 5.15463500 | 2.53509100 | 0.35991000 |
| C | 2.11241600 | -3.13206900 | 1.44765700 |
| H | 2.90964000 | -3.70601400 | 1.92916300 |
| H | 1.15174400 | -3.49565200 | 1.80354700 |
| H | 2.15891800 | -3.32714100 | 0.37223600 |
| C | -2.42021300 | -0.47391400 | 2.43198700 |
| H | -2.71284200 | -1.51507200 | 2.29631300 |
| C | -3.37070300 | 0.47073900 | 2.46631600 |
| H | -3.06309700 | 1.50679900 | 2.60545300 |
| C | -4.83587200 | 0.20273600 | 2.30864200 |
| H | -5.40964600 | 0.62173300 | 3.14003000 |
| H | -5.22427600 | 0.66457300 | 1.39334200 |
| H | -5.03415700 | -0.86947700 | 2.25364200 |
| C | -0.69752300 | -1.20426600 | -1.01319800 |
| C | -1.70975000 | -0.34321000 | -1.26334300 |
| O | -1.62884400 | 0.86275400 | -1.79868500 |
| H | -2.44476700 | 1.37198500 | -1.59409300 |
| C | -1.30866900 | -2.46261000 | -0.50124700 |
| O | -0.75206700 | -3.47020800 | -0.09713400 |
| N | -2.66435600 | -2.27890300 | -0.51682600 |
| H | -3.28405600 | -2.90682300 | -0.02686300 |
| C | -3.05573700 | -0.95952800 | -0.94938300 |
| H | -3.53680500 | -0.39709400 | -0.14095200 |
| C | -3.98607900 | -0.96036400 | -2.15749800 |
| H | -4.89418100 | -1.51296000 | -1.89658200 |
| H | -4.26448900 | 0.07273100 | -2.38779500 |
| O | -3.30930300 | -1.56609200 | -3.24302800 |
| H | -3.89971100 | -1.58409800 | -4.00122600 |
| C | -3.73645300 | 2.89822100 | 0.05211000 |
| O | -3.86371000 | 2.03511400 | -0.79067800 |
| O | -2.51253200 | 3.35578200 | 0.31506400 |
| H | -2.52365400 | 4.04610500 | 0.99113800 |
| C | -4.87549400 | 3.49087700 | 0.82469900 |
| H | -4.73577200 | 3.30428700 | 1.89222900 |
| H | -4.89836600 | 4.57158800 | 0.67223400 |
| H | -5.81062100 | 3.05002200 | 0.49172100 |

------------------------------------------------------------

### SUPPLEMENTARY REFERENCES


1. Li, X., Zheng, Q., Yin, J., Liu, W. & Gao, S. Chemo-enzymatic synthesis of equisetin. *Chem Commun* **53**, 4695-4697 (2017).
2. Kato, N. et al. Control of the stereochemical course of [4+2] cycloaddition during *trans*-decalin formation by Fsa2-family enzymes. *Angew Chem Int Ed Engl* **57**, 9754-9758 (2018).
3. Sawyer, L., Brownlow, S., Polikarpov, I. & Wu, S.-Y. β-Lactoglobulin: structural studies, biological clues. *Int Dairy J* **8**, 65-72 (1998).
4. Tian, Z. et al. An enzymatic [4+2] cyclization cascade creates the pentacyclic core of pyrroindomycins. *Nat Chem Biol* **11**, 259-265 (2015).
5. Zheng, Q. et al. Enzyme-dependent [4 + 2] cycloaddition depends on lid-like interaction of the N-terminal sequence with the catalytic core in PyrI4. *Cell Chem Biol* **23**, 352-360 (2016).
6. Kim, K.W. et al. Trimeric structure of (+)-pinoresinol-forming dirigent protein at 1.95 A resolution with three isolated active sites. *J Biol Chem* **290**, 1308-1318 (2015).
7. Ester, M., Kriegel, H.-P., Sander, J. & Xu, X. A density-based algorithm for discovering clusters in large spatial databases with noise. in *Proceedings of the Second International Conference on Knowledge Discovery and Data Mining* 226–231 (AAAI Press, Portland, Oregon, 1996).
8. Wallace, A.C., Laskowski, R.A. & Thornton, J.M. LIGPLOT: a program to generate schematic diagrams of protein-ligand interactions. *Protein Eng* **8**, 127-134 (1995).
9. Chiu, H.J. et al. Structure of the first representative of Pfam family PF09410 (DUF2006) reveals a structural signature of the calycin superfamily that suggests a role in lipid metabolism. *Acta Crystallogr F* **66**, 1153-1159 (2010).
10. Hiseni, A., Arends, I.W. & Otten, L.G. Biochemical characterization of the carotenoid 1,2-hydratases (CrtC) from Rubrivivax gelatinosus and Thiocapsa roseopersicina. *Appl Microbiol Biotechnol* **91**, 1029-36 (2011).
11. Hiseni, A., Otten, L.G. & Arends, I. Identification of catalytically important residues of the carotenoid 1,2-hydratases from Rubrivivax gelatinosus and Thiocapsa roseopersicina. *Appl Microbiol Biotechnol* **100**, 1275-1284 (2016).
12. Kakule, T.B. et al. Native promoter strategy for high-yielding synthesis and engineering of fungal secondary metabolites. *ACS Synth Biol* **4**, 625-633 (2015).
13. Kato, N. et al. A new enzyme involved in the control of the stereochemistry in the decalin formation during equisetin biosynthesis. *Biochem Biophys Res Commun* **460**, 210-215 (2015).
14. Sato, M. et al. Involvement of lipocalin-like CghA in decalin-forming stereoselective

intramolecular [4+2] cycloaddition. *Chembiochem* **16**, 2294-2298 (2015).

15. Li, L. et al. Biochemical characterization of a eukaryotic decalin-forming Diels-Alderase. *J Am Chem Soc* **138**, 15837-15840 (2016).
16. Li, L. et al. Genome mining and assembly-line biosynthesis of the UCS1025A pyrrolizidinone family of fungal alkaloids. *J Am Chem Soc* **140**, 2067-2071 (2018).
17. Tan, D. et al. Genome-mined Diels-Alderase catalyzes formation of the *cis*-octahydrodecalins of varicidin A and B. *J Am Chem Soc* **141**, 769-773 (2019).
18. Smith, R.H.B., Dar, A.C. & Schlessinger, A. PyVOL: A PyMOL plugin for visualization, comparison, and volume calculation of drug-binding sites. *bioRxiv* (2019).
19. Schrodinger, LLC. The PyMOL Molecular Graphics System, Version 1.3r1. (2010).
20. Jung, J. et al. GENESIS: a hybrid-parallel and multi-scale molecular dynamics simulator with enhanced sampling algorithms for biomolecular and cellular simulations. *Wiley Interdiscip Rev Comput Mol Sci* **5**, 310-323 (2015).
21. Kobayashi, C. et al. GENESIS 1.1: A hybrid-parallel molecular dynamics simulator with enhanced sampling algorithms on multiple computational platforms. *J Comput Chem* **38**, 2193-2206 (2017).
22. Maier, J.A. et al. ff14SB: Improving the accuracy of protein side chain and backbone parameters from ff99SB. *J Chem Theory Comput* **11**, 3696-3713 (2015).
23. Jorgensen, W.L., Chandrasekhar, J., Madura, J.D., Impey, R.W. & Klein, M.L. Comparison of simple potential functions for simulating liquid water. *J Chem Phys* **79**, 926-935 (1983).
24. Wang, J., Wolf, R.M., Caldwell, J.W., Kollman, P.A. & Case, D.A. Development and testing of a general amber force field. *J Comput Chem* **25**, 1157-1174 (2004).
25. Case, D.A. et al. AMBER 2018. *University of California, San Francisco.* (2018).
26. Ryckaert, J.-P., Ciccotti, G. & Berendsen, H.J.C. Numerical integration of the cartesian equations of motion of a system with constraints: molecular dynamics of *n*-alkanes. *J Comput Phys* **23**, 327-341 (1977).
27. Miyamoto, S. & Kollman, P.A. Settle: An analytical version of the SHAKE and RATTLE algorithm for rigid water models. *J Comput Chem* **13**, 952-962 (1992).
28. Darden, T., York, D. & Pedersen, L. Particle mesh Ewald: AnN·log(N) method for Ewald sums in large systems. *J Chem Phys* **98**, 10089-10092 (1993).
29. Essmann, U. et al. A smooth particle mesh Ewald method. *J Chem Phys* **103**, 8577-8593 (1995).
30. Tuckerman, M., Berne, B.J. & Martyna, G.J. Reversible multiple time scale molecular dynamics. *J Chem Phys* **97**, 1990-2001 (1992).
31. Bussi, G., Donadio, D. & Parrinello, M. Canonical sampling through velocity rescaling. *J*

*Chem Phys* **126**, 014101 (2007).

32. Bussi, G., Zykova-Timan, T. & Parrinello, M. Isothermal-isobaric molecular dynamics using stochastic velocity rescaling. *J Chem Phys* **130**, 074101 (2009).
33. Webb, B. & Sali, A. Comparative protein structure modeling using MODELLER. *Curr Protoc Bioinformatics* **54**, 5.6.1-5.6.37 (2016).
34. Olsson, M.H., Sondergaard, C.R., Rostkowski, M. & Jensen, J.H. PROPKA3: Consistent treatment of internal and surface residues in empirical pKa predictions. *J Chem Theory Comput* **7**, 525-537 (2011).
35. Sondergaard, C.R., Olsson, M.H., Rostkowski, M. & Jensen, J.H. Improved treatment of ligands and coupling effects in empirical calculation and rationalization of pKa values. *J Chem Theory Comput* **7**, 2284-2295 (2011).
36. Sugita, Y. & Okamoto, Y. Replica-exchange molecular dynamics method for protein folding. *Chem Phy Lett* **314**, 141-151 (1999).
37. Kamiya, M. & Sugita, Y. Flexible selection of the solute region in replica exchange with solute tempering: Application to protein-folding simulations. *J Chem Phys* **149**, 072304 (2018).
38. Liu, P., Kim, B., Friesner, R.A. & Berne, B.J. Replica exchange with solute tempering: a method for sampling biological systems in explicit water. *Proc Natl Acad Sci U S A* **102**, 13749-13754 (2005).
39. Terakawa, T., Kameda, T. & Takada, S. On easy implementation of a variant of the replica exchange with solute tempering in GROMACS. *J Comput Chem* **32**, 1228-1234 (2011).
40. Wang, L., Friesner, R.A. & Berne, B.J. Replica exchange with solute scaling: a more efficient version of replica exchange with solute tempering (REST2). *J Phys Chem B* **115**, 9431-9438 (2011).
41. Niitsu, A., Re, S., Oshima, H., Kamiya, M. & Sugita, Y. De novo prediction of binders and nonbinders for T4 lysozyme by gREST simulations. *J Chem Inf Model* **59**, 3879-3888 (2019).
42. Oshima, H., Re, S. & Sugita, Y. Prediction of protein-ligand binding pose and affinity using the gREST+FEP method. *J Chem Inf Model* **60**, 5382-5394 (2020).
43. Pedregosa, F. et al. Scikit-learn: Machine learning in Python. *J Mach Learn Res* **12**, 2825-2830 (2011).
